# Dopa decarboxylase is a genetic hub of parental control over offspring behavior

**DOI:** 10.1101/2020.06.23.168195

**Authors:** Paul J. Bonthuis, Susan Steinwand, Wei-Chao Huang, Cornelia N. Stacher Hörndli, Jared Emery, Stephanie Kravitz, Elliott Ferris, Christopher Gregg

**Affiliations:** Department of Comparative Biosciences, University of Illinois at Urbana-Champaign College of Veterinary Medicine, Urbana, Illinois; Departments of Neurobiology & Anatomy, University of Utah School of Medicine, Salt Lake City, Utah; Human Genetics, University of Utah School of Medicine, Salt Lake City, Utah

**Keywords:** Genomic imprinting, parental effects, foraging, dopamine, serotonin, noradrenaline, dopa decarboxylase, behavior, machine learning, CRISPR knock-in mice, hypothalamic-pituitary-adrenal axis, HPA axis

## Abstract

Dopa decarboxylase (DDC) regulates the synthesis of monoaminergic neurotransmitters and is linked to psychiatric and metabolic disorders. *Ddc* exhibits complex genomic imprinting effects that have not been functionally studied. Here, we investigate different noncanonical imprinting effects at the cellular level with a focus on *Ddc*. Using allele-specific reporter mice, we found *Ddc* exhibits dominant expression of the maternal allele in subpopulations of cells in 14 of 52 brain regions, and dominant paternal or maternal allele expression in adrenal cell subpopulations. Maternal versus paternal *Ddc* allele null mutations differentially affect offspring social, foraging and exploratory behaviors. Machine learning analyses of naturalistic foraging in *Ddc*^−/+^ and ^+/−^ offspring uncovered finite behavioral sequences controlled by the maternal versus paternal *Ddc* alleles. Additionally, parental *Ddc* genotype is revealed to affect behavior independent of offspring genotype. Thus, *Ddc* is a hub of maternal and paternal influence on behavior that mediates diverse imprinting and parental effects.

**HIGHLIGHTS:**

- Dopa decarboxylase (Ddc) allelic expression resolved at the cellular level
- Cells differentially express maternal versus paternal *Ddc* alleles
- Maternal and paternal *Ddc* alleles control distinct behavioral sequences
- Parental *Ddc* genotype affects offspring independent of mutation transmission

**eTOC:** Allelic reporter mice and machine learning analyses reveal dopa decarboxylase is affected by diverse imprinting and parental effects that shape finite behavioral sequences in sons and daughters.

## INTRODUCTION

Gene regulatory mechanisms are central players in phenotype evolution (Carroll, 2008; Wray, 2007). However, we do not fully understand gene regulatory mechanisms in mammalian cells or how they shape different phenotypes, including brain functions and behaviors. Recent studies suggest important gene regulatory mechanisms remain to be uncovered at the allele level in the mammalian genome (Kravitz and Gregg, 2019). Genomic imprinting is a form of allele-specific gene regulation that evolved in mammals and flowering plants (Haig, 2000a). It involves heritable epigenetic mechanisms that cause preferential expression of the maternal or paternal allele for some genes in offspring and has important roles in the brain (Gregg, 2014; Kravitz and Gregg, 2019; Perez et al., 2016). Imprinting effects impact the phenotypic effects of a heterozygous mutation according to whether it resides in the maternal or paternal allele (Peters, 2014). The function of imprinting is debated (Haig and Haig, 2004; Spencer and Clark, 2014). A deeper understanding of imprinting could improve our understanding of maternal and paternal influences on mammalian phenotypes.

Canonical imprinting involves epigenetic silencing of one parental allele in offspring. However, using RNASeq profiling, we (Bonthuis et al., 2015), and others (Andergassen et al., 2017; Babak et al., 2015; Crowley et al., 2015; Perez et al., 2015), described imprinted genes that exhibit a bias to express either the maternal or paternal allele at a higher level. We refer to these cases as “noncanonical imprinting” effects (Bonthuis et al., 2015). They are more frequent in the mouse genome than canonical imprinting, exist in wild-derived populations and are enriched in the brain (Bonthuis et al., 2015). Others described this phenomenon as a “*parental allele bias*” or “*nonclassical imprinting*” (Andergassen et al., 2017; Babak et al., 2015; Crowley et al., 2015; Perez et al., 2015). However, we observed that these effects appear to be more than an allele expression *bias*, and may be a cell-specific form of imprinting (Bonthuis et al., 2015). Additionally, we found noncanonical imprinting can interact with inherited heterozygous mutations to affect behavior (Bonthuis et al., 2015). The cell populations impacted by noncanonical imprinting are largely unknown. The impacts of noncanonical imprinting on protein levels and effects on offspring behavior are also unknown.

Here, we further investigate noncanonical imprinting effects at the cellular level in mice and test different roles in behavior. We focus on *dopa decarboxylase* (*Ddc*), an enzyme required for synthesis of monoaminergic neurotransmitters, including dopamine (DA), norepinephrine (NE), epinephrine (E) and serotonin (5-HT). *Ddc* is located in the genome next to the canonical imprinted gene, *Grb10. Grb10* exhibits dominant paternal allele expression in the brain and dominant maternal allele expression in non-neural tissues (Menheniott et al., 2008a; Plasschaert and Bartolomei, 2015). We previously found *Ddc* noncanonical imprinting effects that involve biased expression of the *<u>maternal</u>* allele in the mouse brain (Bonthuis et al., 2015). The effects are strongest in the hypothalamus. *Ddc* imprinting was first uncovered in mice with uniparental duplications and found to exhibit dominant expression of the *<u>paternal</u>* allele in the embryonic heart (Menheniott et al., 2008a). The cellular nature and function of maternal and paternal *Ddc* imprinting effects are not known.

The monoamine system is fundamental. It modulates brain functions, including reward, feeding, motivation, activity, fear, arousal, sensorimotor processes, social behaviors, learning & memory and others (Gershman and Uchida, 2019; Klein et al., 2019; Okaty et al., 2019; Sara, 2009; Volkow et al., 2017). Monoamines have important roles in psychiatric and neurological disorders, including addiction, depression, sleep disorders, obesity, Parkinson’s disease and others (Klein et al., 2019; Marazziti, 2017; Sara, 2009). Monoamine signaling also has roles in non-neural tissues (Arreola et al., 2016; Martin et al., 2017). In the adrenal medulla, subpopulations of DA, NE and E expressing cells modulate stress responses as part of the Hypothalamic-Pituitary-Adrenal axis (Kvetnansky et al., 2009). Additionally, 5-HT and DA regulate gene expression through histone modifications involving serotonylation (Farrelly et al., 2019) and dopaminylation (Lepack et al., 2020), and influence embryonic development (Bérard et al., 2019). Previous studies investigating the functional diversity of monoaminergic cells in the brain focused on biophysical properties, transmitter co-release, connectivity, developmental lineage and molecular profiling (Berke, 2018; Farassat et al., 2019; Gershman and Uchida, 2019; Okaty et al., 2019). Currently, we have little understanding of how allele-specific gene regulatory mechanisms contribute to the functional diversification of monoaminergic cells or how *Ddc* imprinting effects may shape the effects of inherited genetic mutations in the *Ddc* locus. In humans, *DDC* genetic variation is significantly linked to multiple biomedically important phenotypes (Watanabe et al., 2019).

Our study further tests the hypothesis that noncanonical imprinting is a cell-type dependent form of imprinting that shapes ethological behavior. We test whether different noncanonical imprinting effects are linked to neuronal versus non-neuronal brain cell-types. We then perform a high-resolution cellular analysis of different neural and non-neural *Ddc* imprinting effects using targeted knock-in allele-specific reporter mice. To investigate potentially different functional roles for the maternal versus paternal *Ddc* alleles in shaping behavior, we use reciprocal *Ddc* heterozygous mice and test how the parental origin of the mutated allele affects social, foraging and exploratory behaviors. Our approach uses behavioral and machine-learning methods we previously developed to analyze naturalistic foraging in mice (Stacher Hörndli et al., 2019). Our previous study found that foraging patterns are constructed from finite, genetically-controlled behavioral sequences that we call modules. Here, we discover that multiple forms of parental effects are transmitted through the *Ddc* locus and affect behavior. Our results uncover distinct functional effects of the maternal versus paternal *Ddc* alleles, linking each allele to aspects of social behavior and to finite foraging modules in sons and daughters. Additionally, parental *Ddc* genotype is revealed to cause transgenerational effects on behavior that are independent of the offspring’s genotype. Our findings suggest *Ddc* evolved as a genetic hub mediating different parental influences on offspring behavior.

## RESULTS

### Defining imprinted gene expression and imprinting effects in major hypothalamic cell classes

For most noncanonical imprinted genes, we do not know the identity of the brain cell-types that express each gene or exhibit imprinting effects. To investigate the expression of maternally expressed genes (MEGs) and paternally expressed genes (PEGs) in different hypothalamic cell types we analyzed public single-cell RNASeq data from adult mouse hypothalamus (Chen et al., 2017). We calculated the mean expression level for each imprinted gene in previously reported hypothalamic cell types, including 11 non-neuronal and 34 neuronal cell types (Chen et al., 2017). Unsupervised clustering of the results grouped MEGs and PEGs with similar expression patterns across different hypothalamic cell types (**Fig. S1**). The data show that expression of most imprinted genes is detected in both neuronal and non-neuronal cell populations. However, it is not known whether the imprinting effects occur equally in both cell classes.

To identify brain cell types exhibiting imprinting for canonical and non-canonical imprinted genes, we initially attempted single-cell RNA-Seq (scRNA-Seq) profiling of brain cells purified from adult F1 hybrid mice generated from crossing CastEiJ x C57Bl/6J mice to resolve allele-specific expression and imprinting at the cellular level (not shown). However, we found that ∼99% of the >5000 cells profiled are non-neuronal. The cellular bias and known technical noise of the data limited our ability to draw conclusions about allelic expression in subpopulations of cells. Therefore, we developed a different strategy based on studies of gene co-expression networks that found molecular subtypes of brain cells from bulk RNASeq replicates (Kelley et al., 2018; Parikshak et al., 2013; Willsey et al., 2013). These previous methods uncovered co-expression networks by identifying genes with correlated expression patterns across biological replicates. Expanding on this idea, our approach tests whether the magnitude of the *<u>imprinting effect</u>* for a given imprinted gene correlates with the expression level of genes that are known markers of brain cell types (**Fig. 1A,B**). Thus, for genes with cell-type specific imprinting, our approach expects the magnitude of the imprinting effect (ie. maternal – paternal allele expression difference) to be positively correlated to the expression of marker genes for the imprinted cell-types, and not correlated to markers in cells lacking the imprinting effect (**Fig. 1A,B**).

**Figure 1.**
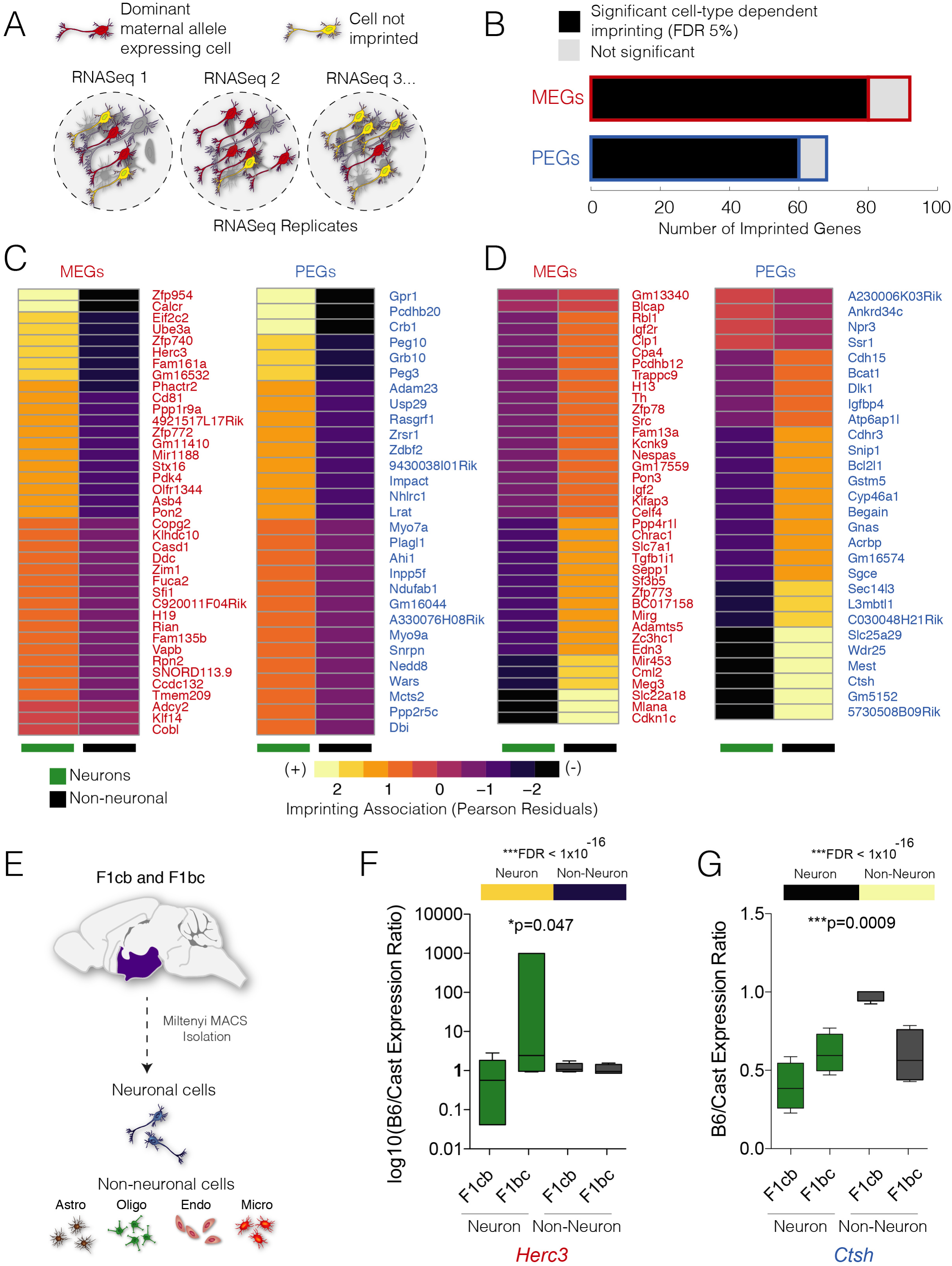
**Imprinting∼Expression correlation networks (IENs) identify major cell classes exhibiting imprinting for canonical and non-canonical imprinted genes.** (A) A schematic depicting the underlying assumptions of IEN analysis that variance in bulk RNASeq detected imprinting magnitude is a function of the prevalence of the imprinted cell population in the sample. As a consequence, the magnitude of the imprinting effect for a gene is expected to correlate with the expression of molecular markers expressed in the imprinted cell-type. (B) The number of MEGs and PEGs with imprinting effects exhibiting statistically significant dependence on the expression of brain cell-type specific marker genes (FDR <5%). (C and D) Canonical and non-canonical MEGs and PEGs with tissue-level imprinting effects predominantly seen in neurons or non-neuronal cells in the hypothalamus. The shown MEGs and PEGs have a significant IEN Chi-Square test result (FDR 5%). The heatmap shows the Pearson residuals that indicate whether the imprinting effect is positively (yellow) or negatively (purple) associated with neurons (left column) or non-neuronal (right column) brain cells. MEGs and PEGs with predominant imprinting in neurons (C) or non-neuronal (D) cells are shown. (E-G) Pyrosequencing validates IEN defined cell-type imprinting effects for noncanonical imprinted genes in the adult mouse hypothalamus. The schematic shows a summary of the experimental workflow to isolate neurons and non-neuronal cells from F1cb and F1bc hypothalamus for pyrosequencing analysis of imprinting (E). Pyrosequencing validation of non-canonical imprinting effects in isolated neuronal versus non-neuronal cells for *Herc3* (F, MEG) and *Ctsh* (G, PEG). IEN predicts preferential expression of the maternal allele in neurons for *Herc3* and the paternal allele in non-neuronal cells for *Ctsh* (heatmap on top, and see Fig. S1). Pyrosequencing confirmed these predictions, found a significant interaction between imprinting effect and cell class (two-way anova p-value shown) and uncovered a maternal allele bias in neurons only for *Herc3* and a paternal allele bias in non-neuronal cells only for *Ctsh*.

To compute *<u>Imprinting ∼ Cell-type Marker Expression</u>* correlation networks, we used our published bulk RNASeq replicates for the hypothalamus (arcuate nucleus region) from adult female F1cb (CastEiJ x C57BL/6J, n=9) and F1bc (C57BL/6J x CastEiJ, n=9) hybrid mice (Bonthuis et al., 2015). We calculated the allele expression difference (maternal – paternal) for each imprinted gene and replicate. We then computed the expression levels of mouse-human conserved markers of neurons, oligodendrocytes, astrocytes, endothelial cells and microglia (McKenzie et al., 2018). An imprinting ∼ marker expression correlation matrix (Spearman Rank) was then calculated to relate the variance in imprinting magnitude (maternal - paternal allele expression difference) to the variance in the expression of cell-type specific marker genes across RNASeq replicates. Internal control studies and methodological details are provided in the supplemental data (**Data S1**) and methods. We performed the analysis for all autosomal imprinted genes in the mouse (Bonthuis et al., 2015) to identify MEGs and PEGs with imprinting effects significantly linked to the expression of neuronal versus non-neuronal cell marker genes (False Discovery Rate (FDR) < 5%). We found 87% of MEGs and 88% of PEGs have significant cell type dependence for imprinting (**Fig. 1B**) and defined the major cell classes linked to the imprinting effects for each gene (**Fig. 1C,D**). Our results define major cell classes driving tissue level detected imprinting effects.

To validate predicted imprinting effects for noncanonical imprinted genes and test whether the imprinting effect manifests as an allele-*bias* or as allele-*silencing* effect, we purified neuronal and nonneuronal cells from the adult female hypothalamus for F1cb and F1bc mice (**Fig. 1E**, STAR methods). We used targeted pyrosequencing of cDNA prepared from the purified cells to evaluate imprinting effects in each cell type for eight noncanonical imprinted genes. *Herc3* is a noncanonical MEG and our correlation analysis indicates imprinting in neurons, but not non-neuronal cells (**Fig. 1C**). Pyrosequencing validated this result and revealed a significant interaction between cross and cell-type, indicating cell-type dependent imprinting with a maternal allele expression bias in neurons (**Fig. 1F**). *Ctsh* is a noncanonical PEG predicted to exhibit imprinting and preferential paternal allele expression in non-neuronal cells (**Fig. 1D**). Pyrosequencing confirmed this prediction and a paternal expression bias in non-neuronal cells only (**Fig. 1G**). Pyrosequencing validated the predicted cell-type imprinting effects for seven of the eight noncanonical imprinted genes tested (p<0.05; cross X cell-type interaction). All cases showed an allele-*bias* rather than complete allele-*silencing*. Therefore, noncanonical imprinting effects continue to manifest as an allelic bias at the level of neuronal versus non-neuronal cells, leaving open the question of whether more pronounced allelic effects occur in more refined cellular subpopulations.

### Noncanonical imprinting causes dominant expression of the maternal *Ddc* allele in discrete subpopulations of brain cells

We next tested the hypothesis that noncanonical imprinting causes dominant expression of one parental allele in small subpopulations of cells and manifests at the protein level. We focused on *Ddc* and generated *Ddc* allele-specific reporter mice in which reporter constructs are knocked-in before the *Ddc* stop codon, placing a c-terminal V5 epitope tag onto the DDC protein for direct detection of one allele and a stable, nuclear eGFP reporter separated from DDC by a P2A cleavage site on the other allele **(Fig. 2A)**. Our *Ddc^eGFP^* line produced robust nuclear eGFP expression in cells for all major monoaminergic nuclei **(Fig. S2A)**. Similarly, the DDC protein is detected in *Ddc^V5^* targeted knock-in mice using immunolabeling with an anti-V5 antibody **(Fig. S2A)**. The two reporter lines were crossed in reciprocal matings to generate compound *Ddc^eGFP^*^/*V5*^ and *Ddc^V5/eGFP^* allelic reporter mice that reveal Ddc expression from the V5 tagged allele and expression of the other allele from nuclear eGFP (**Fig. S2B)**. Since eGFP is stable with a half-life greater than 26 hours (Corish and Tyler-Smith, 1999), we expect any V5+ and eGFP-brain cells to have stable silencing of the eGFP allele for at least days, enabling us to infer persistent allele-specific expression in affected cells.

**Figure 2.**
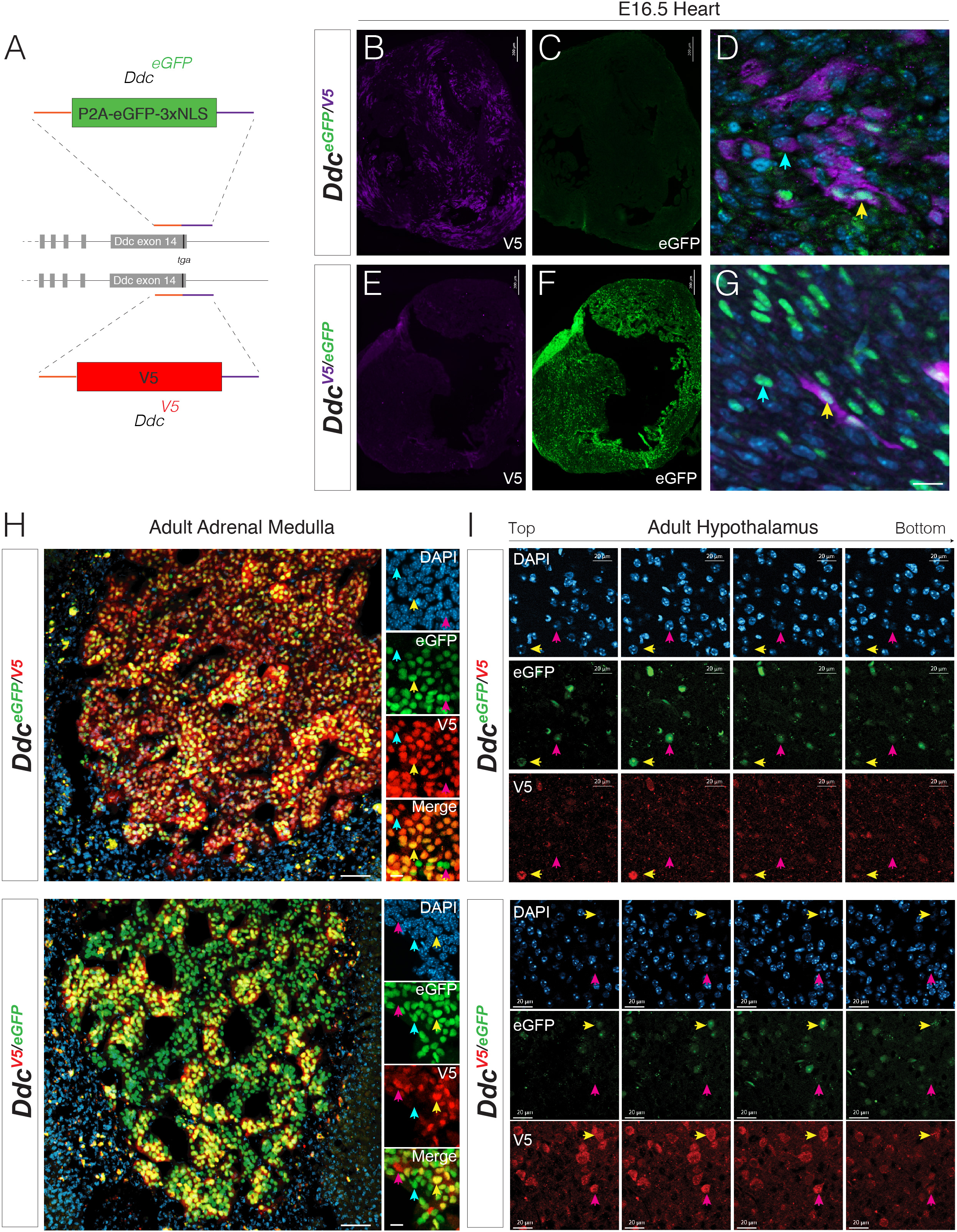
**Allelic reporter mice reveal cell-type dependent differential expression of the paternal and maternal *Ddc* alleles.** (A) Schematic summary of *Ddc* reporter mouse design for eGFP reporter allele and V5 tagged DDC protein allele. (B-G) *Ddc* exhibits dominant expression of the paternal allele in the developing heart. Compound *Ddc^eGFP/V5^* (B-D) and *Ddc^V5/eGFP^* (E-G) allelic reporter mice reveal dominant expression of the paternal *Ddc* allele in subpopulations of cardiac cells in the developing embryonic day (E)16.5 heart. Most DDC+ cells exhibit preferential expression of the paternal (B and F) over the maternal (C and E) allele. However, at higher magnification (D and G), subpopulations of DDC+ myocardial cells that express both parental alleles (yellow arrows) are revealed in addition to those that exhibit dominant expression of the paternal allele (white arrows). Size bar in G: 10 microns. (H) Dominant expression of the paternal and maternal *Ddc* allele in the adult adrenal medulla revealed from *Ddc^eGFP/V5^* and *Ddc^V5/eGFP^* mice (Size bar: 100 microns). Insets show high magnification images of cells in the adrenal medulla (Size bar: 20 microns). Yellow arrows indicate examples of cells co-expressing both parental alleles; blue arrows indicate examples of dominant paternal allele expression; pink arrows indicate examples of dominant maternal allele expression. (I) Subpopulations of DDC+ hypothalamic cells exhibit dominant expression of the maternal allele. Optical sections of images from *Ddc^eGFP/V5^* and *Ddc^V5/eGFP^* adult female hypothalamus reveal subpopulations of neurons expressing both *Ddc* alleles equally (yellow arrows) and subpopulations of neurons with dominant expression of the maternal *Ddc* allele (pink arrows). Z-stack of images shown. Size bars: 20 microns.

We first measured allelic expression in the embryonic day (E)16-17 heart. We found preferential expression of the paternal allele and silencing of the maternal allele in the *Ddc^eGFP^*^/*V5*^ (**Fig. 2B-D**) and *Ddc^V5/eGFP^* (**Fig. 2E-G**) reporter mice, confirming a previous report (Menheniott et al., 2008a). However, we also found heart cells that express both *Ddc* alleles (**Fig. 2D and G**, yellow arrows), in contrast to others exhibiting dominant paternal allele expression (**Fig. 2D and G**, white arrows). Our mice therefore resolved known *Ddc* imprinting effects and uncovered previously unknown allelic effects that could only be resolved with a cellular level analysis. We next began a detailed study of noncanonical *Ddc* imprinting effects in adults.

At the RNA and tissue levels in adults, *Ddc* noncanonical imprinting with a significant maternal allele bias in brain and liver, and a paternal allele bias in adrenal glands (Babak et al., 2015; Bonthuis et al., 2015). We began by investigating *Ddc* imprinting at the cellular and protein levels in the adrenal medulla, which contains DA, NE and E expressing cells that modulate stress responses (Kvetnansky et al., 2009). Overall, we found most cells co-express both alleles, but a general paternal allelic effect is apparent at low magnification (**Fig. 2H**). At higher magnification, discrete subpopulations of adrenal cells with different allelic expression effects are revealed (**Fig. 2H**, insets). We found subsets of DDC+ cells that exhibit dominant paternal allele expression (**Fig. 2H**, blue arrows) and a less frequent subpopulation with dominant maternal allele expression (**Fig. 2H**, pink arrows). These allelic subpopulations of adrenal cells were observed in both the *Ddc^eGFP^*^/*V5*^ and *Ddc^V5/eGFP^* reciprocal reporter lines (**Fig. 2H**). Other DDC+ adrenal cells exhibited biallelic expression (**Fig. 2H,** yellow arrows).

Next, we analyzed the brain. The strongest *Ddc* imprinting effects in brain are observed in the hypothalamus (Bonthuis et al., 2015). In *Ddc^eGFP/V5^* mice, optical sectioning of the hypothalamus revealed DDC+ cells exhibiting dominant *maternal* allele expression (**Fig. 2I,** pink arrows) and DDC+ cells expressing both alleles similarly (biallelic cells) (**Fig. 2I,** yellow arrows). To test whether these cellular subpopulations are also detectable when the parental origin of the allelic reporters is switched, we performed the same analysis in reciprocal *Ddc^V5/eGFP^* mice (**Fig. 2I**). We confirmed small populations of cells exhibiting dominant maternal *Ddc* allele expression. We did not observe cells with dominant paternal allele expression in both crosses. Therefore, noncanonical *Ddc* imprinting is more complex at the cellular level.

Many brain regions harbor monoaminergic neurons; we do not know the extent of DDC imprinting throughout the brain and whether some regions have imprinted cells dominantly expressing the maternal allele, while others have imprinted cells dominantly expressing the paternal allele. To answer this question, we tested 52 different brain regions in adult female mice for the presence versus absence of cells with dominant maternal or paternal DDC allele expression. We imaged coronal sections from the entire rostral-caudal axis of *Ddc^eGFP/V5^* and *Ddc^V5/eGFP^* mouse brains. Every fluorescence image was compared to an adjacent, parallel Nissl-stained section to define the anatomical location of the brain region(s) according to the Allen Brain Reference Atlas. We captured 783 images of DDC+ cell populations from *Ddc^eGFP/V5^* and *Ddc^V5/eGFP^* mice to score whether 1) each image contained imprinted neurons (**Fig. 3A**, AVPV, pink arrows) or not (**Fig. 3B**, VTA, yellow arrows), and 2) which reporter allele was dominant (eGFP vs. V5). All scoring was performed blinded to brain region and reporter cross (*Ddc^eGFP/V5^* or *Ddc^V5/eGFP^*).

**Figure 3.**
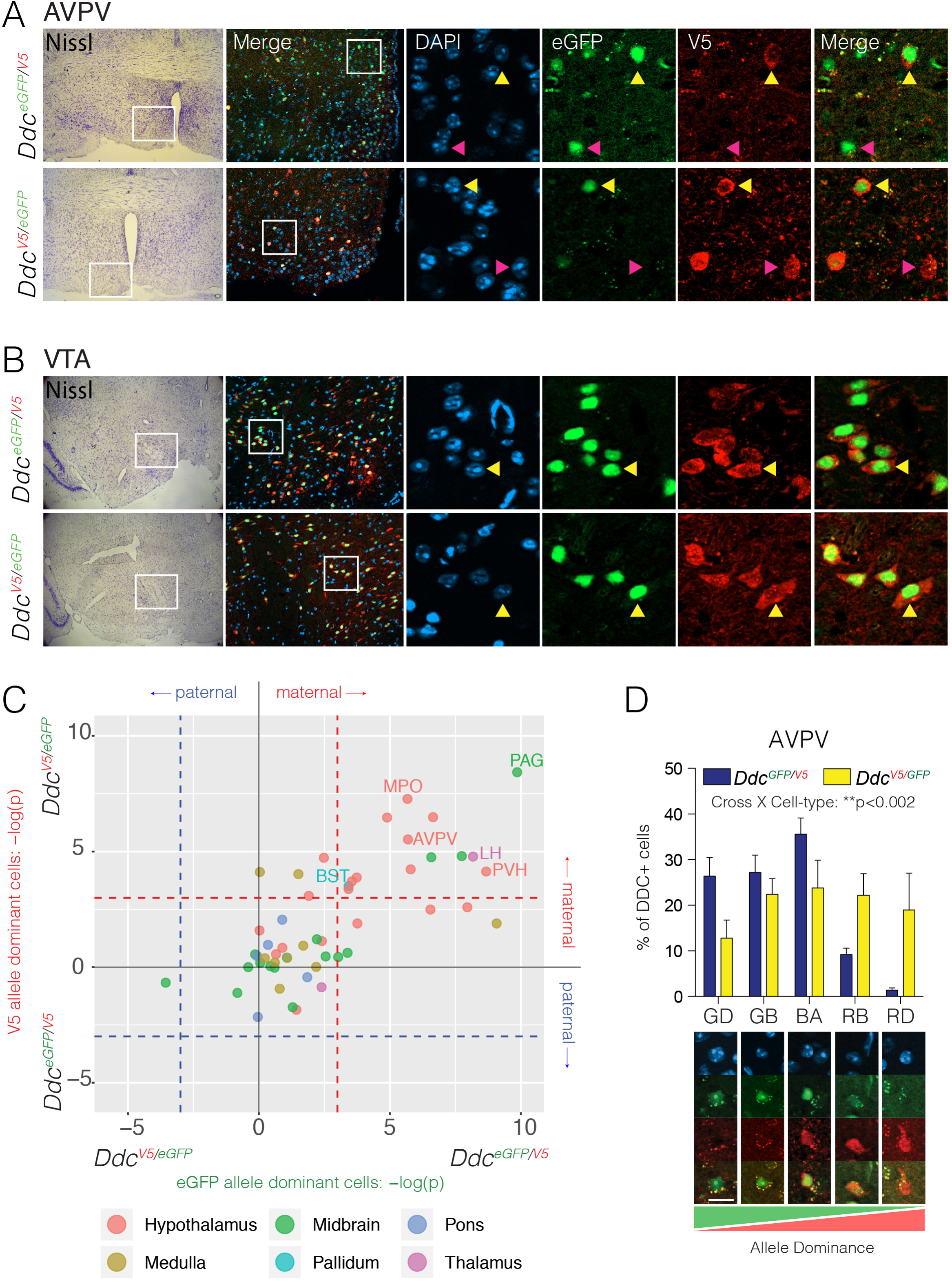
**Identification of 14 adult mouse brain regions containing DDC+ neurons with dominant maternal allele expression.** (A and B) The presence of DDC+ neurons with dominant maternal allele expression is brain region dependent. Maternal allele dominant neurons are observed in the AVPV of the hypothalamus (pink arrow) along with neurons expressing both alleles (yellow arrow) (A). However, in the VTA, only biallelic neurons expressing both alleles are observed (B). The white box in the Nissl labeled section shows the location of the analyzed brain region. The white box in the merged low magnification fluorescence image shows the location of the high magnification examples. (C) The scatterplot shows the identity of brain regions containing DDC+ neurons with dominant maternal allele expression confirmed in reciprocal *Ddc^eGFP/V5^* and *Ddc^V5/eGFP^* crosses and those that do not. Most impacted regions are in the hypothalamus (see legend below). For each brain region, multiple images were captured and blindly scored as having cells with dominant eGFP or V5 allele expressing cells. A Fisher’s test of a contingency table of the scored images for each region and cross was performed to determine whether a significant number of images had maternal or paternal allele dominant cells or neither (see Methods). Brain regions with significant maternal dominance are shown (dashed lines show p<0.05 threshold). Regions with cells having paternal allele dominance in both crosses were not observed. The data show 52 brain regions analyzed by 5 or more optical stacks per region per mouse (n=2; analyzed at 20X magnification). AVPV, anteroventral periventricular nucleus; BST, basal nucleus stria terminalis; LH, Lateral hypothalamic nucleus; MPO, medial preoptic area; PAG, periaqueductal grey; PVH, periventricular nucleus of the hypothalamus. See full atlas in Data S2. (D) Quantification of allele dominance effects at the cellular level in the AVPV in reciprocal *Ddc^eGFP/V5^* and *Ddc^V5/eGFP^* mice. The bar chart shows the percentage of DDC+ cells that are green eGFP+ dominant (GD), green eGFP+ biased (GB), equal biallelic (BA), red V5+ biased (RB) and red V5+ dominant (RD). Examples of each cell classes are shown below. Two-way ANOVA found a significant interaction between cross and allelic cell-type (p<0.002) and the data show the interaction is driven by maternal allele expression dominance (n=5).

To test whether a brain region is significantly enriched for dominant maternal versus paternal allele expressing cells, the number of images containing eGFP allele dominant cells was compared between the two crosses (*Ddc^eGFP/V5^* and *Ddc^V5/eGFP^*) using a Fisher’s Exact Test. The same was done for images containing V5 allele dominant cells (**Fig. 3C**). The results for each brain region and cross are plotted in **Fig. 3C** (colored by major brain divisions). Regions scored with dominant maternal allele expressing cells are plotted with a positive p-value and regions scored with dominant paternal allele expressing cells are plotted with a negative p-value. We uncovered 14 brain regions significantly enriched for cells with dominant expression of the maternal allele (**Fig. 3C**, upper-right quadrant). 38 brain regions were not enriched for imprinted DDC+ neurons (**Fig. 3C**), revealing brain region specificity for the imprinting that supports and extends our previous findings (Bonthuis et al., 2015). No regions exhibited dominant paternal allele expressing cells. Of the 14 brain regions enriched for neurons with dominant maternal Ddc allele expression, nine are located in the hypothalamus (9 of 20 tested; 45%), one is in the pallidum (1 of 2 tested), one in the thalamus (1 of 2 tested) and three reside in the midbrain (3 of 12; 20%). We did not find impacted hindbrain regions (0 of 5 regions in the pons, and 0 of 9 regions in the medulla; **Fig. 3C and see full atlas in Data S2 and brain region definitions in Table S1**).

To further test the findings from our atlas, we performed an independent and quantitative assessment of allele dominant DDC+ neurons in the anterior ventral periventricular area (AVPV), a top region enriched for maternal allele dominant DDC+ cells (**Fig. 3C**). Images of the AVPV were taken from *Ddc^eGFP/V5^* and *Ddc^V5/eGFP^* mice (n=5 mice per cross), and an investigator blind to the reporter cross (and subject) of each image scored every eGFP+ and/or V5+ cell in the image into one of five allelic categories: eGFP dominant, eGFP biased, biallelic, V5 biased or V5 dominant (**Fig. 3D**). We found a significant interaction between transgenic cross and cell category (**Fig. 3D**, p=0.002, two-way ANOVA), indicating a parent-of-origin effect on the proportion of cells in each allelic category. This result indicates that the relative proportion of eGFP versus Ddc-V5 allele dominant cells depends on the parental origin of the allele. A relative distribution shift toward eGFP+ dominant/biased cells occurs in the *Ddc^eGFP/V5^* reporter cross and a reciprocal distribution shift in the Ddc-V5 monoallelic/biased cells occurs in the *Ddc^V5/eGFP^* cross, indicating dominant maternal allele expression (**Fig. 3D**). We did observe some eGFP+ monoallelic cells in the *Ddc^V5/eGFP^* cross, indicating dominant expression of the paternal allele. However, these cells are rare in the reciprocal *Ddc^eGFP/V5^* cross (∼ 3% of cells), indicating they only arise in one cross and likely result from persistent nuclear eGFP expression from an earlier developmental time point. Overall, *Ddc* imprinting in adults provides maternal and paternal influences over the monoamine system at the cellular level in the brain and adrenal medulla, respectively.

### Dominant expression of the maternal *Ddc* allele in the hypothalamus is linked to subpopulations of GABAergic neurons

Monoaminergic neurons are anatomically, molecularly and functionally diverse. Some are glutamatergic, while some are GABAergic (Trudeau and Mestikawy, 2018). This dictates excitatory versus inhibitory neurotransmission. To explore the nature and function of imprinted DDC+ neurons, we tested whether dominant maternal allele expression is significantly linked to glutamatergic or GABAergic neuronal identity in the mouse hypothalamus. We applied the imprinting ∼ marker expression correlation approach introduced above, using marker genes previously found for 18 different GABAergic and 15 different glutamatergic neuron types in the mouse hypothalamus (Chen et al., 2017). We first aggregated the marker genes into two collections, all GABA neurons versus all glutamatergic neurons, and created a *Ddc* imprinting ∼ marker expression correlation matrix. The relative expression of the maternal versus paternal Ddc alleles depends significantly on GABA versus glutamate neuron marker gene expression (P = 0.0051, chi-square test) (**Fig. S3A**). Furthermore, maternal *Ddc* allele expression is positively associated with GABAergic neuron markers, indicating preferential maternal allele expression in GABA neurons (**Fig. S3A**). Immunohistochemical triple labeling of GABA, eGFP and Ddc-V5 in Ddc*^V5/eGFP^* transgenic mice confirmed that hypothalamic cells exhibiting dominant expression of the maternal *Ddc* allele co-express GABA (**Fig. S3B**, white arrows and zoomed images). Expression of GABA is not unique to the imprinted DDC+ neurons, since GABA co-expression was also observed in biallelic DDC+ cells (**Fig. S3B**, yellow arrows).

We next tested whether expression of the maternal versus paternal *Ddc* allele depends significantly on the specific subtypes of GABAergic or glutamatergic neurons. We constructed *Ddc* imprinting ∼ expression correlation matrices using the marker genes for each subtype of neuron (Chen et al., 2017). We found significant dependence on GABAergic and glutamatergic neuronal subtype (**Fig. S3C**, P=0.014, chi-square test). The majority (71%) of neuron types positively associated with maternal *Ddc* allele expression are GABAergic (**Fig. S3C**), of which GABA13 and GABA12 are GABAergic neurons are most strongly linked to maternal *Ddc* allele expression. We found Glu13, Glu6, Glu7, Glu15, Glu1 and GABA10 neurons are negatively associated with maternal *Ddc* allele expression (**Fig. S3C**). Thus, our results link *Ddc* imprinting effects to subtypes of GABAergic hypothalamic neurons.

### *Ddc* imprinting shapes offspring social behaviors in a sex dependent manner

We do not know the functions of the *Ddc* imprinting effects described above. Our data suggest that *Ddc* imprinting may shape maternal and paternal influences on the Hypothalamic-Pituitary-Adrenal axis, and other neural systems. Thus, *Ddc* imprinting could affect behavior; thereby impacting the behavioral effects of inherited heterozygous mutations in *Ddc* depending upon the parental origin of the mutated allele. Given our finding that some brain cell populations exhibit dominant expression of the maternal *Ddc* allele, our <u>primary hypothesis</u> is that loss of maternal *Ddc* allele function in offspring significantly affects offspring behavior and the effects differ from loss of paternal allele function. Alternative, haploinsufficiency and loss of the maternal *Ddc* allele might have no effect, or loss of the maternal versus paternal allele may cause the same phenotype in the offspring regardless of the parent-of-origin. Imprinted genes are proposed to affect social behaviors (Isles et al., 2006; Ubeda and Gardner, 2010) and we therefore began by testing the effects of *Ddc* imprinting on sociability and social novelty seeking behaviors.

We generated heterozygous *Ddc* mutant mice using a reciprocal cross strategy (**Fig. 4A,B**). In the first cross, a heterozygous mother and wildtype father generated offspring that are either wildtype (*Ddc^+/+^*) or have a mutated maternal allele (*Ddc^−/+^*) (**Fig. 4A**). In the reciprocal cross, the father is heterozygous and the mother is wildtype, generating wildtype (*Ddc^+/+^*) or mutated paternal allele (*Ddc^+/−^*) offspring (**Fig. 4B**). This design allows us to contrast the effects of maternal and paternal allele deletion. We tested sociability and social novelty seeking in these offspring using the three-chamber social interaction test (Moy et al., 2004). To evaluate sociability, we compared the time spent in the chamber with the jail containing a male conspecific (Conspecific) to time in the chamber with an empty jail (**Fig. 4C**). Male *Ddc^−/+^* mice with a maternal mutant allele spend proportionately less cumulative time in the chamber with the male conspecific relative to the empty chamber compared to their *Ddc^+/+^* littermates (**Fig. 4D**, mat) (p=0.042; genotype X chamber interaction, mixed linear model). In contrast, significant effects were not observed in *Ddc^+/−^* males with a paternal mutant allele compared to littermate controls (**Fig. 4D**, pat). Other measures, including the cumulative duration spent near the conspecific jail compared to the empty jail, support this result (**Fig. 4D**). We did not observe significant effects in females. Thus, loss of the maternal *Ddc* allele significantly reduces sociability with male conspecifics in sons.

**Figure 4.**
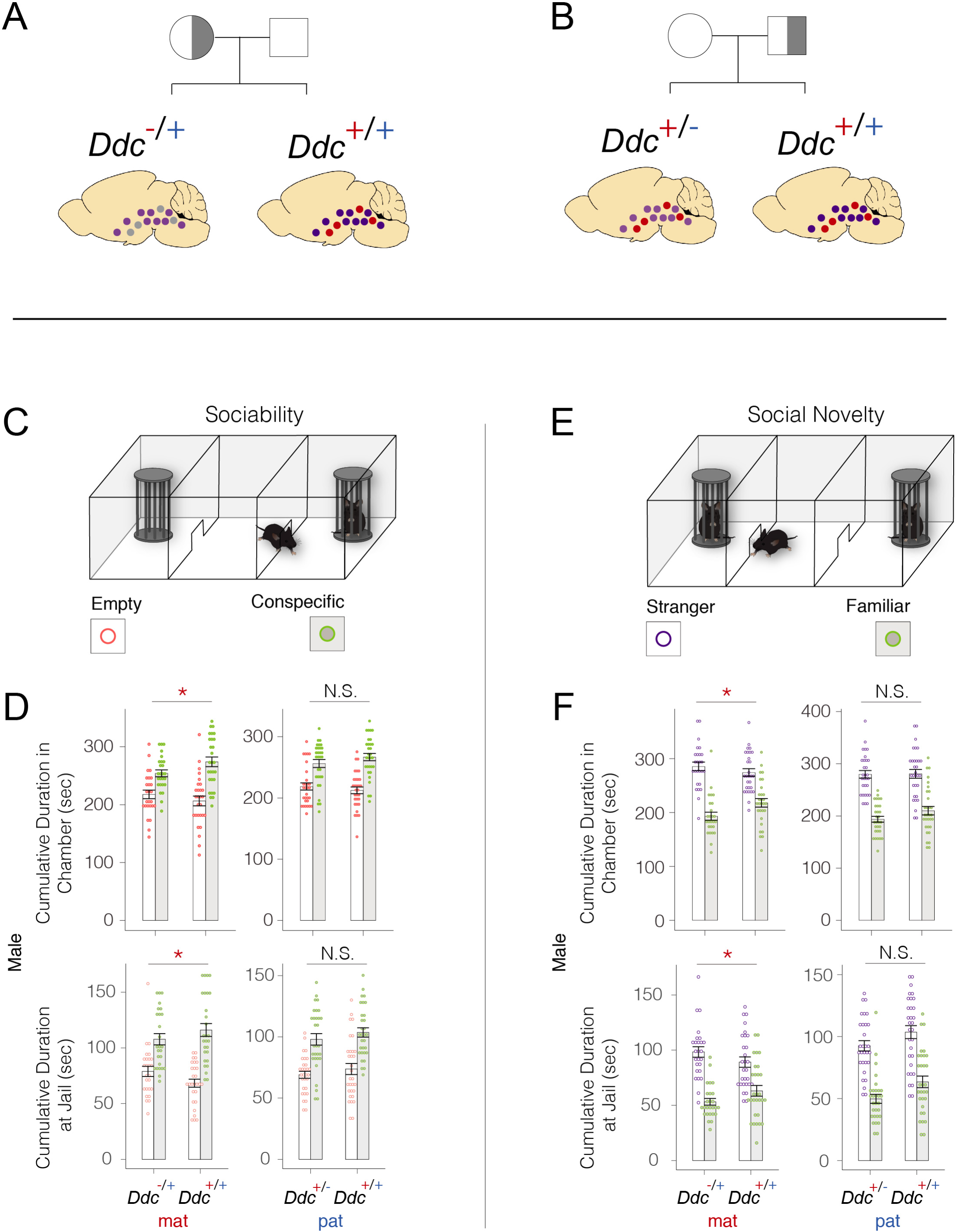
***Ddc* imprinting and loss of maternal allele expression shapes sociability and social novelty seeking in male offspring.** (A-B) Schematics of reciprocal crosses generating mutant maternal (A) and paternal (B) *Ddc* allele heterozygous offspring. Heterozygous dams (half grey circles) were crossed with wild-type sires (white square) to generate *Ddc^−/+^* maternal allele mutants and their *Ddc^+/+^* wt siblings (A). Wt dams (white circles) were crossed with het sires (white square) to generate *Ddc^+/−^* paternal allele mutant and their wt siblings (B). Brain sketches depict monoaminergic neuron populations with biallelic (purple circles), unaffected maternal allele dominant (red circles), and functionally compromised maternal allele dominant (grey circles) *Ddc* expression. (C-D) The data show phenotypic effects uncovered in the three-chambered test for sociability (C). A significant difference in the cumulative duration in the empty (white bars) versus conspecific (grey bars) chamber (top graphs) and in the empty versus conspecific jail area (bottom graphs) was observed for *Ddc ^−/+^* maternal allele mutant heterozygous males compared to their *Ddc^+/+^* littermates (mat; n=26 het, n=28 wt) (D). Mutants spend less time in the conspecific chamber/jail area compared to the empty chamber/jail. A significant effect was not observed for *Ddc ^+/−^* paternal allele mutants (D, pat; n=30 het, n=31 wt). (E-F) The data show phenotypic effects uncovered in the three-chambered test for social novelty (E). Significant phenotypic effects were uncovered in *Ddc ^−/+^* maternal allele mutant heterozygous males compared to their *Ddc^+/+^* littermates (mat) (F), but not in *Ddc ^+/−^* paternal allele mutant heterozygous males (F, pat). *Ddc ^−/+^* males spend relatively more cumulative time in the chamber (F, top graph, mat) and in the jail area (F, bottom graph, mat) with the novel stranger compared to the familiar conspecific. Statistical analyses involved a generalized linear mixed effects model testing for an interaction between chamber/jail and genotype. *P<0.05, N.S., not significant.

Mice prefer to investigate novel mice in their environment compared to familiar mice (Moy et al., 2004). We tested this preference by measuring the time spent in the chamber with a novel mouse (Stranger) versus a familiar mouse (Familiar) (**Fig. 4E**). We found that *Ddc^−/+^* males with a mutant maternal allele, exhibit a significantly increased preference for a novel versus familiar conspecific compared to their *Ddc^+/+^* littermates (**Fig. 4F**, mat, p=0.018). A significant effect was not observed for *Ddc^+/−^* males with a mutant paternal allele (**Fig. 4F**, pat). Our finding is supported by other measures, including the cumulative duration spent near the novel male jail compared to the familiar male jail (**Fig. 4F**, p=0.022). Significant effects were not observed in females. Overall, our results support our primary hypothesis that loss of the maternal *Ddc* allele affects offspring social behavior and the effects differ from loss of the paternal allele. In addition, we found sex dependent effects in that social behaviors in sons are the most strongly affected.

### Testing ethological roles for *Ddc* imprinting: The modular architecture of foraging uncovered in reciprocal *Ddc* heterozygous mutants and littermate controls

In addition to social behaviors, monoaminergic signaling modulates neural systems linked to reward, anxiety, stress, feeding, learning & memory, arousal, sensorimotor functions and other aspects of behavior. Foraging is a rich ethological behavior involving these neural systems. The developmental transition from maternal care to independent foraging changes offspring demands on maternal resources – a proposed driver of imprinting evolution (Haig, 2000a; Lee, 1996). We recently introduced a behavioral paradigm and statistical methodology to study the mechanistic basis of foraging at the level of finite behavioral sequences that we call modules (Stacher Hörndli et al., 2019). Here, we tested the hypothesis that loss of the maternal *Ddc* allele significantly affects offspring foraging and the effects differ from loss of the paternal allele. Moreover, we tested the <u>secondary hypothesis</u> that the maternal and paternal *Ddc* alleles control the expression of distinct foraging modules in offspring. Alternatively, loss of either parental allele may have a generalized effect on all types of foraging sequences or the maternal and paternal alleles affect the same foraging sequences, but the nature of the effect differs.

In our paradigm, the mouse home cage is attached to a foraging arena by a tunnel and mice are permitted to forage spontaneously in two 30-minute phases. During the Exploration phase, naïve mice express behavioral sequences related to the exploration of a novel environment and the discovery and consumption of food in a food patch (pot 2) (**Fig. 5A**). Four hours later, the mice are again given 30 min of access to the arena in the Foraging phase, in which the seeds are now buried in the sand in a new location in pot 4 (**Fig. 5B**). In this second phase, mice express behavioral sequences related to their expectation of food in the former food patch in the now-familiar environment, and also to the discovery of a new, hidden food source (pot 4). By deeply analyzing round trip excursions from the home using an unsupervised machine-learning framework we call “DeepFeats”, we uncover finite, reproducible behavioral sequences, which we call modules (**Fig. 5C**). Here, we investigated how *Ddc* imprinting effects shape foraging module expression in offspring.

**Figure 5.**
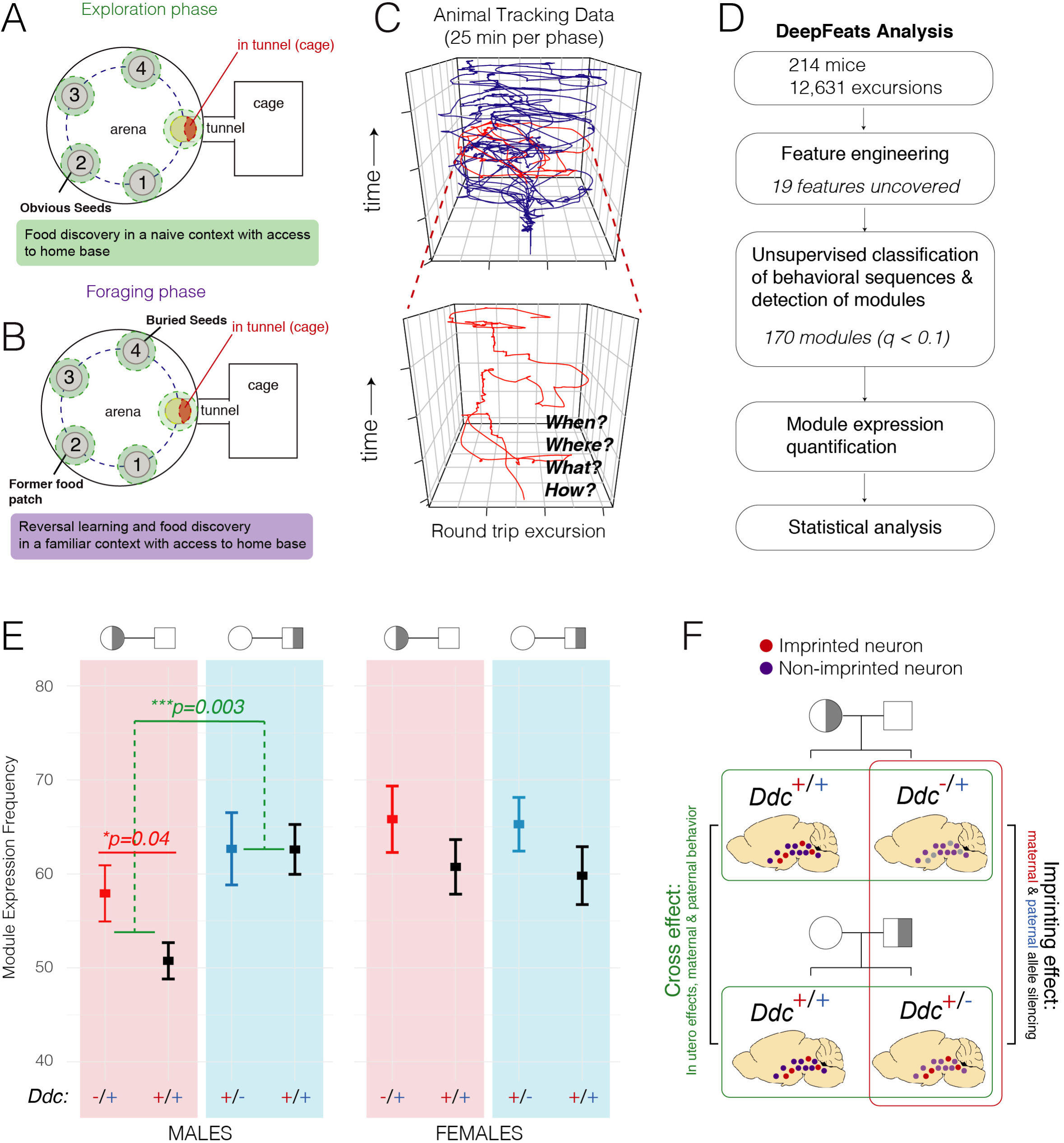
**Modules of economic behavior expressed during foraging reveal *Ddc*-mediated parental effects in adult male and female mice.** (A and B) The schematics show the Exploration (A) and Foraging (B) phase arena layouts to study foraging patterns in a naïve versus familiar context, respectively. In the Exploration phase, the seeds are placed on top of the sand in pot 2. In the Foraging phase, the seeds are moved and buried in the sand in pot 4. (C) DeepFeats analysis segments complex foraging patterns (blue trace) into individual round trip excursions (red trace) from the home and captures measures that describe what the animals do, where they go, how they move and when the excursion is expressed (see Methods). (D) DeepFeats analysis workflow of foraging patterns expressed by the cohort of adult male and female *Ddc ^−/+^*, *Ddc ^+/−^* and *Ddc^+/+^* mice in the study. DeepFeats uncovered 170 modules of economic behavior from 12,631 round trip excursions performed by the mice. (E) The plot shows the aggregated expression of modular excursions by males and females derived from the two reciprocal genetic crosses in the study (D and E). The data reveal a significant main effect of cross influences the expression of modular excursions in males (p=0.003, likelihood ratio test, generalized linear model, Gaussian distribution with log link function). Statistical modeling that absorbs variance due to the cross effect uncovered a significant main effect of the maternal allele genotype (p=0.04; + versus -), but not the paternal allele (p=0.99; + versus -). Significant cross or imprinting effects were not observed in females. Mean±SEM; N = 24-30 animals per genotype and sex; *p<0.05. (F) Schematic depicts the cross between *Ddc* heterozygous mothers X wildtype fathers to generate *Ddc ^−/+^* and *Ddc^+/+^* littermate offspring (top), and the reciprocal cross between *Ddc* heterozygous fathers X wildtype mothers to generate *Ddc ^+/−^* and *Ddc^+/+^* offspring (bottom). This paradigm enables us to uncover the behavioral effects of *Ddc* imprinting and link loss of maternal versus paternal allele function to specific behavioral phenotypes (red box), and reveal and account for cross effects due to heterozygosity of the parents independent of offspring genotype (green boxes).

To begin, we do not know the nature or modular architecture of the foraging patterns expressed by adult *Ddc^−/+^* and *Ddc^+/−^* male and female offspring and *Ddc^+/+^* littermate controls. To investigate, we profiled foraging for 214 mice, including the following genotypes: (1) <u>Females</u>, n=103: *Ddc^−/+^* n= 27, *Ddc^+/−^* n=25, *Ddc^+’/+^* n=24 (‘ indicates heterozygous mother), *Ddc^+/+’^* n=27 (‘ indicates heterozygous father); and (2) <u>Males</u>, n=111: *Ddc^−/+^* n= 29, *Ddc^+/−^* n= 24, *Ddc^+’/+^* n=28, *Ddc^+/+’^* n=30. We captured data for 12,631 round trip foraging excursions from the home and DeepFeats analysis found 19 measures (aka. features) that maximize the detection of candidate modules (**Fig. 5D and S4A-C**). Moving forward with these measures, we partitioned the data into training and test datasets balanced by sex, genotype and cross to identify significantly reproducible clusters of similar behavioral sequences that denote modules (**Fig. S4D**). Our analysis found 170 significant modules from the 12,631 excursions (q<0.1, In-Group Proportion (IGP) permutation test) (**Fig. 5D and S4C,D**). Each module is numbered based on the training data clustering results. Having defined the modular architecture of the cohort’s foraging patterns, we analyzed the results to test our hypotheses.

### Foraging reveals that *Ddc* mediates multiple forms of parental influence on offspring behavior

To determine whether *Ddc* imprinting affects the overall expression of foraging modules, we counted the number of modules expressed by each mouse, which is a measure of the number of times they enter the arena, and analyzed the data according to sex and genotype. Unexpectedly, we found that *Ddc^−/+^* and *Ddc^+/+^* male offspring generated from the cross between *Ddc* heterozygous mothers and C57BL/6J (B6) fathers express significantly fewer foraging modules than *Ddc^+/−^* and *Ddc^+/+^* males generated from the reciprocal cross (p=0.003, likelihood ratio test for cross effect, Gamma distribution **Fig. 5E**). The data show female offspring are not significantly affected (**Fig. 5E**). Indeed, a significant interaction effect between sex and cross was observed after absorbing variance due to *Ddc* genotype (het vs wt) in the offspring (p=0.03, likelihood ratio test). Therefore, *Ddc* heterozygosity in the parents causes a significant phenotypic effect in sons independent of the son’s genotype (heterozygous or wildtype). We refer to this form of parental effect as a “cross effect” and it could be caused by the *in utero* environment or behavior of *Ddc* heterozygous mother or father.

The presence of significant cross effects on offspring foraging requires that we absorb variance due to the cross effect prior to testing for imprinting effects. We used this nested modeling strategy to test our primary hypothesis that loss of the maternal *Ddc* allele causes behavioral effects that differ from loss of the paternal allele in sons and daughters. We found that the main effect of the paternal allele genotype (wildtype (+) versus mutant (-) allele) on module expression is not significant (sons: p=0.9, daughters: p=0.2; likelihood ratio test), indicating that loss of the paternal allele does not significantly affect overall module expression levels (**Fig. 5E**). In contrast, after absorbing variance due to cross and paternal allele effects with a nested model, we found a significant main effect of the maternal allele genotype (+ versus -) in sons (p=0.04, **Fig. 5E**). Significant effects were not observed in daughters (p=0.3, **Fig. 5E**), though this does not preclude the possibility of effects on specific modules that are not revealed from overall module expression. Overall, the results support our primary hypothesis that loss of the maternal *Ddc* allele in offspring significantly affects foraging behavior and has phenotypic effects that differ from the paternal allele. Additionally, we found *Ddc* imprinting effects on behavior depend upon the sex of the offspring and uncovered *Ddc*-mediated cross effects on behavior (**Fig. 5F**).

### *Ddc* imprinting and cross effects control the expression of groups of related foraging modules

Foraging modules are discrete behavioral sequences that differ in terms of their form, function, expression timing, context of expression and regulation (Stacher Hörndli et al., 2019). However, some modules have similarities. For example, multiple different modules can be linked to food patch exploitation, each involving a different approach to a common goal - food consumption. Our data reveal *Ddc* mediates multiple different parental effects on offspring, including maternal and paternal imprinting and cross effects. Here, we determine whether the 170 modules uncovered in these mice include groups of related modules and whether *Ddc*-mediated imprinting and/or cross effects significantly impact entire groups of related module. Alternatively, different *Ddc*-mediated paternal effects might only act at the level individual modules, rather than affecting module groups. This analysis begins to test our secondary hypothesis that *Ddc* imprinting effects impact specific modules and the maternal and paternal *Ddc* alleles control distinct subsets of foraging modules. Additionally, we test whether cross effects impact specific modules.

We began by defining groups of related modules. For each module, we computed the centroid, defined as the average values of the 19 measures that cluster behavioral sequences together in a module. Centroids summarize how the different behavioral measures delineate different module types and we display them in a heatmap (**Fig. 6A**). Unsupervised hierarchical clustering of the centroid data revealed 23 module groups (**Fig. 6A)** and seven partitions that are singleton modules that did not group with others (**Fig. 6A**, eg. module-74). For each module group, tested whether the aggregated expression of the modules in the group is significantly affected by cross effects, the genotype of the paternal allele and/or the maternal allele. We found that 27% of groups exhibit a significant main effect of cross in males, and no cross effects in females (**Fig. 6B;** p<0.05 in green columns; summary in **Fig. 6C**). These results reveal that cross effects impact the expression of specific groups of modules in sons. For example, the expression of module Group-1 is decreased in *Ddc^+/+^ and ^−/+^* sons born from *Ddc* heterozygous mothers compared to sons from wildtype mothers (**Fig. 7A**). In contrast, there are no Group-1 cross effects in daughters (**Fig. 7A**) and some other module groups do not have cross effects in either sex, such as Group-5 (**Fig. 7B** and **see 6B**, p>0.05, green columns).

**Figure 6.**
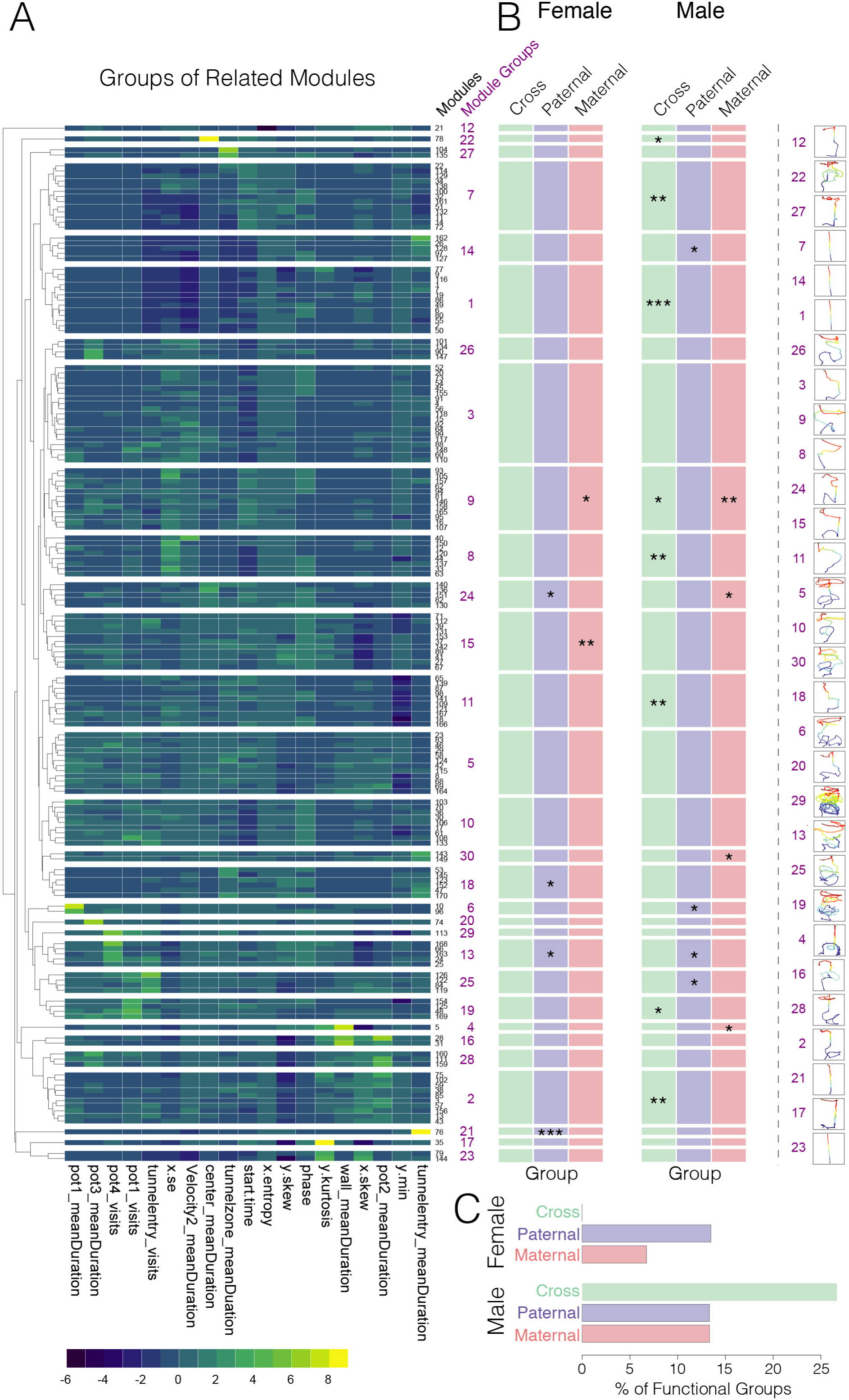
**Identification of modules of foraging behavior and parental effects from *Ddc* ^−/+^, *Ddc* ^+/−^ and *Ddc*^+/+^ adult male and female mice.** (A) The heatmap shows the centroids for each of the 170 significant behavior modules uncovered in the mice and the relative weights of the 19 measures (x-axis) capturing the behavioral sequences assigned to each module. Modules are numbered according to the training data clusters they arose in (black type, right side y-axis). Unsupervised hierarchical clustering was performed to define groups of modules with similar patterns. The results reveal 30 groups of related modules (purple type, right side y-axis). (B) The chart summarizes the results of the statistical modeling performed on each group of related modules and the results correspond to the groups in the heatmap (A). The significant effects of cross (green), paternal allele genotype (blue) and maternal allele genotype (red) on the expression of a group of modules are indicated in the colored columns. The colored traces on the far right show examples of the movement tracking patterns for modules in each grouping and show different groups involve different movement patterns. Movement traces are ordered according to the heatmap results. *p<0.05, **p<0.01, ***p<0.001. N = 24-30 animals per genotype and sex. (C) The bar plots show the percentage of module groups with significant cross, paternal and maternal effects. The color scheme matches the scheme of the chart (B).

**Figure 7.**
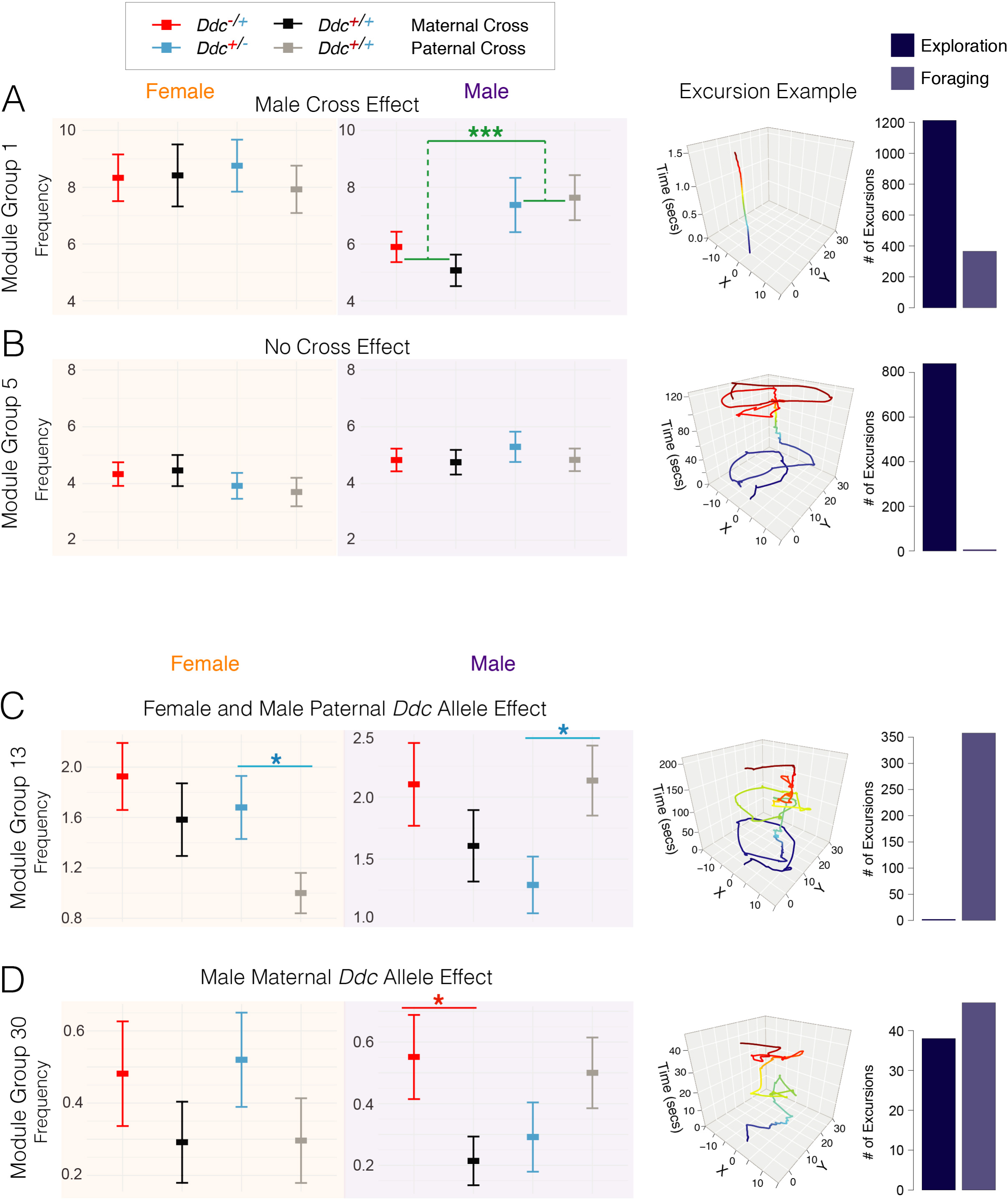
***Ddc* cross and imprinting effects shape foraging module expression in a sex-dependent manner.** (A and B) The plot shows the expression of Group-1 modules is significantly impacted by cross effects in males, but not females (A). This effect is group specific and does not significantly affect other groups of modules, such as Group-5 (B). On the right side, representative traces of the X-Y movement patterns over time (seconds) for Group-1 and Group-5 modules are shown and they differ. The barplots of the number of Group-1 and Group-5 modules expressed in the Exploration (dark blue) versus Foraging (light blue) phases show that both module groups are context dependent and preferentially expressed in the Exploration phase. Green stars indicate significant main effect of cross (see Figure 6B). Genotypes indicated by top legend. (C) The plots show the expression frequency of Group-13 modules is impacted by *Ddc* imprinting and significantly affected by the genotype of the paternal allele in females and males. A sex difference is observed, such that *Ddc^+/−^* females exhibit increased expression compared to *Ddc^+/+^* littermate controls, while males show the opposite effect. Blue stars indicate the significant main effect of paternal allele genotype (see Figure 6B). A representative trace of Group-13 movement patterns is shown to the right and the barplot reveals preferential expression of these modules in the Foraging phase. (D) The plots show sex dependent imprinting effects on the expression frequency of Group-30 modules, which exhibit a significant main effect of the maternal allele in males. Red star indicates the significant main effect of the maternal allele genotype (see Figure 6B). The trace shows Group-30 movement patterns are distinctive from the other modules shown (A-C) and are expressed in the Exploration and Foraging phases. Mean±SEM; N = 24-30 animals per genotype and sex; *p<0.05, **p<0.01, ***p<0.001.

After absorbing variance due to cross effects, we uncovered module groups significantly affected by *Ddc* imprinting. From males and females, we found 7 groups significantly affected by the paternal *Ddc* allele genotype (- versus +) (**Fig. 6B;** p<0.05, blue columns, main effect of paternal allele) and 5 different groups are affected by the maternal *Ddc* allele genotype (**Fig. 6B;** p<0.05, red column, main effect of maternal allele) (totals in **Fig. 6C**). Thus, the maternal and paternal *Ddc* alleles significantly impact different groups of modules. Moreover, we found that the affected module groups differ between males and females (**Fig. 6B**). For example, the expression of Group-13 is affected by the genotype of the paternal *Ddc* allele, but not the maternal allele, and the effect is sexually dimorphic (**Fig. 7C**). Group-13 expression is decreased in *Ddc^+/−^* sons compared to littermate controls, but significantly increased in *Ddc^+/−^* daughters (**Fig. 7C**). On the other hand, Group-30 is affected by the maternal allele genotype, but not the paternal allele, and the parental effect occurs in sons, but not daughters (**Fig. 7D**).

Interestingly, Group-13 modules are distinguished by strong interactions with the Foraging phase food patch (pot 4) (**Fig. 6A,** increased “pot4_visits”; and **Fig. 7C**). Feeding in the naïve Exploration phase is associated with Group-2, Group-28 and Group-16 modules, which are distinguished by centroids showing more time at the Exploration phase food patch (pot 2) (**Fig. 6A,** increased “pot2_mean duration”). However, we did not find significant paternal or maternal allele effects on these groups of Exploration phase feeding modules in sons or daughters (**Fig. 6B;** p>0.05, blue and red columns in males and females). These results suggest the paternal *Ddc* allele affects feeding-related behaviors in the Foraging phase, but not the Exploration phase, revealing the importance of the context for revealing phenotypic effects. Moreover, the maternal *Ddc* allele genotype did not significantly affect any of these feeding-linked module groups, and cross effects significantly impacted Group-2 modules in sons. Our data show that *Ddc*-mediated maternal and paternal imprinting effects and cross effects differentially affect groups of related modules in sons and daughters, leaving open the question of how these parental effects extend to the level of individual modules.

### Identification of individual foraging modules linked to specific *Ddc*-mediated parental effects

Individual modules are discrete behavioral sequences that we expect are linked to different neural and molecular mechanisms and serve different functions in different contexts. Parents could shape offspring behavior patterns by modulating the expression of individual modules. Alternatively, parental effects might only affect behavior broadly, by affecting module groups or overall behavioral drives, such as hunger. A simple change to hunger, for example, would presumably manifest as a net increase or decrease to all feeding-related modules, rather than changes to specific module groups or individual modules. We tested how *Ddc* imprinting and cross effects manifest at the level of the individual modules. To begin, we investigated the possibility that parental effects cause novel modules to form in offspring. However, we found that all 170 modules are expressed by *Ddc^+/+^* males and females derived from the cross with a *Ddc^+/+^* wildtype mother, which shows that none of the modules are <u>only</u> expressed in response to a particular parental effect. Thus, parental effects do not appear to cause the formation of new modules in offspring. Instead, the parental effects might shape the expression of a baseline set of modules.

We first assessed whether individual modules within module groups are differentially expressed and therefore behave like distinct units of behavior. In support, our generalized linear model found that 78% of module groups have a significant main effect of individual “module type” in males and/or females, indicating that individual modules within these groups differ significantly in their expression frequency (**Fig. S5B,C**, p<0.05, main effect of module, dark green Module column). This result suggests the individual modules are distinct units of behavior. To determine how individual modules are impacted by *Ddc*-mediated parental effects, we examined interactions between module type and different parental effects on module expression, including cross effects, paternal allele genotype and maternal allele genotype (**Fig. S5B,C**). Post-tests revealed the identities of the affected modules. We found that XXX% of module groups have a significant interaction between module type and one or more parental effects (**Fig. S5C**), revealing differential effects at the level of individual modules. In one example, Group-9 modules have significantly increased aggregated expression in *Ddc^−/+^* in sons and daughters (**Fig. 8A,B**). However, at the level of individual modules, we found a significant interaction between module type and maternal allele genotype in sons, revealing module specific effects (**Fig. 8C**). Post-tests showed that modules -16, -157 and -158 in Group-9 exhibit increased expression in *Ddc^−/+^* sons compared to littermate controls (**Fig. 8C**). We also observed that module-94 is uniquely impacted by cross effects (**Fig. 8C**). These individual modules differ in terms of their structure and expression context (ie. phase), suggesting different functions (**Fig. 8D**).

**Figure 8.**
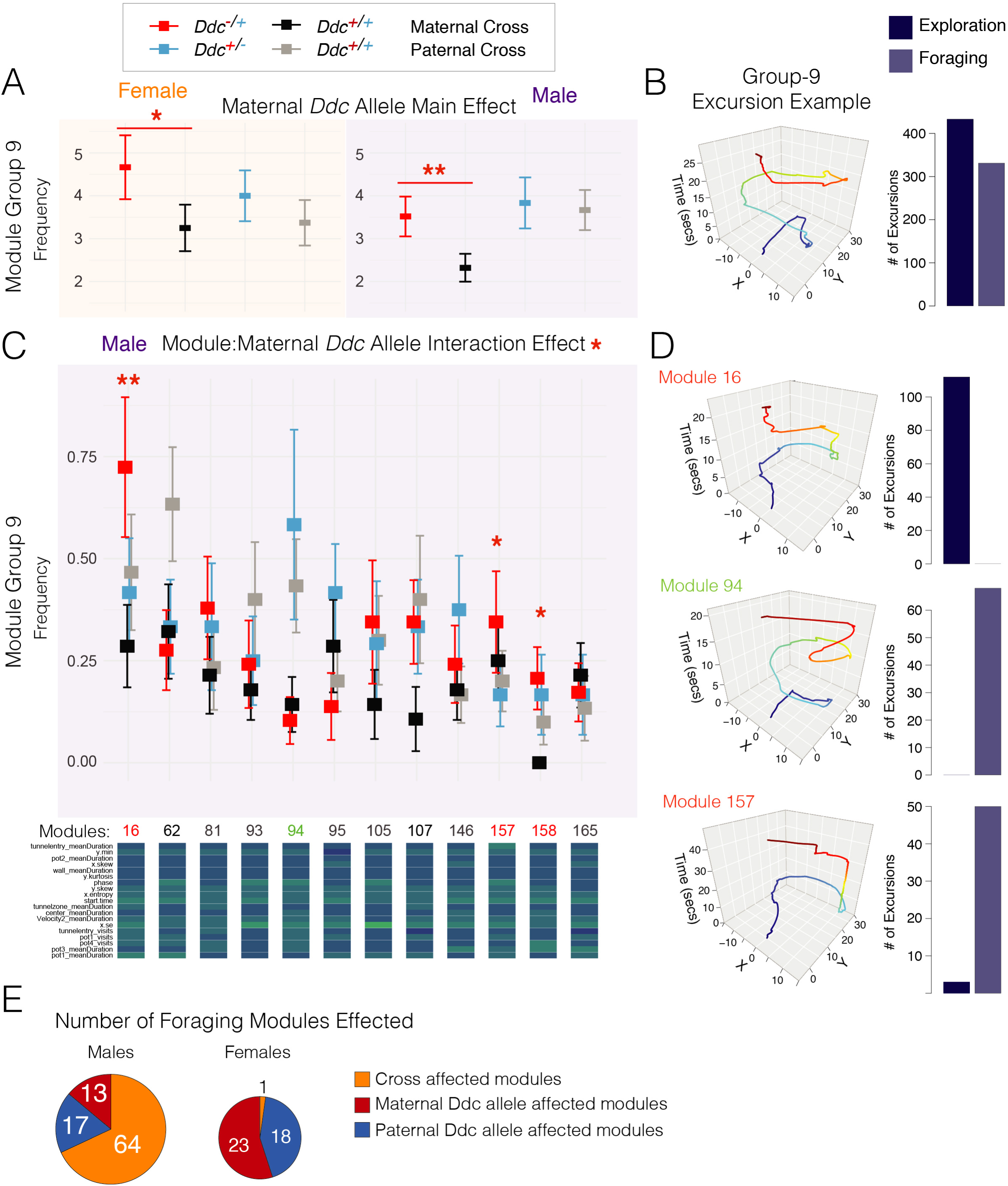
***Ddc* imprinting effects impact the expression of individual modules of foraging behavior in offspring.** (A and B) The plots show the expression frequency of Group-9 modules is significantly impacted by a main effect of the genotype of the maternal *Ddc* allele in females and males (A). A representative movement trace of Group-9 modules is shown and the barplot shows that these modules are expressed in the Exploration and Foraging phases (B). Genotypes indicated by top legend. Mean±SEM; N = 24-30 animals per genotype and sex. (C) The plots show the expression frequency of each module in Group-9 by genotype and cross for males. A significant interaction between module and maternal allele was found (Figure S5B) and the data show the identities of the individual modules with significant effects after absorbing variance due to the cross (module-16, module-157, module-158). Module-94 is highlighted as an example of an individual module with a cross effect. The relative weights of the measures characterizing each module in the group are shown in the heatmap below (data from Figure 6A), indicating differences between Group-9 modules. Red stars indicate modules significantly contributing to the module:maternal allele interaction effect. *p<0.05, **p<0.01. Mean±SEM; N = 24-30 animals per genotype. (D) The traces show representative movement patterns for individual modules affected by the maternal allele genotype (module-16 and -157). The module affected by cross (module-94) is shown for contrast, showing the behavioral sequences differ for individual modules, despite overall similarities. Moreover, the barplots show that the individual modules differ in terms of frequency of expression in the Exploration versus Foraging phase contexts. (E) The pie charts show the total numbers of foraging modules significantly impacted by parental effects, including the maternal *Ddc* allele genotype (red), paternal *Ddc* allele genotype (blue) and cross effects (orange) in males and females. The counts include all modules in module groups with a significant parental effect and the specific modules impacted for module groups that have a significant interaction effect between module type and parental effect (see Figures 6 and S5).

Overall, *Ddc* cross, paternal allele and maternal allele effects extend to the level of individual modules and shape offspring behavior patterns, in part, by affecting these discrete units of behavior. We tallied the total numbers of modules significantly impacted by each of the different *Ddc*-mediated parental effects (**Fig. 8E**). Finally, in an extended behavior analysis presented in a Supplemental Data section (**Data S3** and **S4**), we uncovered additional *Ddc* cross, paternal allele and maternal allele effects on specific aspects of social behavior (**Data S3**) and in standard lab tests of exploratory behaviors in males and females (**Data S4A-D**). We also found that <u>none</u> of the *Ddc*-mediated parental effects significantly alter offspring body weights, indicating no obvious growth or obesity effects (**Data S4E**). These supplemental data further support our major finding that *Ddc* mediates multiple different parental effects on offspring behavior.

## DISCUSSION

Genomic imprinting is an epigenetic mechanism through which mothers and fathers differentially affect gene expression in offspring. However, the cellular nature and functional effects of noncanonical imprinting (aka. parental allele biases), and the complex imprinting effects impacting *Ddc,* are not well understood. Here, we defined noncanonical imprinted genes with imprinting effects linked to neuronal versus non-neuronal brain cells and found an allele bias persists at this level of cellular analysis. We then used allelic reporter mice to get finer cellular resolution of noncanonical imprinting for *Ddc* and uncovered complexities. In adults, dominant expression of the <u>maternal</u> *Ddc* allele was revealed in subpopulations of cells in 14 brain regions, most of which are in the hypothalamus and we found an association with GABAergic neurons. Dominant expression of the <u>paternal</u> and <u>maternal</u> *Ddc* alleles was found in different subpopulations of adrenal cells. Paternal allele expression was also observed in subpopulations of embryonic cardiac cells. We found DDC+ cells that co-express both parental alleles in all examined tissues. Functional studies determined that *Ddc* imprinting effects interact with inherited heterozygous *Ddc* mutations and shape offspring social, foraging and exploratory phenotypes, but not body weight. Machine learning dissections of foraging in *Ddc^−/+^*, *Ddc^+/−^* and *Ddc^+/+^* offspring uncovered 170 discrete modules of economic behavior and parental effects at different levels of analysis. The maternal and paternal *Ddc* alleles are revealed to affect different module groups and individual modules. Additionally, we uncovered *Ddc* cross effects in which specific modules are affected in sons according to the *Ddc* genotype of the parents independent of the offspring genotype. Sexually dimorphic cross and imprinting effects were uncovered in social, foraging and exploratory behavior tests. Our results are foundational in that they reveal noncanonical imprinting creates functional cellular diversity, *Ddc* mediates multiple different parental effects on behavior and how these effects shape ethological behavior in sons and daughters at different levels.

### Noncanonical Imprinting as a Functional and Cell-Dependent Form of Allele-Specific Gene Regulation

Noncanonical imprinted genes that exhibit a bias to express one allele higher than the other have been described by us, and others (Andergassen et al., 2017; Bonthuis et al., 2015; Crowley et al., 2015; Perez et al., 2015). It is debated whether these effects are functional or an epiphenomenon. Our current study supports a functional role for these effects by revealing that (1) the *Ddc* maternal allele bias in the adult mouse brain involves dominant maternal allele expression in discrete subpopulations of brain cells and manifests at the protein level, (2) the effects are linked to GABAergic neurons, (3 we uncovered subpopulations of adrenal cells with maternal and paternal *Ddc* imprinting, and (4) loss of the maternal *Ddc* allele affects behavior differently from loss of the paternal allele. In some cells, we observed an allele bias and it is unclear whether these cellular allele biases are functional and contribute to behavioral changes or whether the behavioral effects are related to cells with allele silencing effects. Cell-type dependent imprinting is a known phenomenon (Gregg, 2014; Perez et al., 2016) and, as our understanding of the cellular regulation and function of noncanonical imprinting grows, the value of differentiating these effects from canonical imprinting may change.

MEGs and PEGs are postulated to function antagonistically in offspring (Haig, 2000a). Our analysis of single cell RNASeq data revealed MEGs and PEGs co-expressed with *Ddc* at the cellular level in the hypothalamus, including the noncanonical MEG, *Th (Tyrosine hydroxylase),* the noncanonical PEGs, *Gpr1 (G-protein coupled receptor 1),* and *Sec14l3 (SEC14 Like Lipid Binding 3),* and the canonical PEG, *Dlk1 (Delta-like kinase 1).* These genes may be additional sources of maternal and paternal influence in DDC+ cells, revealing an axis of parental control in the monoamine system. Additionally, various approaches have uncovered functional subpopulations of DA, NE and 5-HT neurons (Berke, 2018; Farassat et al., 2019; Gershman and Uchida, 2019; Okaty et al., 2019). Determining the connectivity patterns and physiological properties of monoaminergic cell populations distinguished by different imprinting effects will deepen our understanding of the functional cell types and circuits. Finally, studies profiling imprinting effects in mice differ in terms of their power to detect different effects and the total numbers of imprinted genes are debated (Andergassen et al., 2017; Babak et al., 2015; Bonthuis et al., 2015; Crowley et al., 2015; Perez et al., 2015). With large sample sizes and a sensitive statistical method, we, and others, found noncanonical imprinting is more prevalent in the mouse genome than canonical imprinting (Bonthuis et al., 2015; Crowley et al., 2015). However, the cellular nature and functions of most cases have yet to be studied.

### The *Ddc* Locus is a Hub of Parental Control On Offspring Behavior: Imprinting and Cross Effects within the Monoamine System

Previous studies uncovered various mechanisms of parental influence on offspring phenotypes, including the effects of maternal and paternal behavior (Alter et al., 2009; Curley and Champagne, 2016), the *in utero* environment (Lindsay et al., 2019), maternally imprinted genes and paternally imprinted genes (Haig, 2000a; Keverne, 2001; Perez et al., 2016). Our data indicate that *Ddc* is a single genetic locus that contributes to many, and perhaps all of these different parental effects. We found *Ddc* exhibits imprinting of the paternal and/or maternal allele in subpopulations of DDC+ adult brain cells, adrenal cells, and developing cardiac cells. *Grb10,* which is located next to *Ddc* in the genome, exhibits a different imprinting pattern, involving dominant expression of the paternal allele in the brain and the maternal allele in non-neural tissues (Arnaud et al., 2003; Blagitko et al., 2000; Hitchins et al., 2001; Miyoshi et al., 1998). Most imprinted genes exhibit imprinting of the <u>same</u> parental allele across all affected tissues and developmental stages (Andergassen et al., 2017; Bonthuis et al., 2015; Crowley et al., 2015; Perez et al., 2015). Therefore, *Ddc* and *Grb10* imprinting effects are atypical, suggesting they are subject to a unique crucible of maternal and paternal evolutionary pressures. In addition to these unusual imprinting effects, we found *Ddc*-mediated cross effects on behavior. Collectively, the data suggest that *Ddc* has evolved to be a hub mediating multiple different forms of parental influence on offspring behavior in mice.

Our data suggest the intriguing possibility that maternal and paternal *Ddc* allele imprinting shapes the functionality of the adult Hypothalamic-Pituitary-Adrenal (HPA) axis (Kvetnansky et al., 2009), with dominant maternal expression in several hypothalamic cell populations and a mixture of adrenal cells with dominant paternal and maternal allele expression. This axis plays key roles in modulating behavior and stress responses, suggesting that *Ddc* imprinting in the HPA axis is a source of parental control over these responses. Additionally, *Ddc* imprinting in some embryonic cardiac cells suggests the possibility of phenotypic effects caused by imprinting and monoamine signaling changes at critical developmental stages. We may not yet know all of the ages, tissues and cell populations impacted by *Ddc* imprinting. Further, future studies using conditional deletions of the maternal versus paternal *Ddc* alleles in specific cell populations are now needed to link cellular imprinting effects to specific behavioral phenotypes and modules. Conditional genetic approaches will also help to further dissect *Ddc* imprinting effects from cross effects.

*Ddc*-mediated cross effects on offspring behavior could be caused by changes to maternal physiology, the *in utero* environment and/or parental behaviors. A previous study found that *Tph1^−/−^* mothers lacking the enzyme synthesizing peripheral 5-HT have smaller embryos independent of the genotype of the offspring (Côté et al., 2007). The authors did not report an effect in *Tph1* heterozygous mothers, which were used as controls. Our study found that *Ddc* heterozygosity in parents significantly affects offspring behavior without significant effects on body weight. Future studies are needed to determine whether *Ddc* cross effects are mediated by 5-HT, DA, NE and/or E signaling. Roles for maternally-derived 5-HT in embryonic development are the best understood (Brummelte et al., 2017). Finally, imprinted genes can significantly affect parental behaviors, such as *Peg1/Mest* (Lefebvre et al., 1998) and *Peg3* (Champagne et al., 2009; Li et al., 1999) (but see (Denizot et al., 2016)). Therefore, *Ddc*-mediated cross effects might also involve changes to maternal or paternal behaviors that in turn shape offspring behavior. Future studies are needed to determine the mechanistic basis of *Ddc* cross effects.

*Ddc* cross effects most strongly impacted the behaviors of male mice. DDC modulation of *in utero* 5-HT levels might influence the development of neural circuits that are sensitive to the effects of sex hormones; thereby shaping behavior in a sexually dimorphic manner. Our data raise the possibility that genetic variants impacting other enzymes required for monoamine biosynthesis could also cause transgenerational effects on offspring behavior. Deep analyses of rich foraging patterns in mouse models could help to test the transgenerational behavioral effects of some known genetic variants associated with human phenotypes (Watanabe et al., 2019). There are potential biomedical implications for our observation because 5-HT levels are proposed to influence autism risks, which is three-fold more prevalent in males than females (Muller et al., 2016; Veenstra-VanderWeele et al., 2012).

Of potential importance is that *Ddc* has been reported to show imprinting in mouse extraembryonic tissues involving paternal allele expression in the visceral yolk sac epithelium and yolk sac (Andergassen et al., 2017). If loss of *Ddc* expression in extraembryonic tissues causes transgenerational effects on offspring behavior, this mechanism could link *Ddc* cross effects and imprinting effects into a unified <u>transgenerational genetic axis of parental influence</u>. Thus, further studies are needed to determine whether the parental origin of a *Ddc* heterozygous mutation in the parents significantly influences *Ddc*-mediated cross effects on offspring behavior and whether environmental factors, like diet, impact cross effects. There may be adaptive advantages to shaping offspring foraging through parental effects on monoamine biosynthesis pathways (Brummelte et al., 2017). Such effects might shape offspring survival and reproductive success by changing the expression of discrete foraging modules to improve offspring foraging success in the environment. Indeed, foraging patterns are under strong selective pressures that shape life histories and must be well adapted to environmental conditions (Lee, 1996; Nislow and King, 2006; Stephens et al., 2007). The functions of parental effects on offspring social behaviors have been considered by others (Haig, 2000b; Isles et al., 2006; Ubeda and Gardner, 2010). Further studies are needed to uncover the potential adaptive advantages of different *Ddc*-mediated parental effects.

Finally, *DDC* imprinting appears to occur in humans, though follow up studies are needed to confirm and better characterize the effects (Babak et al., 2015; Baran et al., 2015). Large-scale genome-wide association studies have linked genetic variation at the *Ddc* locus to risks for multiple major diseases, including bipolar disorder, schizophrenia, depression, addiction, anorexia nervosa, type 2 diabetes, obesity and other biomedically-important behavioral, metabolic, cardiovascular, endocrine and immunological phenotypes (Watanabe et al., 2019). These disorders are phenotypically variable and the mechanisms involved are unclear, but could involve *DDC* imprinting and cross effects.

### Resolving and Regulating the Modular Architecture of Complex Foraging Patterns

Vertebrate economic choices that shape reward, effort and risk evolved under selective pressures for foraging and function to help balance caloric intake with the costs of energetic demands and predation risks (Stephens et al., 2007). Understanding the architecture and regulation of these complex economic behavior patterns is important and fundamental. The field of neuroeconomics proposes that many chronic health problems have roots in reward, effort and risk decisions (DeStasio et al., 2019). In addition, mental illnesses, like bipolar disorder, schizophrenia and depression, have been framed as disorders of neuroeconomic processes (Sharp et al., 2012). However, the mechanisms shaping complex economic behavior patterns are not well understood. Genomic imprinting evolved in mammals and the nursing to foraging transition at weaning uniquely shaped mammalian life histories and brain development (Lee, 1996), suggesting that different parental and imprinting effects may strongly shape mammalian foraging and economic behaviors. Previously, we found support for this idea and discovered foraging patterns in mice are constructed from finite, genetically controlled modules defined from round trip excursions from the home (Stacher Hörndli et al., 2019). Here, we have begun to dissect the parental, molecular and cellular mechanisms controlling foraging modules. So far, we found *Ddc*-mediated imprinting and cross effects do not cause new modules to form in offspring, but affect the expression frequency of specific module groups and individual modules in a sex dependent manner. Future studies are needed to test potential effects on module expression timing and sequential order to better understand how *Ddc*-mediated effects shape higher level economic patterns. We also do not know whether complex social behaviors can be dissected into finite modules controlled by *Ddc* or what foraging modules might be revealed in richer and more naturalistic foraging environments. Our understanding of how different parental, molecular and cellular mechanisms shape complex behavior, health and disease phenotypes will improve with deeper machine-learning dissections of behavior in the future.

## SUPPLEMENTAL INFORMATION

The supplemental information includes five supplemental figures, one supplemental table and a supplemental data results section.

## ACKNOWLEDGEMENTS

The statistical methodology in the study was supervised and developed by statisticians in the University of Utah Biostatistics Core Facility, including Drs. Greg Stoddard and Alun Thomas. We thank Drs. Monica Vetter and Jan Christian and members of the Gregg lab for critical reading of the manuscript and input on the study. We thank Dr. Channabasavaiah B. Gurumurthy and Rolen M. Quadros (University of Nebraska Medical Center) for help generating the *Ddc* allelic knock-in reporter lines. Single cell RNASeq studies were performed in the University of Utah High Throughput Genomics Core facility. This work was supported by funding from the National Institutes of Health (R01AG064013, R21MH120468, R01MH109577 and R21MH118570 grants to C.G.) and a K99 award to P.B. (K99MH111912).

## AUTHOR CONTRIBUTIONS

P.B. and S.S. performed studies of *Ddc* allelic reporter mice; P.B. and S.S. performed the behavior studies; P.B., S.S., J.E, C.S.H. and C.G. analyzed the behavior data; W-C.H., E.F, S.K. and C.G. performed the genomics studies; P.B., S.S. and C.G. wrote the manuscript with input from all of the authors.

---

**Figure S1.**
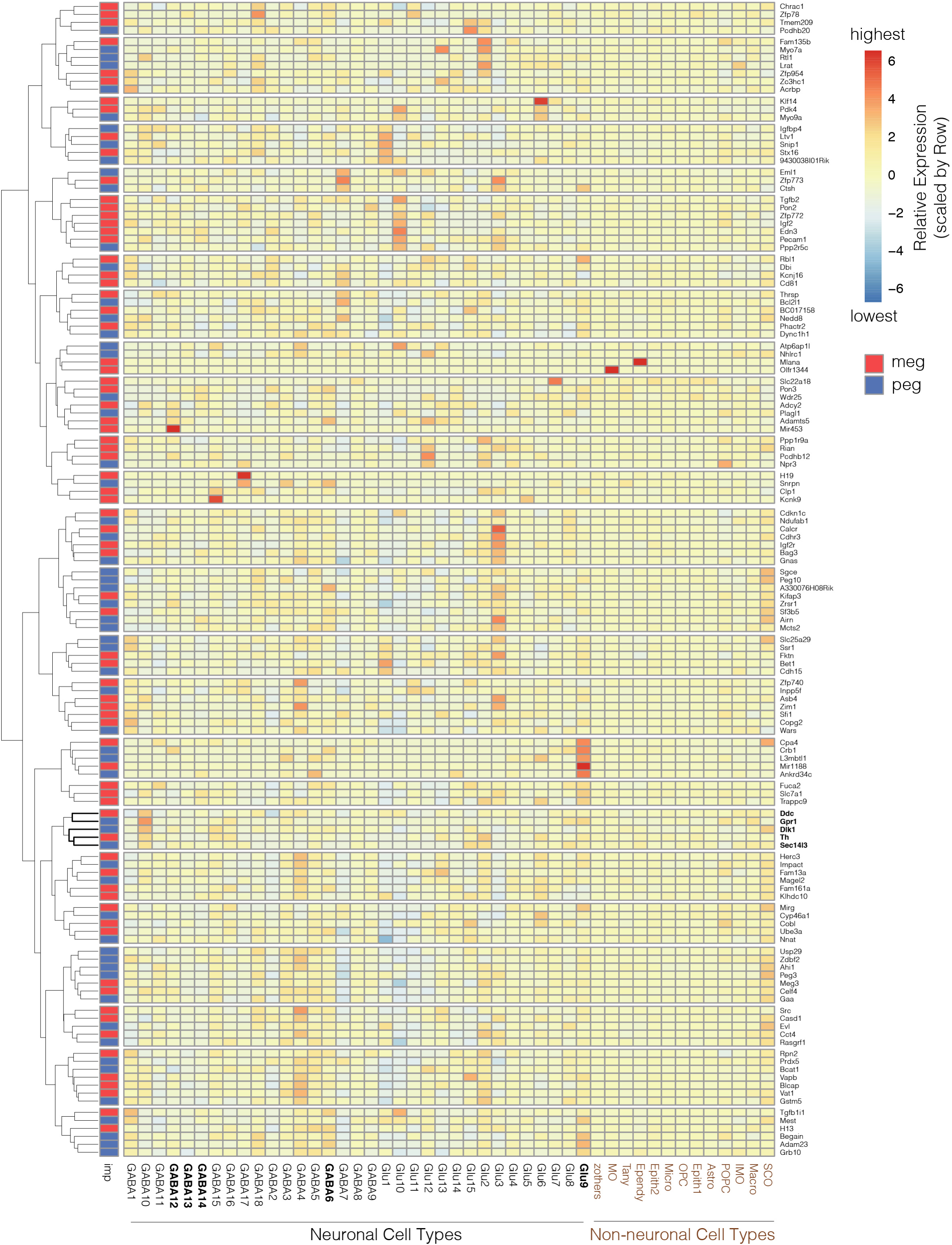
**Mouse hypothalamus scRNA-Seq data reveals sets of MEGs and PEGs with similar expression in specific hypothalamic neuronal and non-neuronal cell-types.** Related to Figure 1. The heatmap depicts the mean expression level of each imprinted gene computed from normalized read counts for cells categorized to the same hypothalamic cell-type. Unsupervised clustering on the data reveals 25 groups of MEGs and PEGs with similar cellular expression patterns. The data reveal all imprinted genes are expressed in both neuronal and non-neuronal hypothalamic cells. The imprinted gene group containing *Ddc* is highlighted in bold on the y-axis (right side). The cell-types with putative *Ddc* imprinting effects are highlighted in bold on the x-axis (see Figure S3). Data from (Chen et al., 2017).

**Figure S2.**
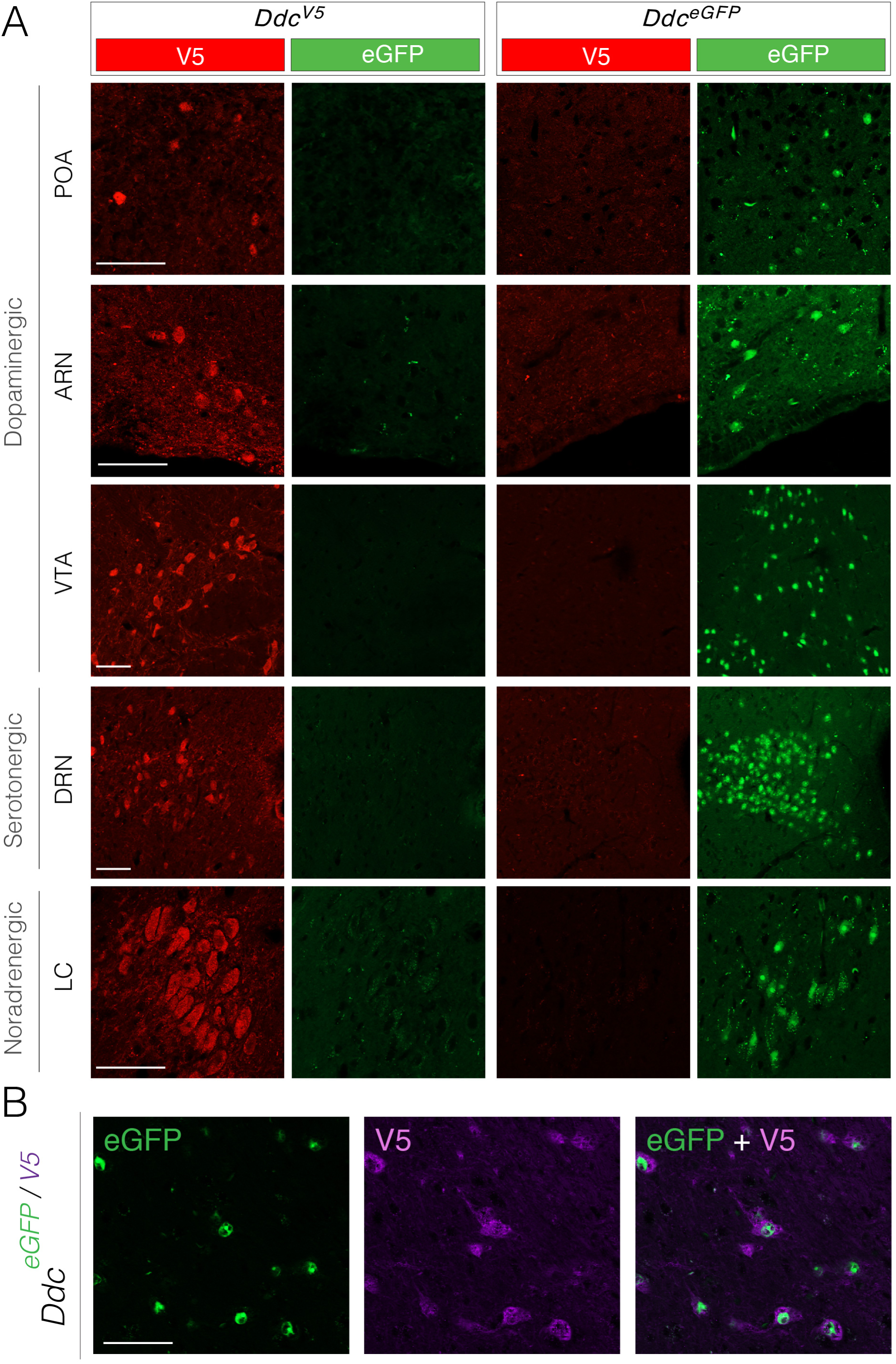
**Allelic reporter mice label major monoaminergic cell populations in the brain.** Related to Figure 2. (A) Images show expression of the V5 tagged DDC protein in *Ddc^V5^* mice and nuclear eGFP protein expression in *Ddc^eGFP^* reporter mice for major monoaminergic brain nuclei in the adult brain. Dopaminergic regions shown include the preoptic area (POA), arcuate nucleus (ARN) and ventral tegmental area (VTA); serotonergic regions include the dorsal raphe nucleus (DRN); noradrenergic regions shown include the locus coeruleus (LC). Size bars are 50 µm. (B) Images show expression of the eGFP and V5 alleles in the VTA of compound *Ddc^eGFP/V5^* reporter adult mice generated by mating *Ddc^V5^* and *Ddc^eGFP^* reporter lines. Size bar is 50 µm.

**Figure S3.**
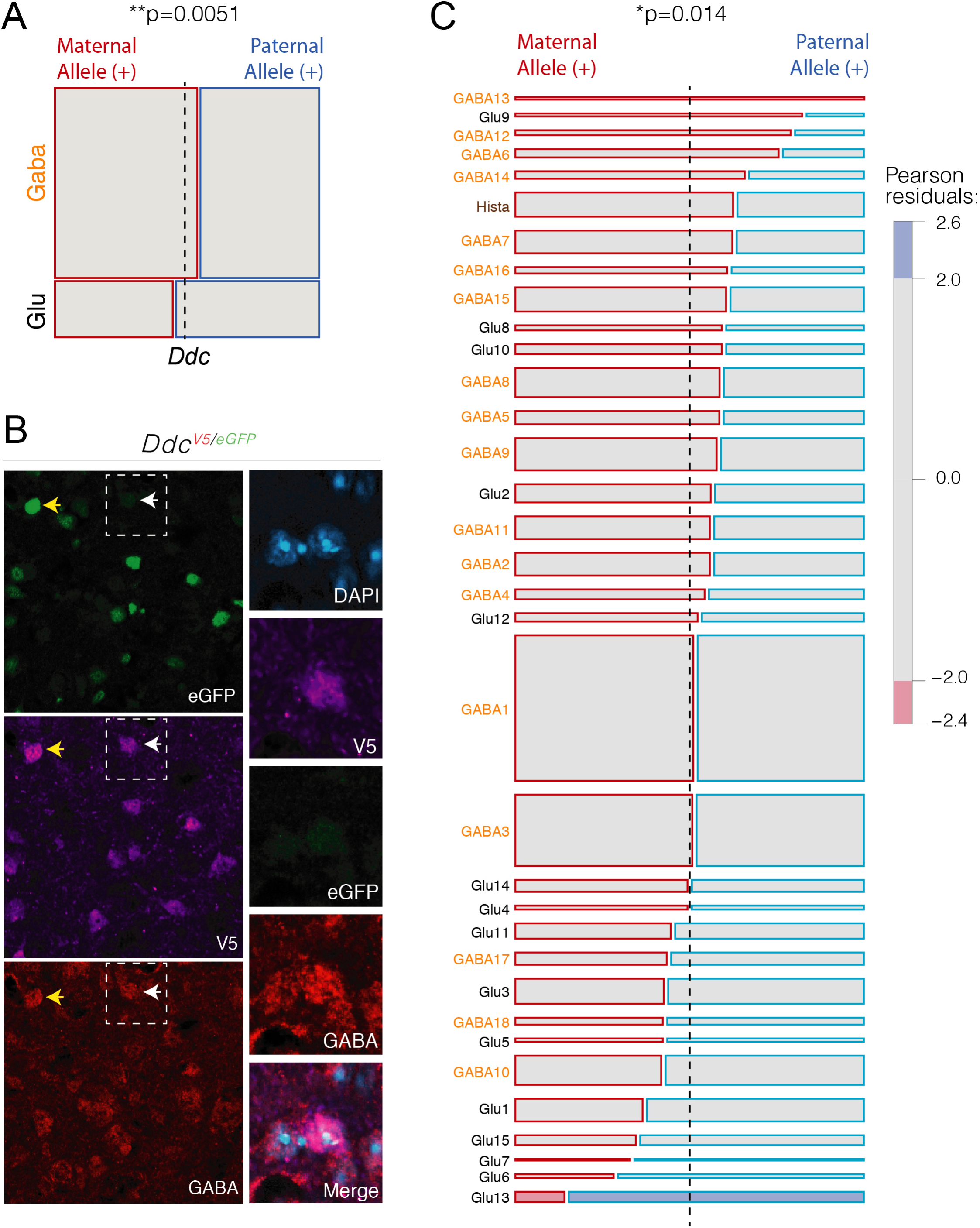
***Ddc* imprinting effects involving dominant expression of the maternal *Ddc* allele are more strongly associated with GABAergic compared to glutamatergic hypothalamic neurons.** (A) The mosaic plot shows the results of an IEN analysis for the correlation of *Ddc* imprinting effects in hypothalamus bulk RNA-Seq data (ARN region) to the expression of markers of GABAergic (Gaba) and Glutamatergic (Glu) hypothalamic neurons (See Figure S1F for method summary). The Chi-Square test results depicted in the mosaic plot reveal *Ddc* imprinting effects are significantly dependent on neuron type (p-value shown) and dominant maternal allele expression is positively associated with Gaba neurons. The Pearson residuals are plotted in the bars in the horizontal plane and the thickness of the bars is weighted by the number of marker genes for each neuron type. The dashed black line indicates the center point for the null hypothesis of no association effect. (B) Co-immunolabeling of GABA, eGFP and V5 in *Ddc^v5/egfp^* mice confirms DDC protein expression in GABAergic neurons in the hypothalamus and GABAergic neurons exhibiting dominant expression of the maternal *Ddc* allele. An example of a GABA+ neuron expressing both parental *Ddc* alleles equally is shown by the yellow arrow. An example of an imprinted GABA+ neuron with dominant expression of the maternal V5 tagged allele, but no expression of the paternal *Ddc* allele tagged with the stable nuclear eGFP reporter is shown by the white arrow. The images on the right are magnified images of the boxed region surrounding the imprinted cell. (C) The mosaic plot shows the results of an IEN analysis for the correlation of *Ddc* imprinting effects to markers of different subtypes of hypothalamic neurons. The Chi-Square test found significant dependence on cell-type (p-value shown). Dominant expression of the maternal *Ddc* allele is most positively associated with marker genes expressed in GABA13, Glu9, GABA12, GABA6 and GABA14 neurons, and others with red bars extending past the right side of the dashed black center point line. Maternal *Ddc* allele expression is negatively associated with markers expressed in several Glu neuron types (Glu13, Glu6, Glu7, Glu15 and Glu1) and other neuron types with red bars on left side of the dashed black center point line. Gaba neuron types (orange text) are relatively more positively associated with dominant maternal allele expression than Glu neuron types (black text).

**Figure S4.**
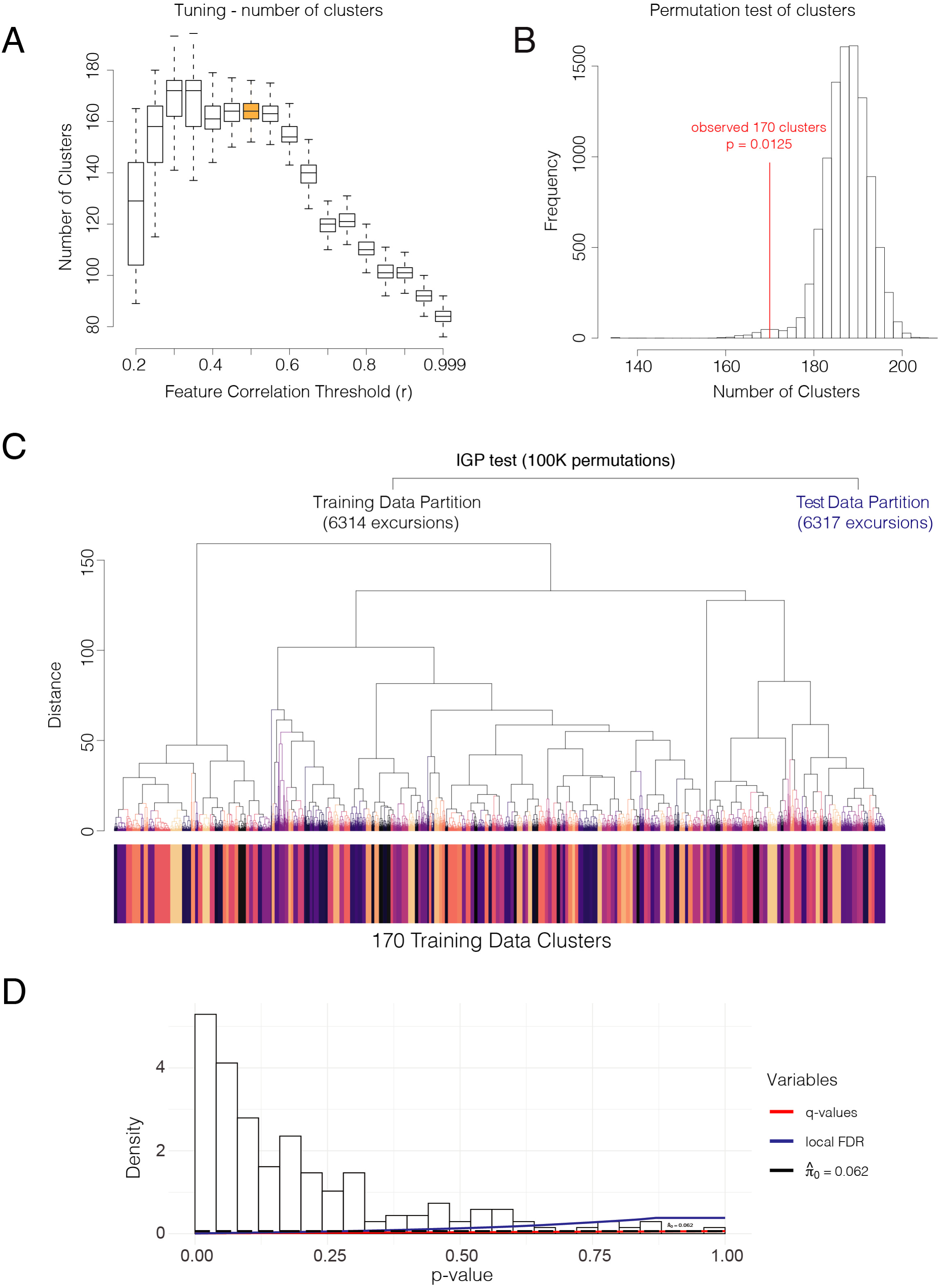
**Identification of modules of economic behavior from *Ddc ^−/+^*, *Ddc ^+/−^* and *Ddc^+/+^* adult male and female mice.** Related to Figure 5. (A) The boxplot shows the number of behavioral sequence clusters found at different correlation thresholds for 59 measures describing round trip excursions from the home. The calculation is performed 1000 times for each threshold. The data show that retaining measures correlated at r < 0.5 maximizes cluster detection and yields stable results. (B) The histogram shows the results of a permutation test of the clusters found using dynamic tree cutting from the 19 measures retained a correlation threshold of r <0.5. The number of clusters detected in the true data (red line) is significant compared to clusters identified from data in which the retained measures are randomly permuted, supporting bona fide clusters in the true data. The results of 10,000 permutations are shown. (C) The dendrogram shows the clusters identified in the genotype and sex balanced training partition of the behavioral sequence data by dynamic tree cutting (deep split =4; minimum cluster size = 20). The clusters are revealed from 6,314 excursions and the 19 measures retained at the correlation threshold of r<0.5. IGP testing was performed using the centroids from the training data clusters and the excursions in the test data partition. (D) The histogram shows the q-value analysis results of the IGP test p-values for each of the 170 training data clusters. At q<0.1, all 170 training data clusters are significant.

**Figure S5.**
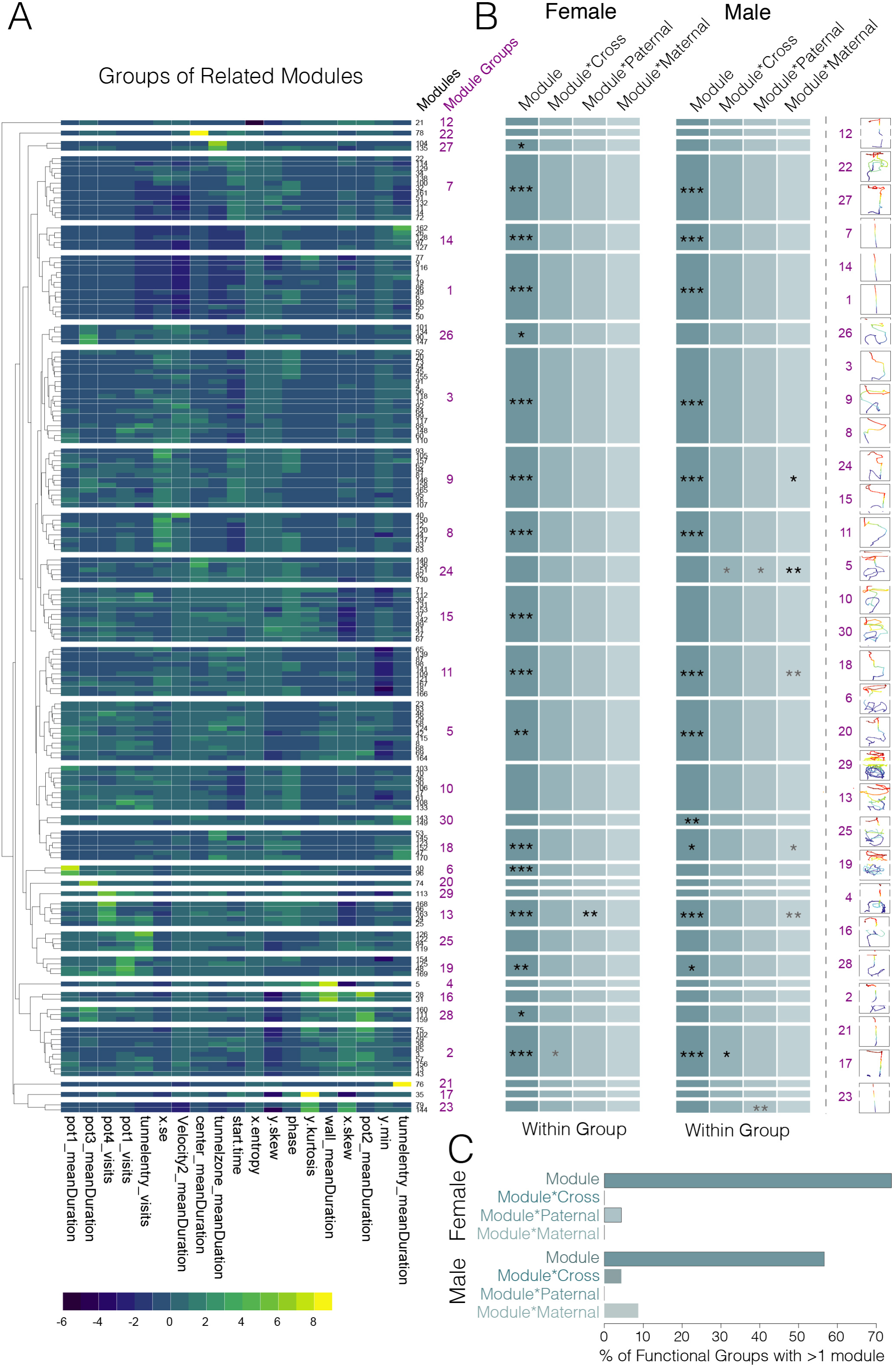
**Identification of module groups with parental effects on individual modules from Ddc −/+, Ddc +/− and Ddc+/+ adult male and female mice.** Related to Figure 6. (A) The heatmap is repeated from Figure 6 and shows the centroids for each of the 170 significant behavior modules (black type, right y-axis) clustered into 30 groups of related modules (purple type, right side y-axis). (B) The chart summarizes the results of the statistical modeling performed on each group of related modules and the results correspond to the groups in the heatmap (A). The significant main effect of module type (Module) and interactions between module type and cross (Module*Cross), paternal allele genotype (Module*Paternal) and maternal allele genotype (Module*Maternal) are shown in the grey columns. The stars indicating significance for interaction effects are black if the main effect is also significant and dark grey if the main effects are not significant. The colored traces on the far right show examples of the movement tracking patterns for modules in each grouping and show different groups involve different movement patterns. Movement traces are ordered according to the heatmap results. *p<0.05, **p<0.01, ***p<0.001. N = 24-30 animals per genotype and sex. (C) The bar plots show the percentage of module groups that have a main effect of module type and interaction effects within a group. Only significant interactions effects for which the main effect is also significant are shown. The color scheme matches the scheme of the chart (B).

## Supplemental Tables

**Table S1. Key to atlas of adult brain regions containing DDC+ neurons with dominant maternal allele expression.** Key to Figures 3 and Data S2

The table provides definitions for points plotted in Figs. 3C and S2. Abbreviated Region — A brain region(s) is labelled according to the online Allen Mouse Brain Reference Atlas version 1: coronal (https://mouse.brain-map.org/experiment/thumbnails/100142143?image_type=atlas). Images of DDC+ cell clusters captured by the 20X coronal field of view and grouped together for statistical analysis often contained more than one anatomically defined brain region in the Reference Atlas and are therefore named together with hyphenations. *Brain Region* — Definitions for the brain region(s) abbreviations. *Domain* — Major anatomical structure in which brain regions are located. *Ddc-Gfp/V5* images (n) — number of images from DdceGfp/V5 cross. Ddc-V5/Gfp images (n) — number of images from DdcV5/Gfp cross. GFP dominant cells Fisher’s (p) — p-value from Fisher’s exact test for cross difference where positive values indicate cell subpopulations with GFP dominant expression from the maternal allele, and negative values indicate cell sub populations with GFP dominant expression from the paternal allele. V5 dominant cells Fisher’s (p) — p-value from Fisher’s exact test for cross difference where positive values indicate cell subpopulations with V5 dominant expression from the maternal allele, and negative values indicate cell sub populations with V5 dominant expression from the paternal allele. Rostral Allen Atlas section — Coronal section number in the Reference Atlas at the rostral end of the region’s range. Caudal Allen Atlas section — Coronal section number in the Reference Atlas at the rostral end of the region’s range.

## Supplemental Data Legends

**Data S1. Imprinting∼expression correlation networks (IENs) for the identification of major imprinted cell-types for imprinted genes.** Related to Figure 1.

(A and B) These data confirm that our bulk RNA-Seq data yields the expected co-expression relationships for previously identified mouse-human conserved molecular markers of neurons, astrocytes and oligodendrocytes. Variance in the expression level of the neuron marker, *Ahi1,* across RNA-Seq replicates (n=18) reveals a positive correlation to another conserved neuron maker, *Syn2,* but not to the oligodendrocyte marker, *Mobp,* or the astrocyte marker, *Sox9* (A). For 1000 conserved neuron (N), oligodendrocyte (O) and astrocyte (A) marker genes, the mean co-expression correlation for all pairwise comparisons (Spearman Rho) is shown in the bar chart (B). The data reveal that genes expressed in the same cell-type are more positively correlated. Neurons and glia are most strongly differentiated.

(C and D) Scatterplots show that variance in the magnitude of imprinting effects (maternal allele expression – paternal allele expression) across bulk RNA-Seq replicates (n=18) reveals correlations to the expression levels of cellular marker genes. *Ube3a* is a MEG known to be imprinted in neurons and the data show that the *Ube3a* maternal-paternal allele difference is positively correlated to the expression of the neuron marker, *Ahi1* (C), as expected. *Peg3* is a PEG imprinted in neurons and the data show the *Peg3* maternal-paternal allele difference is negatively correlated to the expression level of *Ahi1*, as expected (D).

(E and F) Summary of our IEN testing methodology to define imprinting in neuronal versus non-neuronal brain cells. Published mouse-human conserved marker genes for neurons and non-neuronal cells were identified (E). The numbers of marker genes with expression levels that are positively or negatively correlated to the imprinting effects (maternal-paternal allele difference) of an imprinted gene of interest are computed (F). A Chi-Square then determines whether or not the imprinting effects are dependent on linkage to the cell-type markers and the Pearson residuals indicate whether maternal or paternal alleles are linked to a particular cell-type (neuronal versus non-neuronal).

(G and H) Mosaic plots show a demonstration of the IEN testing methodology for *Ube3a* (G) and *Peg3* (H). The results show the imprinting effect for each gene is significantly dependent on cell-type (p-value shown). The Pearson residuals reveal the type of cell that is associated with the preferentially expressed parental allele for the imprinted gene. The data show that the expression of the maternal *Ube3a* allele is positively associated (yellow) with neurons compared to non-neuronal cells (purple indicates a negative association) (G). In contrast, for *Peg3,* the paternal allele is positively associated with neurons (H).

**Data S2. An atlas of adult brain regions containing DDC+ neurons with dominant maternal allele expression.** Related to Figure 3.

The scatterplot shows the identity of brain regions containing DDC+ neurons with dominant maternal allele expression confirmed in reciprocal *Ddc^eGFP/V5^* and *Ddc^V5/eGFP^* crosses and those that do not. Most impacted regions are in the hypothalamus (see legend below). For each brain region, multiple images were captured and blindly scored as having cells with dominant eGFP or V5 allele expressing cells. A Fisher’s test of a contingency table of the scored images for each region was performed to determine whether a significant number of images had maternal or paternal allele dominant cells or neither (see Methods). Brain regions with significant maternal dominance are shown (dashed lines show p<0.05 threshold). Regions with cells having paternal allele dominance in both crosses were not observed. The data show 52 brain regions analyzed by 5 or more optical stacks per region per mouse (n=2, analyzed at 20X magnification). The brain regions are labeled based on Allen Brain Atlas annotations.

**Data S3. *Ddc* cross, maternal allele, and paternal allele effects manifest in offspring social behaviors.** Related to Figure 4.

(A-C) The plots show data for social behavior measurements with a significant cross X chamber interaction for sociability in females in a two-way mixed linear model (A), and main effect of cross in females (B) and males (J) in a nested model.

(D-G) The plots show data for female social behavior measurements with a significant main effect of maternal allele genotype (+ versus -) after absorbing cross effects. Movement patterns in females during the sociability (D and E) and social novelty (F and G) tests are affected, including the cumulative distance traveled (D and F) and mean time in the conspecific (E) and stranger (G) chambers. (p-values shown, likelihood ratio test, mixed linear model, n=20-30). A significant effect of paternal allele genotype was not observed. In the plots, each dot indicates data for one mouse, the box plot shows the mean ± the standard error of the mean.

(H) The plot shows data for social novelty seeking with a significant main effect of paternal allele genotype in females. A significant effect of maternal allele genotype was not observed for this measure. (p-value shown, likelihood ratio test, mixed linear model, n=20-30).

**Data S4. Exploratory behaviors in the open field and light-dark box tests reveal cross and imprinting effects, respectively, in offspring.** Related to Figure 7.

(A and B) The plots show data for measures with a significant main effect of the cross in the open field test for males (p-values shown, likelihood ratio test, mixed linear model, n=20-30). Significant effects were not observed in females.

(C and D) The plots show data for measures with a significant main effect of the maternal (C) or paternal (D) *Ddc* allele genotype in the light-dark test behavioral tests for females (p-values shown, likelihood ratio test, mixed linear model, n=20-30). Significant effects were not observed in males.

(E) The plots shown body weight data for adult males and females. Significant cross or imprinting effects were not observed (n>30).

## STAR METHODS

### RESOURCE AVAILABILITY

#### Lead Contact

- Further information and requests for resources and reagents should be directed to and will be fulfilled by the Lead Contact, Christopher Gregg.

#### Materials Availability

- Plasmids generated in this study are available from the Gregg lab and will be deposited into AddGene in the future with peer-reviewed publication of the paper.
- Mouse lines generated in this study are available from the Gregg lab and will be deposited into JAX labs in the future with peer-reviewed publication of the paper.

#### Data and Code Availability

- All code is available from the Gregg lab
- All data is available from the Gregg lab and will be deposited into public repositories in the future with peer-reviewed publication of the paper.

### EXPERIMENTAL MODEL AND SUBJECT DETAILS

#### Mice

##### Housing and husbandry

All experiments were conducted in compliance with protocols approved by the University of Utah institutional animal care and use committee (IACUC). F1 hybrid and *Ddc* Allele-Tag mice (see below) were bred and housed on ventilated racks at the University of Utah Comparative Medicine Center on a 12hr light cycle, 6am on and 6pm off; *Ddc Het* (see below) mice were bred and housed on static racks in the Biopolymers Building near the lab’s behavioral testing room on a 12hr reversed light, 11pm on and 11am off. All mice were given water and food (Harlan-Teklad 2920X soy protein-free) *ad libitum*, with the exception of a single overnight fast for mice tested for foraging behavior (see METHOD DETAILS, *Behavior*, foraging). Adult breeders (6weeks to 1year of age) were paired continuously, and pups were weaned at postnatal day 21 (P21) and cohoused with up to five same-sex littermates or similar aged same-sex mice of the same line; mice were never singly housed. As needed, ear punches were taken at P7 (*Ddc* Hets) or P17-P21 for both genotyping biopsy samples and mouse identification purposes. Before dissections of brain and embryonic heart tissue, mice were put to sleep with isoflurane gas anesthesia and decapitated.

##### B6CASTF1/J (F1bc) and CASTB6F1/J (F1cb)

Reciprocal F1 hybrid offspring were produced from C57Bl/6J dam X CAST/EiJ sire (F1bc offspring) and CAST dam X C57Bl/6J sire (F1cb offspring) mating crosses. Whole hypothalamic tissue was dissected from adult females for dissociation, purification of neurons and non-neurons, and RNA isolation.

##### Ddc Allele-Tag mice

Ddc-6His-P2A-eGFP-3xNLS (*Ddc^eGFP^* line) and Ddc-V5-P2A-mRuby2-3xNLS (*Ddc^V5^*) line constructs were designed and assembled by the Gregg lab (see Allele-Tag Construction in METHOD DETAILS), and the University of Nebraska Medical Center Mouse Genome Engineering Core Facility (Omaha, NE) used these constructs to perform CRISPR mediated homology directed repair for targeted insertion of the reporters immediately before the stop codon of the gene dopa decarboxylase (GRCm38/mm10; chr11:11815230) into C57BL/6J mice (see Easi-CRISPR Targeted Mutagenesis in METHOD DETAILS). Targeted insertion into the genome of F0 founder mice was confirmed by PCR amplifications using primers that flanked the 5’ and 3’ genome integration sites, and sanger sequencing confirmed that the reporter was in-frame with the *Ddc* coding sequence, contained no indels, and no non-synonymous amino-acid substitutions.

F0 founder mice were shipped to the University of Utah where they were backcrossed into the C57Bl/6J background strain for five generations to reduce propagation of any potential unknown off-target CRISPR mutations before producing homozygous reporter lines. We found that the mRuby reporter was not detectable and directly labeled for the V5 protein tag to detect expression in the line referred to as *Ddc^V5^*. Reciprocal crosses of *Ddc^eGFP^* dams X *Ddc^V5^* sires and *Ddc^V5^* dams X *Ddc^eGFP^* sires produced *Ddc^eGFP/V5^* and *Ddc^V5/eGFP^* offspring, respectively, for microscopy studies. All brains and adrenal glands from reporter mice were collected from adult females (Atlas, P65; AVPV P79-197), and expression of the reporters was restricted to brain regions with known monoaminergic cell populations. Embryonic heart was collected from both sexes between E16-18.

##### *Ddc* heterozygote mutants

Germline heterozygous *Ddc* mutant mice were made by crossing CMV-cre (Jax, Stock No: 006054) X *Aadc^flox7^* lines (Zhang et al., 2011). *Aadc^flox7^* mice have loxP recombination sites flanking exon-7 of the *Ddc* gene; this line was rederived at the University of Utah Transgenic Gene-Targeting Mouse Facility by *in vitro* fertilization (IVF) from cryopreserved sperm donated by the lab of Raymond C. Harris. (Vanderbilt University School of Medicine). Exon-7 CRE-recombinant excision was confirmed by PCR genotyping and Sanger Sequencing, and the resulting heterozygous *Ddc^Δ7^* line was backcrossed for 10+ generations into the C57Bl/6J background strain with the CMV-cre transgene removed. A battery of behavioral tests (see *behavior* section of METHOD DETAILS below) of both male and female mice of all genotype groups began between 8-10 weeks of age (P54-75) and lasted for five weeks; one task per week in the same order.

##### Genotyping

Ear punches taken at P7-P21 were lysed in 75µL of 25mM NaOH + 0.25mM EDTA with a 1 hour incubation in a thermalcycler at 98°C. Lysates were then pH neutralized with an equal volume of 40mM Tris.HCl, pH5.5. Two µL of lysates were then added to make 20µL PCR reactions with DreamTaq Green Master Mix (ThermoFisher, K1081) and 0.5µM primers (Table: Genotyping Primers).

**Table:**
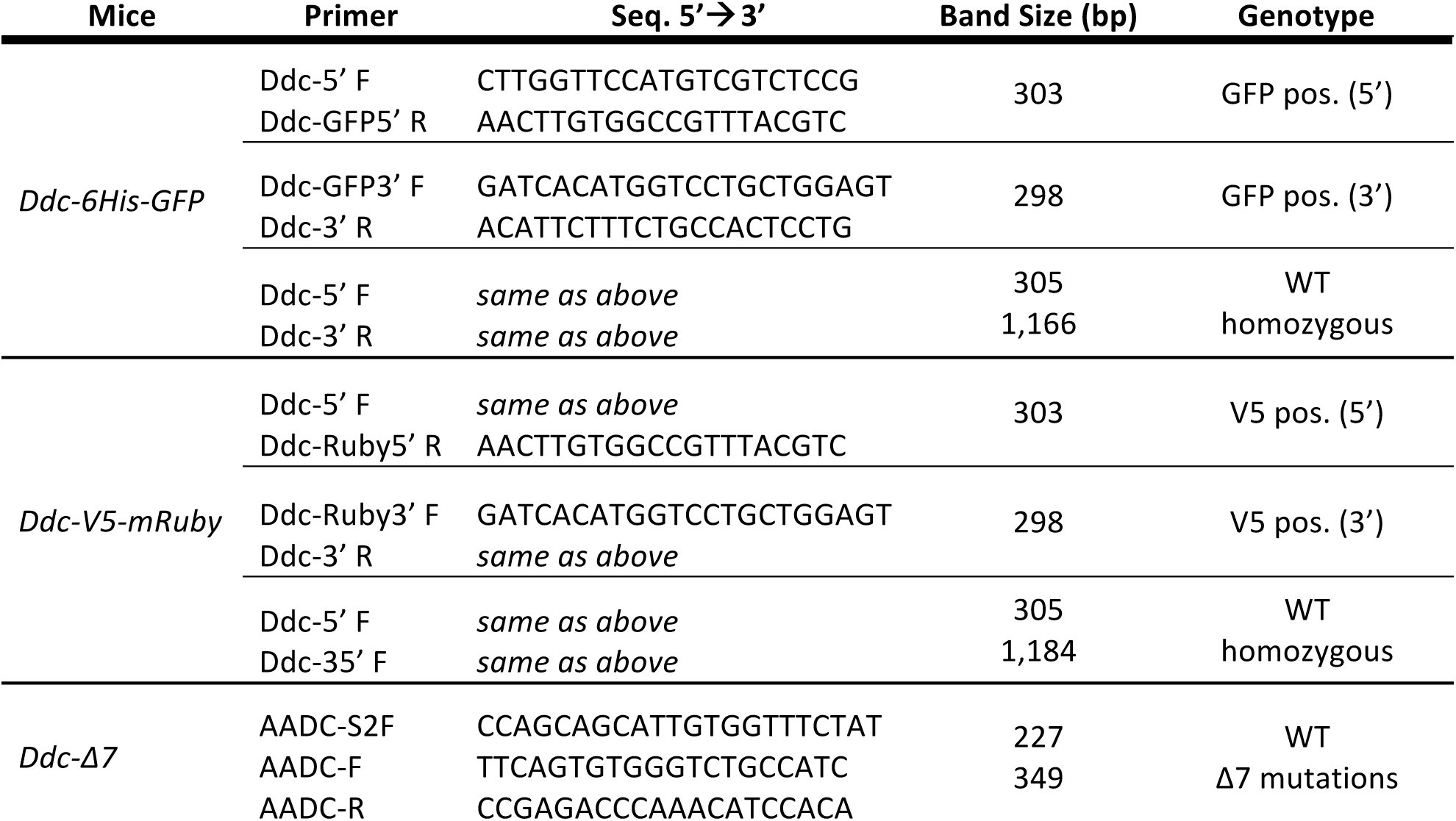
Genotyping Primers

### METHOD DETAILS

#### Pyrosequencing validation of cell-type specific imprinting

Adult female F1bc and F1cb reciprocal hybrid mice (see above) were euthanized, brains were removed from the skull and washed in cold Dulbecco’s phosphate buffered saline (D-PBS with calcium, magnesium glucose, and pyruvate) and whole hypothalami were dissected. Five hypothalami were pooled per sample and disassociated into single-cell suspensions using the the gentleMACS Octo Dissociator with heaters (Miltenyi Biotec, #130-090-427), gentleMACS C Tubes (Miltenyi, #130-093-237), and Adult Brain Dissociation Kit for mouse and rat (Miltenyi, #130-107-677) according to the kit protocol. After the red blood cell removal step, dissociated brain cells were pelleted and resuspended in 1ml D-PBS with Ca2+ and Mg2+ 0.5% bovine serum albumin (BSA). Cells were then purified into neuron and non-neuron fractions by magnetic separation using the Neuron Isolation Kit for mouse (Miltenyi, 130-115-389) according to the manufacturer’s instructions. Briefly, non-neuronal cells were first magnetically labelled by incubating cell suspension with Non-Neuron Cell Biotin-Antibody Cocktail (mouse) and Anti-Biotin MicroBeads. The entire cell suspensions (neurons and magnetically labelled non-neurons) were then run over MACS LS columns (Miltenyi, #130-042-401) attached to a MidiMACS Separator magnet. The magnetically labelled non-neuronal cell fraction was retained in the LS Columns in the magnetic field of the magnet, while the neuronal fraction was collected in the flow-through. The non-neuronal cells were then collected by removing the column from the magnet, adding buffer to the column, and pushing out with the plunger supplied with the column into a collection tube. Following column separation, neuronal and non-neuronal cell fractions were pelleted in a 1.5ml microcentrifuge tubes. Cell lysis and RNA isolation were performed by using the Invitrogen PureLink RNA micro kit according to the manufacturer’s instructions (ThermoFisher, 12183016). RNA was the converted into cDNA using qScript cDNA supermix (QuantaBio, 95048) according to manufacturer’s methods.

Pyrosequencing Allelic Quantification (AQ) analysis measured the ratio of maternal to paternal allele expression of imprinted genes from neuron and non-neuron cell fractions isolated from F1cb and F1bc mice using the Pyromark Q24 System (Qiagen, 9001514) according to manufacturer protocols and our published methods (Bonthuis et al., 2015). PyroMark Assay Design Software designed amplification and sequencing primers (Table: Pyrosequencing AQ Assay Primers) for AQ assays that contain strain distinguishing SNPs (Cast vs B6) within the genes. Four to six samples per strain (F1cb and F1bc) and cell-type fraction (neuron vs. non-neuron), four groups in total, were PCR amplified to measure allelic expression for each gene. An interaction effect between strain (F1cb and F1bc) and cell type (neuron and non-neuron) in two-way ANOVAs validated cell-type specific imprinting effects in the brain.

**Table:**
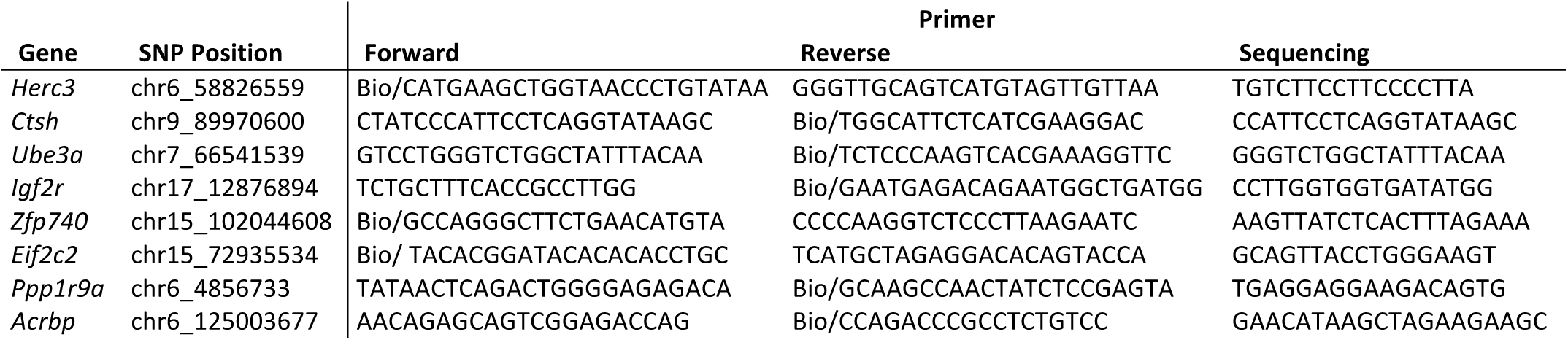
Pyrosequencing AQ Assay Primers

#### Allele-Tag reporter mice

##### Recombinant DNA construction

Dopa decarboxycalse (*Ddc*) allele-tag plasmid constructs for knockin to the C57Bl/6J genome were made using the Gibson Assembly method to join *Ddc* homology arms and reporter constructs into the pCRII-TOPO vector. Two custom reporter constructs were designed and synthesized as 1000ng of ∼1kb gBlocks from Integrated DNA Technologies: Ddc40HA-6His-P2A-eGFP-3xNLS and Ddc40HA-V5-P2A-mRuby2-3xNLS. Ddc40HA-6His-P2A-eGFP-3xNLS consists of the last 40bp of DDC coding sequence c-terminally conjugated with a 6His epitope tag, P2A self-cleaving peptide sequence, eGFP conjugated with three c-terminal copies of a nuclear localization sequence (3xNLS), stop codon, 37bp of mutated *Ddc* 3’-UTR (mUTR) to prevent homology repair between the stop codon and a CRISPR cut site 34bp downstream (preserving the 3’-splice junction of exon 14), followed by 40bp of un-mutated *Ddc* intron 14 sequence. Ddc40HA-V5-P2A-mRuby2-3xNLS was similarly designed except the *Ddc* c-terminal coding sequence is conjugated with a V5 epitope tag, and the fluorescent reporter is mRuby2-3xNLS.

1000ng gBlock constructs were diluted in TE to 10ng/ul. *Ddc* homology arms with approximately 1kb sequence to the left (LHA) and to the right (RHA) of the *Ddc* stop codon were PCR amplified using genomic DNA template isolated from C57Bl/6J by Phenol/Chloroform extraction and ethanol precipitation methods. The 3’-end of the LHA product (Primers: LHA F1, 5’-atctgtccaaggccaagagc-3’, LHA, R1 5’-ttctttctctgccctcagcac-3’) ends 1bp upstream of the stop codon end extends ∼1kb in the 5’-direction. The 5’-end of the RHA product (RHA F1 5’-acatctgtttccttgtggaggc-3‘, RHA R1 5’-gaccaaagactgccctggaa-3’) begins 40bp downstream of the stop codon and extends ∼1kb in the 3’-direction. For assembly, the entire linear pCRII-TOPO vector (Invitrogen **K460001)** was PCR modified using primers that anneal to the 3’-ends of the open multiple-cloning site, and contain a 40bp 5’-overhang with either homology to 3’-end of the RHA (PCRIITopo_DdcRHA_F1) or to the 5’-end of the LHA (PCRIITopo_DdcLHA_R1). LHA, RHA, and modified plasmid were PCR amplified using Phusion HF polymerase (NEB M0531S), and purified using EZNA Cycle Pure columns (Omega BiotTek, D6492-01) according to manufacturers’ protocols. In separate 20µL assembly reactions, 75ng (∼0.12 pmol) of Ddc40HA-6His-P2A-eGFP-3xNLS or Ddc40HA-V5-P2A-mRuby2-3xNLS allelic reporter constructs were stitched together with 75ng (∼0.12 pmol) of each homology arm (LHA and RHA) into 100ng (∼0.04 pmol) of the overhang modified pCRII-TOPO vector in Gibson Assembly Master Mix (NEB, E2611) for 1hr at 50°C. The Gibson assemblies were then diluted 1:4 in nuclease free water and transformed into One Shot TOP10 (Invitrogen, C404003) competent E. coli cells, plated onto ampicillin selective LB agar plates (supplemented with IPTG and XGAL for blue-white selection), and transformed colonies were picked and grown in 3ml LB cultures to purify plasmid DNA with EZNA Plasmid Mini Kit (Omega BioTek, D6942-02). Construction of assembled plasmids (pCRII-TOPO-Ddc-6His-eGFP-3xNLS-mUTR, and pCRII-TOPO-Ddc-V5-mRuby2-3xNLS-mUTR) was confirmed by sanger sequencing.

##### Easi-CRISPR Targeted Mutagenesis

*Ddc* Allele-Tag mice were made using the *Easi-CRISPR* methodology at the University of Nebraska, Transgenic Core Facility (Quadros et al., 2017). Briefly, allele-Tag plasmid constructs (pCRII-TOPO-Ddc-6His-eGFP-3xNLS-mUTR, and pCRII-TOPO-Ddc-V5-mRuby2-3xNLS-mUTR) were used as template for PCR amplification using GoTaq Long PCR Master Mix (Promega, M4021) and ultramer synthetic oligonucleotides (DDC-GFP-IVTRT F and DDC-IVTRT R, and DDC-mRuby-IVTRT F and DDC-IVTRT R) as primers to make cassettes for use in single-stranded DNA (ssDNA) synthesis by *in vitro* transcription and reverse transcription (IVTRT). PCR products were purified using Wizard SV Gel PCR Clean up System kit (Promega A9282). From 5’ to 3’, the cassettes contained 12 buffer nucleotides (atatcggatccc), a T7 transcriptional promoter (TAATACGACTCACTATAG), 78bp of *Ddc* LHA upstream of the stop codon, the Allele-Tag reporters (either 6His-P2A-eGFP-3xNLS, and V5-P2A-mRuby2-3xNLS), *Ddc* stop codon, 12bp of mutated 3’UTR to eliminate Crisper RNA (CrRNA) protospacer sequence to prevent re-cutting after homology directed repair, followed by 58bp of RHA sequence starting 13 nucleotides downstream of the end of the stop codon. RNA was synthesized by *in vitro* transcription using 1.5-2.5ng of DNA cassettes and mMESSAGE mMACHINE T7 Ultra Kit (Ambion, AM1354) according to manufacturer’s instructions by incubating overnight at 37°C, followed by adding 1µl of TURBO DNAse at 37°C for 15 min., and purified with MEGAclear Kit (Ambion, AM1908) according to manufacturer’s instructions. Then cDNA was synthesized using 3-5ug of RNA and M-MuLV Reverse Transcriptase (NEB M0253S) in 30µl reaction according to manufacturer’s protocol with 90 min. 42°C incubation and 5min. 80°C inactivation. RNA was removed by adding 3µl of RNAseH and incubating at 37°C for 30min. Resulting, ssDNAs were gel purified by running on 1% low melting agarose electrophoresis at 135V for 30min., staining with EtBr and excising ssDNA bands, and extracting from gel slices with Wizard SV columns using 25µl of prewarmed injection buffer for the first and second elution. Injection mix was assembled by mixing crRNA and trRNA (IDT, CAT# 1072534) at 1:2 ratio and annealing in a thermalcycler. 1µl of annealed gRNA (crRNA:trRNA) and 0.6µl of Cas9 protein (3.3ug/µl Final Concentration) was mixed into injection buffer and incubate at room temp. for 20-30 min. to form ribonucleoprotein (RNP) complexes. Final volume of injection mix is 50µl with final concentrations of 20ng/µl Cas9 protein and 10ng/µl gRNA. Added ssDNA to 10ng/µl in injection mix and filter through Millipore Centrifugal Filter units (UFC30VV25) by centrifugation at 13.5rpm for 5min. at room temp. The RNPs and ssDNA were then microinjected into single-cell embryos to cut the genomic DNA approximately 17bp downstream of the stop codon, and the ssDNA could be used as template for homology directed repair (HDR) and insertion into the genome at the *Ddc* locus. Injected embryos were then grown in vitro and implanted into pseudo pregnant females. F0 pups were screened for targeted insertion of the reporter construct by PCR and sanger sequencing across the 5’ and 3’ ends of the RHA and LHA to ensure integration into the genome.

#### Histology

##### Tissue preparation

Under deep isoflurane anesthesia, adult reporter mice were transcardially perfused with PBS, to clear blood, followed by ∼25-50ml of 4% paraformaldehyde to fix tissues. After perfusion, brains were dissected from the skull and adrenals from the abdomen, post-fixed in 5ml of 4% paraformaldehyde for ∼48hrs at 4°C, cryoprotected by immersing serially in 15% and then 30% sucrose at 4°C until the tissues sank, embedded and frozen into OCT in cryomolds using a methanol dry-ice bath, and stored at −80°C until sectioning. Coronal brain sections from regions of interest were then cut on a cryostat at 20µm thickness to directly mount onto Superfrost Plus slides (Fisher, #12-550-15), air-dried, and stored in a slide box at −80°C until Nissl stained or immunolabelled. Brain and adrenal tissue labelled as floating sections were processed as above except they were cut at 30µm thickness, collected into vials of antifreeze solution (300g sucrose, 10g polyvinylpyrrolidone 300ml ethylene glycol, and 0.1M phosphate buffer to 1L) and stored at −20°C until immunolabeling.

For embryonic heart tissue, pregnant mothers were euthanized under isoflurane anesthesia before dissection out the uteri containing embryos into ice-cold PBS in a petri dish. Embryos were then dissected from the uterus and extraembryonic tissue and transferred to a clean dish with fresh PBS. Embryonic hearts were micro-dissected under a stereo microscope, immersed in 4% paraformaldehyde for ∼48hrs at 4°C, exchanged with 15% and 30% sucrose, frozen in OCT molds made from aluminum foil, and stored at −80°C. 10µm embryonic heart sections were cut on a cryostat, mounted onto Superfrost Plus slides, and stored in a slide box at −80°C until staining.

##### Nissl Stain

For the brain Atlas of *Ddc* maternal dominant expressing cells, adjacent parallel sections were Nissl stained to identify neuroanatomical landmarks according to the Allen Mouse Brain Reference Atlas. Slides with 20µm sections were immediately submerged in 1:1 ethanol/chloroform overnight, then rehydrated through 100%, 95%, 70%, 50% ethanol to distilled water. Sections were then stained for 10 min. in prewarmed 0.1% cresyl violet made fresh with 0.3%v/v glacial acetic acid in a 45°C oven. Slides were then quickly rinsed in distilled water and differentiated for 2-30 min. in 95% ethanol while checking microscopically for best result. Sections were then dehydrated in 100% ethanol 2 × 5min., cleared in xylenes 2 × 5min, and cover-slipped with EcoMount (Biocare Medical, EM897L) mounting medium.

##### Immunolabelling

Slides with 20µm brain sections (Figs. 2I, S2, 3) were removed from −80°C storage, air-dried and hydrophobic barrier pen was drawn around sections. By Immersing slides in Copland jars on an orbital shaker with gentle-agitation, sections were hydrated in PBS, permeabilized for 10min. in PBS + 0.2% triton-X 100 and washed 3 × 5 min. with PBS + 0.025% triton-X 100. In a humidified chamber sections were covered and blocked with 10% normal donkey serum (Lampire Biological Products #7332100) and 1% BSA in PBS for 2hr. at room temp., and then incubated overnight at 4°C with goat polyclonal anti-V5 primary antibody (abcam ab9137) diluted 1:1000 in primary buffer (1% BSA in PBS). The next day, sections were washed 3 × 5 min. with PBS + 0.025% Triton-X 100 in Copeland jars at room temp. with gentle agitation. Then, sections were covered with donkey anti-goat-568 or 647 (Invitrogen #A32816, #A-11057) diluted 1:250 in PBS (no triton) and incubated for 2 hr. at room temp. protected from light in a humidified chamber. Sections were then washed 3 × 5 minutes with PBS, and cover-slipped with Vectashield anti-fade mounting media with DAPI (Vector Labs, H-1200). Embryonic heart sections (Fig. 2B-G) were immunolabelled similarly except they were cut to 10µm thickness, blocked for 30 min. with 10% donkey serum (no BSA), primary buffer contained 10% donkey serum (no BSA), secondary was diluted 1:200, and sections were cover-slipped with ProlongGlass + DAPI (ThermoFisher, P36981).

Floating sections were transferred from vials to wells of a 12-well polystyrene microtiter plate for immunolabelling; plates were placed on an orbital shaker to gently agitate sections for all washes and incubations. To remove antifreeze, sections were fist washed with 6 × 5 min. PBS exchanges. Then, sections were blocked with 10% normal donkey serum + 0.3% Triton-X 100 for 1 hour at room temperature, and incubated with primary antibodies (chicken anti-GFP 1:2000, abcam ab13970; goat anti-V5 1:1000, abcam ab9137; rabbit anti-GABA 1:500, Sigma A2052) overnight at 4°C. The next day, sections were washed 3 × 20 min. with 0.1% Triton-X 100 in PBS exchanges. While protecting from light, sections were then incubated with secondary antibodies (donkey anti-chicken-488 1:200, Jackson Immuno 703-545-155; donkey anti-rabbit-568 1:200, Invitrogen A10042; donkey anti-goat-647 1:200, Invitrogen A-21447) in 1% normal donkey serum + 0.1% Triton-X 100 in PBS for 2 hrs. at room temp., washed for 3 × 20min. with 0.1% Triton-X 100 in PBS, and finally washed in 2 PBS exchanges. Using a paintbrush and 0.5X PBS, sections were float-mounted onto positively charged slides, and cover-slipped with ProlongGlass + DAPI mounting media.

##### DDC Brain Atlas

The entire fixed brains of *Ddc^GFP/V5^* and *Ddc^V5/GFP^* adult female mice were sectioned at 20µm thickness and mounted onto positive charged slides in five parallel series containing representative coronal sections along the entire rostral caudal axis. Every section from one series for both crosses was fluorescently immunolabled for V5 as described above to detect cells expressing the *Ddc^V5^* allele, while expression of the *Ddc^GFP^* allele was visualized by native GFP fluorescence concentrated in the nucleus. A second series with parallel sections to the immunolabelled series was Nissl stained. Epifluorescence images of DDC+ cell clusters were captured on a Zeiss AxioImager 2 as optical slices using ApoTome2 structured illumination, a 20X objective, fluorescence filters (DAPI, GFP/488, TexasRed/568, 647) and Zen2 software. Nissl stained sections were captured with brightfield illumination, a 5X objective, and multiple images capturing entire coronal sections were stitched together using a motorized stage and Zen2 tiling. The fluorescence images captured in the camera’s 20X field of view were compared to an adjacent, parallel Nissl stained section and the Allen Brain Reference Atlas to identify and label the anatomical location(s) of the DDC+ cells (Fig. 3A,B). Images were then assigned a random number and a researcher blind to cross and anatomical location manually scored whether the images contained GFP dominant, GFP biased, biallelic, V5 biased, and/or V5 dominant DDC expressing cells. Sections from AVPV of five *Ddc^GFP/V5^* and five *Ddc^V5/GFP^* adult female were prepared as for the Atlas and a researcher blind to genotype and cross counted the total number of all GFP and V5 biased and dominant allele expressing, and biallelic cells in every image of the region.

#### Behavior

Mice were tested once per week between 11am-6pm (dark phase) for five weeks in the following order of behavior tasks: foraging, week 1; open field, week 2; elevated zero maze, week 3; light-dark box, week 4; and social preference, week 5. Bedding was left unchanged for 5-8 days prior to each test. Mice were transferred in their home cages and habituated to the testing room with the door closed and ambient room lights off for at least 30min. prior to any test. All test arenas were placed on an elevated stage encircled by black curtains with four cameras, and infrared and white light (when needed) fixtures mounted on a pole high above (∼6ft) the stage. One, two, or four mice were tested at a time depending on test (see below). Mice movements were tracked with Ethovision XT 14 software (Noldus) under infrared illumination to collect behavioral data. Each arena was recorded by an individual camera that was centered and zoomed on the arena to fill the entire image space, focused on the subject, and aperture optimally opened to both enhance contrast of the black subject with background and to eliminate oversaturated glare from overhead light reflections. Subject identification numbers and independent variables of sex, age, and genotype were recorded into Ethovision with each test mouse. After testing, mice were housed into a new clean cage with a handful of soiled bedding from their previous home-cage. Between subjects, and before first trial, behavioral apparatuses were wiped-down with 70% ethanol to clean and remove odors, and males were always tested before females on any given day of testing. After Ethovision captures videos and tracks mouse movements, an investigator used the *Tracking Editor* to correct missed samples and tracking errors.

##### Foraging

Foraging behavior testing was performed as detailed previously (Stacher Hörndli et al., 2019). The detailed protocol published in the supplementary material for this previous study was followed.

##### Open field

Two 40×40cm open field arenas enclosed by 40cm high walls, made from white melamine laminated (impervious to liquids) pressboard, were placed adjacent to each other on the testing room stage. Arenas were illuminated to 170lux with two track-lighting, bendable gooseneck LED fixtures positioned to minimize both shadows and glare from light-source reflections.

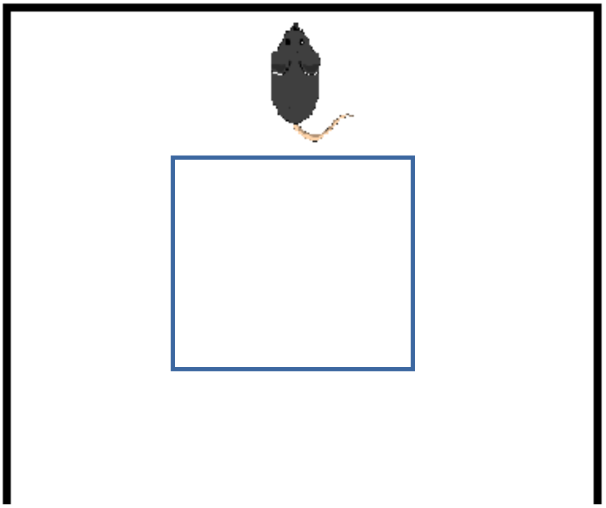

Ethovision *Experimental Settings* used two cameras at a resolution of 800 × 600 with a merged image of both arenas and center-point detection of subject. *Detection Settings* used “automatic setup” with a C57Bl/6J mouse exploring the arena, settings were adjusted so that missed samples were minimized (i.e to zero), and different test days used duplicated detection settings with an updated image capture of the arena background (arenas without a subject). Using a background image *Arena Settings* were adjust to defined arena areas, zones, and calibrated distances to include a 12cm wide wall-zone perimeter surrounding a 16×16cm^2^ center-zone. New trials were defined for every two mice tested simultaneously. After starting a new trial recording in Ethovision, mice were grabbed by their tail from their homecage and gently place into the wall-zone of an open-field arena facing the wall furthest from the researcher, the curtains were closed around the stage, and the experimenter left the room. Trial control settings were set to start tracking the center-point of the mouse two seconds after the subject was detected in the arena, and end tracking after 20min. The number of fecal-boli left in the arena was recorded before cleaning the arenas and starting the next trial. After all subjects were tested, the Ethovision *Analysis Profile* was set to export the following measures: number of fecal boli, total distance moved (cm), maximum velocity (cm/s), cumulative duration moving (s), frequency of times moving (n), mean duration of each movement (s), latency to enter center-zone (s), cumulative duration in center-zone (s), frequency of entries into center zone (n), mean duration in center-zone per entry (s), and mean duration in wall-zone per entry (s).

##### Elevated Zero Maze

A single elevated zero-maze (Stoelting, 60190) for mice was placed on the test-room stage, and was recorded in the dark with infrared illumination and a single camera mounted above. The zero maze consisted of a circular platform with a 50cm inner-diameter and outer-diameter, that is held above the stage with leg supports. The maze is divided into equal quadrant sectors: two closed sectors on opposite sides of the maze enclosed by high inner and outer walls, and in-between closed sectors were two equally sized open sectors without walls. The inner walls were made with infrared transparent plastic to track the mouse with the camera through the walls. Ethovision *Experiment Settings* were set to 800 × 600 resolution with one arena and center-point detection. *Arena settings* with a background image of the maze (maze without subject) for each test day was used to define the arena area, open (open sectors) and closed (closed sectors) zone areas, and to calibrate distance by the inner diameter (50cm) of the circular platform. *Trial Control* was set to start tracking when the subject was detected in the maze for two seconds and stop tracking five minutes later. Before the first trial, *Detection Settings* were adjusted with “automatic setup” to minimize missed and subject not found samples while a C57Bl/6J mouse explored the maze; different test days used duplicated detection settings with an updated background image. Smoothing was set to “high”. A single mouse was tested in each *trial* added to the *Trial List*, which defines the subjects and their independent variables. Under *Acquistion*, a background image was first captured, and then after pushing the “start trial” button the subject mouse was lifted by the tail from its’ home-cage and gently place at the junction between an open and a closed sector facing away from the researcher and toward an open sector. The curtains were closed around the stage while the experimenter remained in the room during the 5 min. trial. After testing all subjects the Ethovision *Analysis Profile* was set to export the following measures: total distance moved (cm), maximum velocity (cm/s), cumulative duration in open areas (s), frequency of entries to an open area (n), mean duration in open area per entry (s), latency to enter closed area (s), and mean/minimum/maximum acceleration (cm/s^2^).

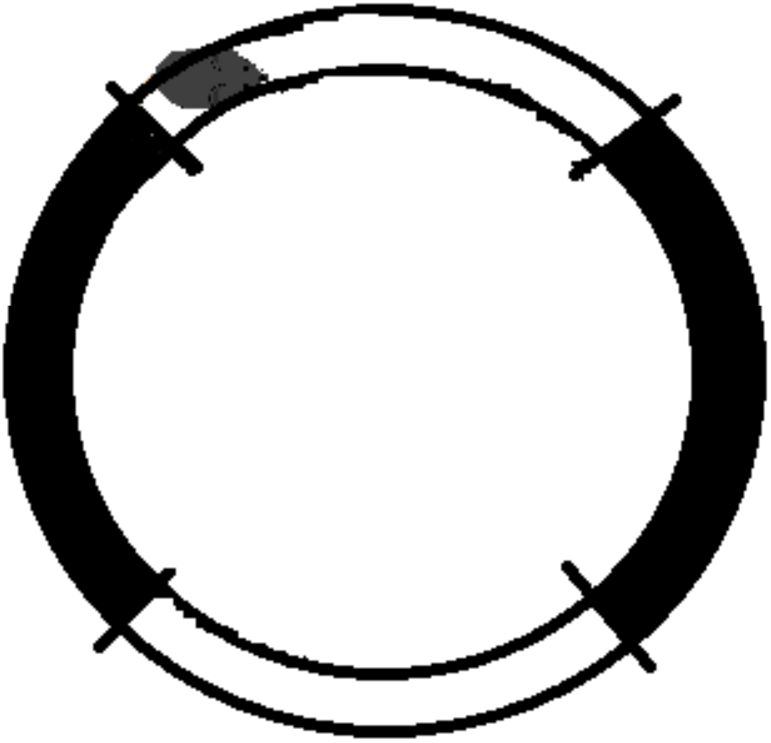

##### Light Dark Box

A single light-dark box (Stoelting 63101) was set on the testing-room stage underneath a camera, LED white lights, and infrared lights mounted above. The box was separated into light and dark chambers by a central divider with a door in the middle for the mouse to pass freely between the two sides. The dark chamber outer walls, lid, and central divider were made of plastic that blocked white light but transmitted infrared light for tracking mice in the dark with the infrared sensitive camera. The light chamber outer walls were made from transparent plastic and had no lid so that it could be illuminated from above by aversive white light (∼100-200 lux). Ethovision *Experiment Settings* were set to 800 × 600 resolution with one arena and center-point detection. *Arena settings* defined the arena area, the light and dark chamber zones, and calibrated distance. *Trial Control* was set to start tracking when the subject was detected in the maze for two seconds and stop tracking after 20 min. *Detection Settings* were adjusted with “automatic setup” to minimize missed and subject not found samples while a C57Bl/6J mouse explored the maze; different test days used duplicated detection settings with an updated background image. Smoothing was set to “high”. A single subject was tested in each trial of the *Trial List*, which defined the subjects and their independent variables. After capturing the background image and starting the trial under *Acquisition,* the subject mouse was lifted by the tail from its’ home-cage and gently placed into the center of the light side facing away from the researcher with the door to the dark side on the right side of the mouse. The curtains were then closed around the stage and the experimenter left the room for the 20 min. trial. After testing all subjects, the Ethovision *Analysis Profile* was set to export the following measures: total distance moved (cm), maximum velocity (cm/s), frequency of entries into the light side (n), cumulative duration in the light side (s), latency to enter the dark side (s), number of transitions from the dark side to the light side (n), and mean, minimum and maximum acceleration (cm/s^2^).

##### Social Preference and Social Novelty

A standard three-chambered box (Stoelting 60450), with center, left and right chambers separated by doors, was used to test social preference and social novelty behavior. In the week prior to testing, social stimulus conspecific adult C57Bl/6 males were habituated to being confined to the round jail-cells used in the three chambered box for 30 min. for 3-5 days. Tests were recorded in the dark with infrared illumination, and two mice were tested in separate mazes at a time. In the *social preference* task one chamber (left or right) was occupied with a conspecific confined to a jail-cell, while the opposite chamber (right or left) contained an empty jail-cell. To begin the social preference task, the test mouse was habituated to the center chamber for five minutes with the doors closed to the left and right chambers. The experimenter then started Ethovision recording and removed the doors between the chambers to allow the test mouse to investigate the chamber and jail-cell containing the conspecific, and the empty chamber and jail-cell, for 10 min. Immediately following the 10 min. social preference test, the test mouse was returned to the center chamber with the doors closed to begin the social novelty task. For the social novelty task, a novel “stranger” conspecific was placed into the previously unoccupied jail cell while the now “familiar” conspecific used in the preceding social preference task remained in its’ jail cell. The experimenter then removed the doors while Ethovision recorded the test mouse investigating the chambers and jail cells with either the “familiar” or “stranger” conspecific for an additional 10min. trail. Conspecific stimulus mice were not reused in consecutive tests, and use of the left or right chamber for the first/familiar and empty/stranger conspecifics was counterbalanced by sex and genotype. Ethovision *Analysis Profile* exported the following measures for the *social preference* task: total distance moved (cm); maximum velocity (cm/s); cumulative duration (s), frequency of entries (n), mean duration per entry (s), and latency to enter (s) the conspecific chamber; cumulative duration (s), frequency of entries (n), mean duration per entry (s), and latency to enter (s) the empty chamber; cumulative duration (s), frequency of entries (n), and mean duration per entry (s) in center chamber; cumulative duration (s), frequency (n), mean duration (s), and latency(s) for nose to investigate conspecific jail (s); cumulative duration (s), frequency (n), mean duration (s), and latency (s), for nose to investigate empty jail; and fecal boli (n). The same measures were exported for the social novelty task, except the chambers and jails contained either a “familiar” or “stranger” conspecific.

### QUANTIFICATION AND STATISTICAL ANALYSIS

#### Imprinting ∼ Expression Correlation Network Analysis

Published RNASeq maternal and paternal allele expression data and gene expression data for the arcuate nucleus region of the adult female hypothalamus for F1cb (n=9) and F1bc (n=9) mice was obtained (Bonthuis et al., 2015). For each biological replicate, the difference in the expression of the maternal and paternal allele was computed for each gene (maternal – paternal). The correlation of the maternal and paternal allele expression difference to the expression of published conserved human-mouse cell-type marker genes in the data was computed. The top 1000 most specific marker genes for mouse-human conserved markers of neurons, oligodendrocytes, astrocytes, endothelial cells and microglia were used (McKenzie et al., 2018). We then tallied the number of positively and negatively correlated marker genes for each category and a Chi-square test of the dependence of the allelic expression difference on the cell type category was computed. A mosaic plot was computed using the vcd package in R, which revealed the pearson residuals and associations between cell-types and parental allele expression.

#### Single Cell RNASeq Data Analysis

Published adult mouse hypothalamus single cell RNASeq data generated by (Chen et al., 2017) were downloaded from GEO (gene expression omnibus). We extracted the gene expression data for each cell assigned to each cell-type, according to definitions in the published study. The mean expression level for each gene was computed across all cells for a given type. We then extracted the data for all cell types and imprinted genes, based on imprinted genes identified in our previous study (Bonthuis et al., 2015). Unsupervised hierarchical clustering of imprinted gene cellular expression profiles was performed in R using Euclidean distance and the Ward.D2 method.

#### Pyrosequencing

For each AQ assay, the percent expression from C57Bl/6J and Castaneous alleles were converted to log ratios; log(B6/Cast). Data was then analysed by two-way ANOVA in R with *Cross* (levels: F1cb, F1bc) and *Cell-type* (levels: neuron, non-neuron), and a significant interaction effect *Cross X Cell-type* validated cell-type specific imprinting in the brain.

#### Brain Regions with DDC imprinted cell subpopulations

##### Atlas Fig. 3A-C

After determining whether each image exclusively contained subpopulations of cells with dominant GFP or dominant V5 allelic expression, or not, they were decoded for cross and brain region and organized into 52 groups according to brain region(s) for statistical analysis (n > 7 per region). Fisher’s Exact tests determined for each region whether there was a significant difference in the number of images from *Ddc^GFP/V5^* compared to *Ddc^V5/GFP^* containing GFP dominant cell subpopulations; the same was done for subpopulations of V5 dominant cells. Each region was then plotted with X- and Y-coordinates of Fig. 3C, with the X-axis representing the -log(p) value for a cross difference in GFP dominant subpopulations and on the Y-axis representing the -log(p) for a cross difference in V5 dominant subpopulations. On both axes the -log(p) values were plotted in the positive direction when allele dominant expression comes from the maternal allele (i.e. from GFP of the *Ddc^GFP/V5^* cross, and from V5 of the *Ddc^V5/GFP^* cross) and in the negative direction when allele dominant expression comes from the paternal allele (i.e. from GFP of the *Ddc^V5/GFP^* cross, and from V5 of the *Ddc^GFP/V5^* cross). Therefore, regions in the upper right coordinate of the plot represent regions that show dominant maternal allele expression from both crosses.

##### AVPV Fig. 3D

Counts of the number GFP dominant (GD), GFP biased (GB), biallelic (BA), V5 biased (RB), and V5 dominant (RD) cells from all images of the AVPV were calculated as percent of the total number of all DDC+ cells for each individual; n = 5 *Ddc^GFP/V5^* and n=5 *Ddc^V5/GFP^*. These data were analysed with a two-way ANOVA with *Cross* and *Cell-type* main factors in GraphPad Prism 5 software.

#### Behavior

##### Social Preference and Novelty

Significant impacts of maternal or paternal allele *Ddc* mutations on behavior in the three-chambered box social preference and novelty tasks were measured with linear mixed models in R using the lme4 package (Figs. 4, S6A). The difference in cumulative duration (in seconds) of time spent in the empty compared to conspecific chamber, or investigating the conspecific or empty jail, were first fit to the following linear mixed model: lmer(Time ∼ Genotype + Chamber + Genotype:Chamber + (1|Mouse)). In this model the main factors *Genotype* (levels = wt, het) and *Chamber* (levels = conspecific, empty) are fixed effects, and *Mouse* is a random intercept effect that accounts for each individual subject scored simultaneously on both levels of chamber. Contrasts for genotype and chamber in the model were set to “contr.sum” for effects coding. Every *Sex* (male and female) and *Cross* (maternal het, and paternal het) combination were analyzed separately; therefore, four analyses for each dependent variable consisting of either males from the maternal het cross, males from the paternal het cross, females from the maternal het cross, or females from the paternal het cross were performed. Analysis of Deviance tables with Type III Wald test statistics were then calculated using the Anova function within the car package to determine which factors and their interaction significantly reduce deviance in the models. Significant interaction effects of *Genotype* by *Chamber* indicates that a heterozygous maternal or paternal allele impacts social preference or novelty phenotypes relative to their wild-type, same-sex littermates. A test for cross effects (levels = het mother, het father) in the data were analyzed similarly with the following mixed effects model: lmer(Time ∼ Cross + Chamber + Cross:Chamber + (1|Mouse)) with data collapsed for levels of *Genotype* and measured separately for each *Sex* (Fig. S6A).

##### Behavioral Analyses with Cross Effects

Cross effects in the data were absorbed in a deep analysis of maternal and paternal allele effects on aspects of social behavior using generalized linear models; and also on analyses of Open Field, Elevated Zero Maze, and Light-Dark Box exploratory and anxiety related behaviors. Generalized linear models were fit using the *glm* function in R with factors of *Cross* (levels: maternal het, paternal het), *Paternal Allele* (levels: wt, het), and *Maternal Allele* (levels: wt, het); i.e. *glm(Variable ∼ Cross + PatAllele +MatAllele)*. The *glm* models used the following distributions depending on the type of data: gamma (link=”inverse) for latency data, quasipoisson (link=”log”) for over-dispersed count data, and gaussian (link=”identity”) for all other measures. The anova function in R was then used to generate Analysis of Deviances Tables with p-values to determine which terms significantly reduced deviance in the models.

##### Foraging Behavior

###### Excursion Data Capture

Our DeepFeats approach for analyzing modularity in foraging was performed as previously described (Stacher Hörndli et al., 2019) with some modifications and advances. In our study, mice were tracked with Noldus Ethovision software. Noldus settings were used to define regions of interest in the foraging arena and indicated when the mouse was in each area. To ensure the tracking is equivalent across different mice, a Procrustes transformation of the XY coordinates was performed to put every tracking file in the same coordinate space. The track coordinates were zero’d to the center of the tunnel to the home cage. We then generated custom code in R to parse the raw Noldus tracking files into discrete, round trip home base excursions from the home cage tunnel. Each excursion is assigned a unique ID key that we call the Concise Idiosyncratic Module Alignment Report (CIMAR) string key. It stores the coordinates of the excursion in the data and the CIMAR string includes metadata regarding the mouse number, excursion number, sex, age, genotype and phase. Next, custom code compares the CIMAR coordinates to the raw Noldus data files and constructs a new dataset that extracts 57 measures from the Noldus output, which we use to initially statistically describe each excursion. The 57 measures are presented in (Stacher Hörndli et al., 2019) and are designed to capture a relatively comprehensive array of different behavioral and locomotor parameters, as well as describe interactions with food and non-food containing patches and exposed regions in the environment. These measures consist of shape, frequency, order and location statistics of an animal’s X and Y movements, numbers of visits and time spent at different features in the arena, including food patches (Pots#2 and 4), non-food containing patches (Pots#1 and 3), the tunnel zone, wall zone and center zone of the arena and data describing locomotor patterns, including velocity, gait and distance traveled. The 57 measures for each excursion are scaled (normalized and centered across excursions) because they are in different units.

##### Identification of a Set of Behavioral Measures to Resolve Modularity in Excursions

This section details the methods to define the set of behavioral measures that best resolve candidate modules. A data matrix was constructed in which the rows are excursions performed by the mice, labeled by CIMAR keys, and the columns are the 57 behavioral measures. A correlation matrix was constructed from the data using the Pearson correlation statistic. The measures were then systematically filtered from the data as detailed in the main text based on different correlation thresholds using the “findCorrelation” function in the caret package in R. With this approach, the absolute values of pairwise correlations are considered. If two variables have a high correlation, the function looks at the mean absolute correlation of each variable and removes the variable with the largest mean absolute correlation. We systematically threshold the data in r = 0.5+ value increments as shown in the main text to identify the best set of measures to resolve clusters of excursions. At each threshold, the retained measures are used in an unsupervised clustering analysis to define clusters of excursions. We used the Ward.D2 minimum variance method implemented using the “hclust” function in R to perform the clustering and define compact, spherical clusters. We then statistically define discrete excursion clusters from the results using the Dynamic Tree Cut algorithm (Langfelder et al., 2008). This is a powerful approach because it is adaptive to the shape of the dendrogram compared to typical constant height cutoff methods and offers the following advantages: (1) identification of nested clusters; (2) suitable for automation; and (3) can combine the advantages of hierarchical clustering and partitioning around medoids, giving better detection of outliers. We detect clusters using the “hybrid” method and use the DeepSplit parameter set to 4 and the minimum cluster size set to 20. The total number of clusters detected is quantified at each correlation threshold. Conceptually, more relaxed correlation threshold cutoffs could reduce cluster detection by retaining redundant measures that mask important effects from other measures. On the other hand, thresholds that are too stringent could reduce cluster detection by pruning informative measures. Our objective is to identify the threshold that uncovers the most informative and sensitive set of measures for resolving different clusters of excursions, setting the stage for the discovery of potential modules.

##### Statistical Validation of Significant Clusters of Excursions Based on Retained Behavioral Measures

In our study, Dynamic Tree Cut will deeply cut branches in a dendrogram generating large numbers of small clusters if there are few bona fide relationships in the data. Thus, to test whether bona fide clusters of excursions exist in the data we implemented a random sampling procedure in R in which we randomly sample from the matrix of the retained behavioral measure data to break the relationships between the excursions and the measures. The sampled null data matrix is then subjected to the same clustering and quantification procedure to determine the number of clusters found by Dynamic Tree Cut. A null distribution is created from 10,000 iterations and compared to the observed number of clusters, which is expected to be significantly less than the null due to bona fide biological relationships between the excursions and set of retained measures. A lower tailed p value was computed to test this outcome. In a modification compared our previous study,

##### IGP Permutation Test to Identify Significant Modules of Behavior

To test whether reproducible modules of behavior exist in the data for the foraging excursions, we use the in-group proportion (IGP) statistical method for testing for reproducible clusters between two datasets (Kapp and Tibshirani, 2007). We built a modified version of this function for parallelized computing to speed the analysis for large number numbers of permutations. The excursion data for the mice is separated into a training data and test data partition for reproducibility testing. A balanced partition was generated according to genotype, sex and phase factors using the “createDataPartition” function in the caret package in R. Unsupervised hierarchical clustering was performed on the Training data partition excursions and clusters were defined using Dynamic Tree Cut. Next, the centroids for each training data cluster were computed as the mean values of the behavioral data for the excursions in the cluster. The training data centroids were then used to compute the IGP statistic for each training data cluster based on the test partition data, thereby evaluating the reproducibility of each cluster.

In a modification compared to our previous study (Stacher Hörndli et al., 2019), we created a custom IGP permutation test that is based on a distance calculation, rather than the correlation implementation in the clusterRepro R package. The distance IGP testing framework was written in C++ and speeds the permutation test by many fold and is a more accurate replication of the clustering parameters used in the test data. We used this approach to compute p values for each cluster to determine whether the IGP value is greater than chance. False positives due to multiple testing errors were controlled using the q-value method (Storey, 2002). Modules are thus defined as significantly reproducible training partition excursion clusters (q < 0.1). Each module detected was assigned an ID number and individual excursions in the data were annotated based on the module they match to. This approach facilitated quantifications of module expression frequency by the mice.

##### Generalized Linear Modeling and Likelihood Ratio Testing of Module Expression Frequency

To statistically evaluate the factors that significantly affect module expression frequency, we used generalized linear modeling functions implemented in R. The full and nested models tested are detailed in the text for each analysis. The generalized linear modeling is performed using a Poisson or quasipoisson distribution, as indicated in the text. We tested the goodness-of-fit of the full model with a chi-square test of the residual deviance and degrees of freedom in R. If the test result was not significant (p > 0.05), we concluded the model fit the data and proceeded based on Poisson-distributed errors. If the Poisson model was not a good fit, we used a quasipoisson model. The likelihood ratio test comparing the full and nested models was performed using the anova() function in R with the additional option test = “Chisq.” Our statistical framework is based on a primary and secondary hypothesis testing strategy to manage multiple testing (Bender and Lange, 2001), as indicated in the main text. Likelihood ratio testing for full versus nested models was performed to define significant main and interaction effects in the data. Individual mice were treated as random replicates of the parental and genetic factors. The following full model was used to test aggregated module expression data for males and females (Fig. 5): glm(expression ∼ cross + paternal allele genotype + maternal allele genotype). The following full model was used to test expression data for grouped modules with similar centroid patterns for males and females (Fig. 6 and S5): glm(expression ∼module + cross + paternal allele genotype + maternal allele genotype + module:cross + module:paternal allele genotype + module:maternal allele genotype)

### DATA AND CODE AVAILABILITY

All raw data and code are available from the Gregg lab. The raw data will be deposited in a public database following publication in a peer-reviewed journal.

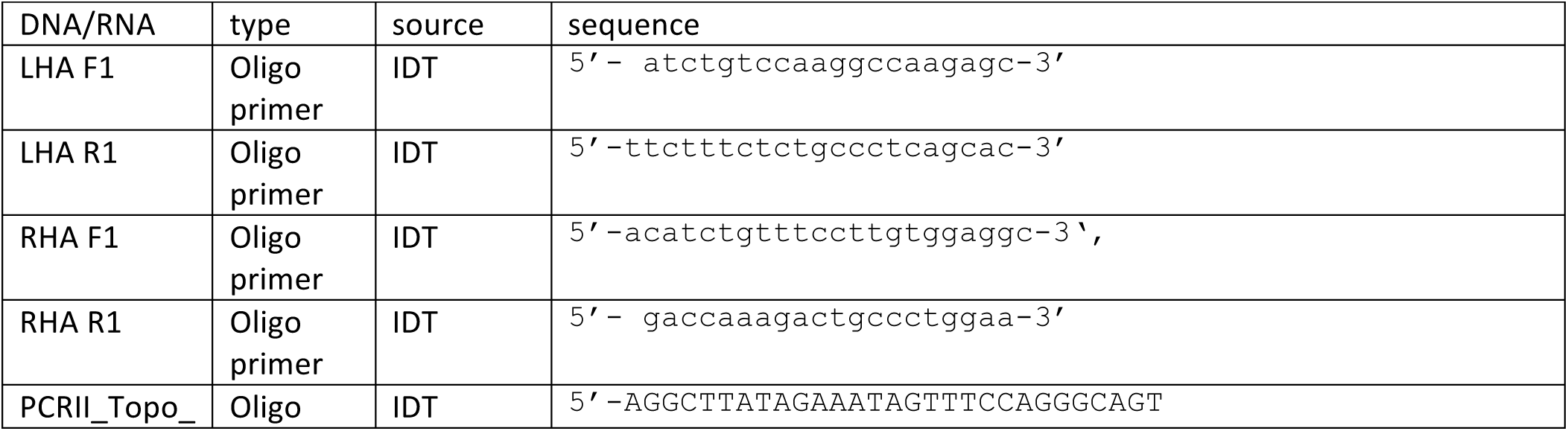
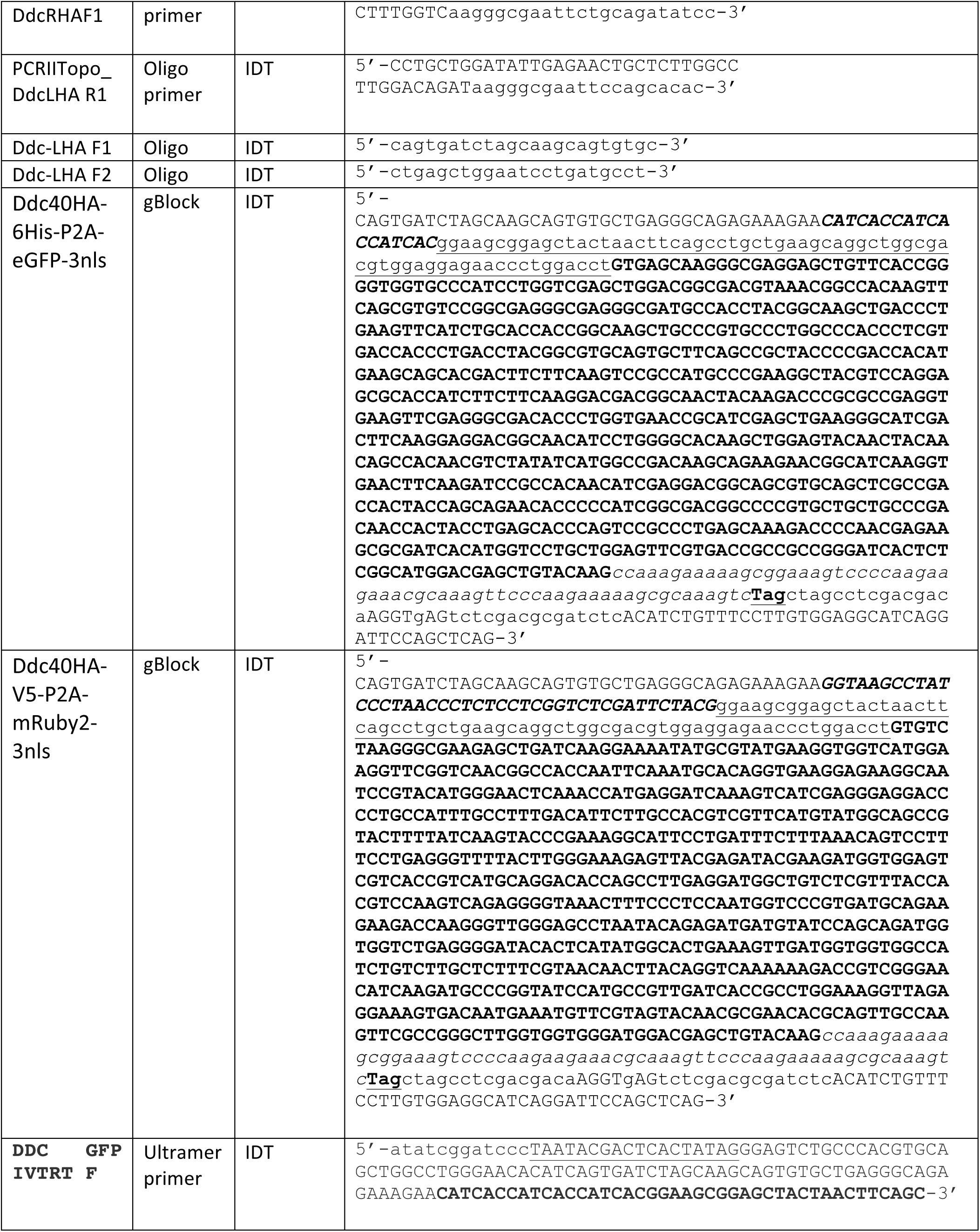
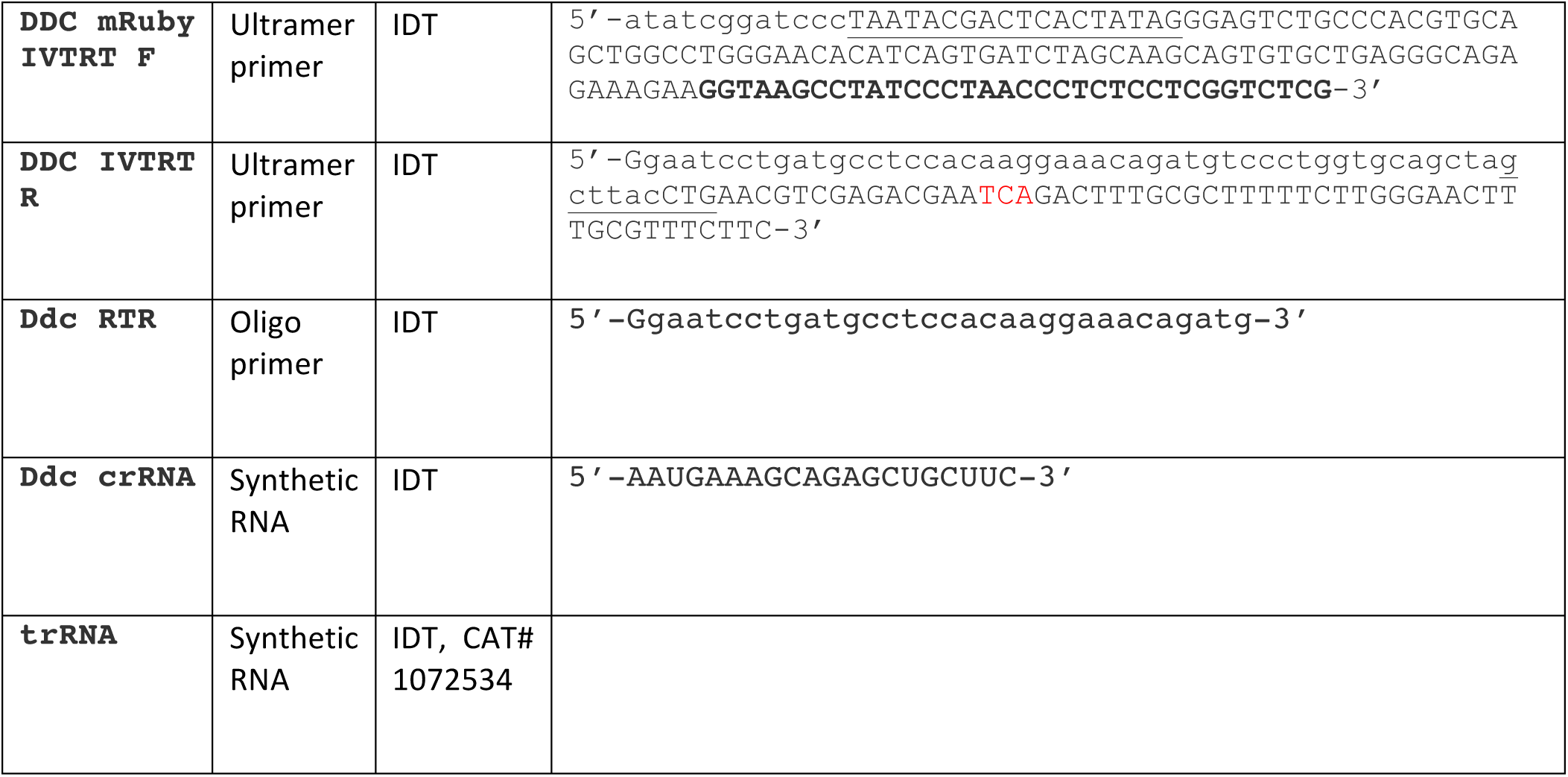
Synthetic DNA and RNAs.

