## Supplemental Data for "Dopa decarboxylase is a genetic hub of parental control over offspring behavior"

### Bulk RNA-Seq Imprinting~Expression Network Analysis

Here, we expand upon the details and controls involved in the Imprinting~Expression network analysis to define major cell classes with imprinting effects for different imprinted genes. In a control study, we computed a gene co-expression correlation matrix for the marker genes to test whether the data behave as expected. For the control study, we found the expected cell-type dependent gene co-expression relationships. For example, the conserved neuron marker gene, *Ahi1*, is correlated to another neuron marker gene, *Syn2*, but not to the oligodendrocyte marker, *Mopb*, or the astrocyte marker, *Sox9* (**Data S1A**). This finding generalized to all marker genes, such that marker genes for a given cell-type are more highly correlated to other markers of that cell type than to markers of other cell types (**Data S1B**). Thus, our datasets behave as expected and we turned to the analysis of imprinting ~ marker networks.

To begin we focused on two imprinted genes that are known to exhibit imprinting in neurons and not glia. *Ube3a* is a MEG known that preferentially expresses the *maternal* allele in neurons and not glia (Albrecht et al., 1997; Judson et al., 2014; Sato and Stryker, 2010; Yamasaki et al., 2003). Consistent with this, we observed a positive correlation between the *Ube3a* imprinting effect (maternal-paternal allele expression difference) and expression of the neuron marker, *Ahi1* (**Data S1C**). Peg3 is a PEG that expresses the *paternal* allele in neurons (He and Kim, 2014), thus we expect an inverse correlation between the maternal-paternal allele expression difference and *Ahi1* expression, which is observed (**Data S1D**). These results show how maternal and paternal imprinting effects correlate to the expression of cell-type marker genes in our approach. To include all 1000 marker genes for each cell-type in our analysis (**Data S1E**) and perform statistical testing for cell-type dependency of imprinting effects, we computed imprinting ~ marker correlation matrices for each MEG and PEG and then tallied the number of marker genes that are positively (rho > 0) or negatively (rho < 0) correlated to the imprinting effect (**Data S1F**). A Chi-square test of independence is then performed to ascertain whether the imprinting effect is dependent on cell type and the Pearson residuals reveal how maternal and paternal allele expression relates to specific cell types. For example, this analysis correctly shows *Ube3a* imprinting depends significantly on neuronal versus non-neuronal cell types and the maternal allele is positively associated with neurons (**Data S1G**). In contrast, the paternal allele for PEG3 is significantly associated with neurons (**Data S1H**), which is the expected result.

**Extended Behavior Analysis: *Ddc*-mediated imprinting and cross effects impact multiple dimensions of behavior**

Our deep analysis of foraging uncovered *Ddc*-mediated imprinting and cross effects on behavior and revealed that, in some cases, our statistical models must first absorb variance related to the parental cross effect before testing for potential maternal or paternal allele effects in offspring. We therefore carried out additional studies to: (1) Determine whether cross effects also impact offspring social behaviors, (2) Determine whether loss of the maternal *Ddc* allele affects female social behavior after absorbing variance related to the cross, and (3) Determine whether *Ddc*-mediated imprinting and cross effects can be further confirmed in lab tests of exploratory behaviors, including the light-dark box, elevated zero maze and open field tests.

To test for cross effects on sociability and social novelty seeking behaviors, we again used nested generalized linear models that first test the effect of the cross and then whether inheritance of a mutant paternal or maternal allele causes significant effects compared to $Ddc^{+/+}$ littermate controls. The results reveal significant cross effects on female (**Data S2A and B**) and male (**Data S2C**) sociability. Independent of the genotype of the offspring, we found females derived from the cross with a mutant mother spend more cumulative time in the conspecific chamber compared to the empty chamber, while females from the reciprocal cross do not (**Data S2A**). Additionally, latency to sniff the jail containing the conspecific is significantly reduced in $Ddc^{-/+}$ and $Ddc^{+/+}$ females from the maternal mutant cross (**Data S2B**). *Ddc*$^{-/+}$ and *Ddc*$^{+/+}$ male offspring from mutant mothers spent significantly more cumulative time in the zone near the jail with the conspecific compared to males from the reciprocal cross (**Data S2C**). Thus, cross effects shape female and male offspring social behaviors.

Our initial analyses showed no significant imprinting effects on female social behaviors. However, after absorbing variance due to the cross effect, we found a significant main effect of the maternal allele genotype and that *Ddc*$^{-/+}$ females travel an increased distance in the sociability test (**Data S2D**) and spend significantly less time in the Conspecific mouse chamber per visit (**Data S2E**). In the social novelty test, we also found a significant maternal allele genotype effect on the distance traveled and that *Ddc*$^{-/+}$ females travel farther (**Data S2F**), and spend less time in the Stranger mouse chamber per visit (**Data S3G**). In testing the effects of losing the paternal *Ddc* allele, we found a significant effect the frequency of sniffing the Stranger Mouse jail in the social novelty test and that on *Ddc*$^{+/-}$ females sniff the jail more frequently than controls (**Data S2H**). Therefore, after accounting for cross effects, the genotype of the maternal *Ddc* allele affects female social behaviors differently from loss of the paternal allele. Taken with our initial results in males (**Fig. 4**), we conclude that *Ddc* imprinting shapes male and female social behaviors.

Finally, to further investigate *Ddc* imprinting and cross effects, we performed lab tests of exploratory and anxiety-like behaviors, including open field, elevated plus maze, and light-dark box tests. Significant cross effects were found in males in the open field test and, similar to our foraging test, no significant cross effects were identified in females (**Data S3A, B**). Males from a mother with a mutant *Ddc* allele had fewer visits to the center zone (**Data S3A**) and spent less time moving during the test (**Data S3B**). We did not observe significant imprinting effects in the open field test. However, after absorbing variance due to the cross, we uncovered significant imprinting effects in daughters in the light-dark box test, such that loss of the maternal *Ddc* allele caused significant behavioral changes involving an increased number of transitions from the light to dark sides (**Data S3C**). In contrast, loss of the paternal allele resulted in decreased total time in the light side (**Data S3D**). We did not observe significant cross or imprinting effects on male or female body weights (**Data S3E**). These added behavioral tests further support our major conclusion that *Ddc* mediates multiple different parental effects on offspring behavior.

Data S1

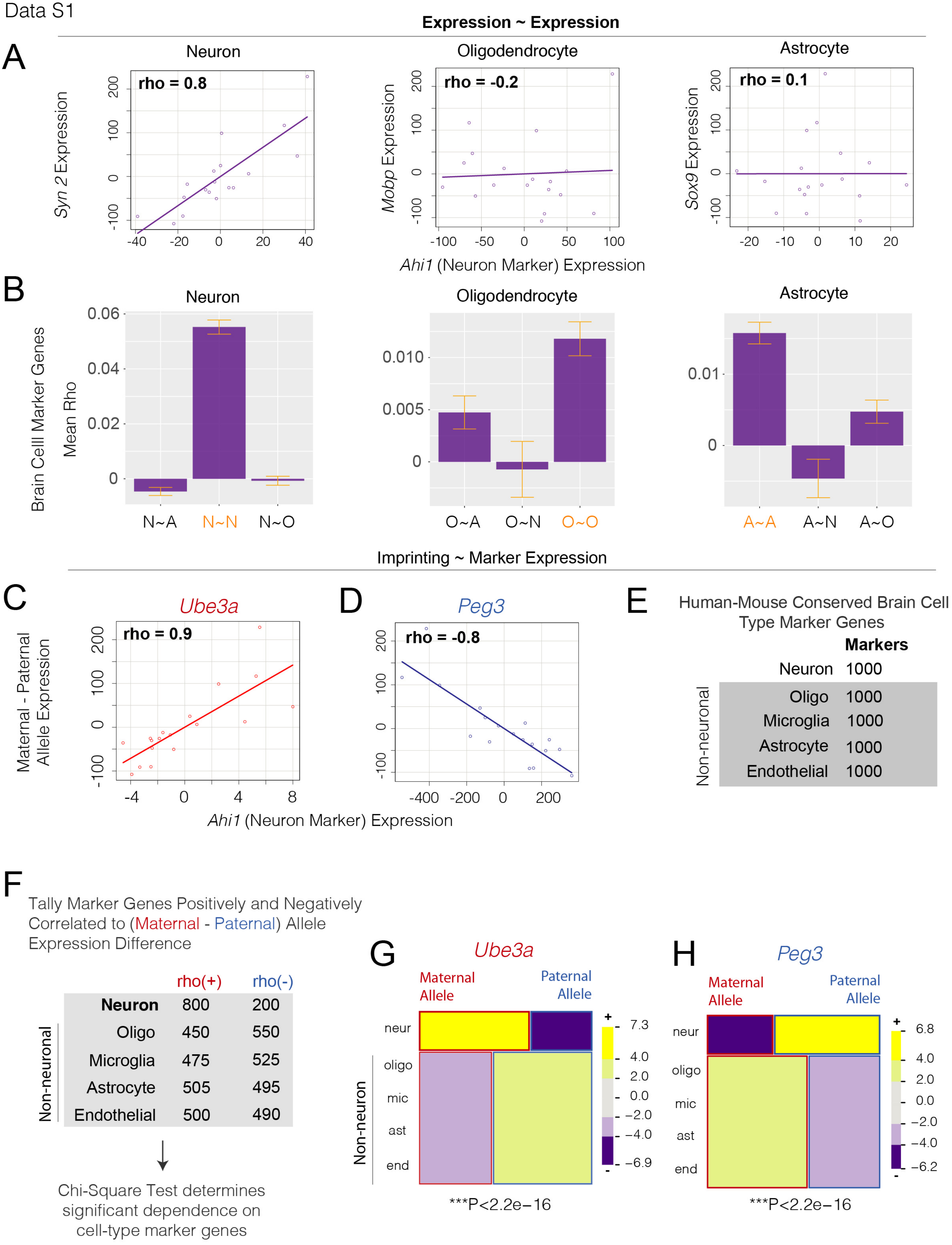

Expression ~ Expression
A
Neuron
Oligodendrocyte
Astrocyte
rho = 0.8
rho = -0.2
rho = 0.1
Syn 2 Expression
Mobp Expression
Sox9 Expression
Ahi1 (Neuron Marker) Expression
B
Brain Celll Marker Genes
Mean Rho
N~A
N~N
N~O
O~A
O~N
O~O
A~A
A~N
A~O
Imprinting ~ Marker Expression
C
Ube3a
rho = 0.9
Maternal - Paternal Allele Expression
D
Peg3
rho = -0.8
Ahi1 (Neuron Marker) Expression
E
Human-Mouse Conserved Brain Cell Type Marker Genes
Markers
Neuron 1000
Non-neuronal
Oligo 1000
Microglia 1000
Astrocyte 1000
Endothelial 1000
F
Tally Marker Genes Positively and Negatively Correlated to (Maternal - Paternal) Allele Expression Difference
rho(+) rho(-)
Neuron 800 200
Non-neuronal
Oligo 450 550
Microglia 475 525
Astrocyte 505 495
Endothelial 500 490
Chi-Square Test determines significant dependence on cell-type marker genes
G
Ube3a
Maternal Allele
Paternal Allele
neur
oligo
mic
ast
end
Non-neuron
7.3
4.0
2.0
0.0
−2.0
−4.0
−6.9
***P<2.2e−16
H
Peg3
Maternal Allele
Paternal Allele
neur
oligo
mic
ast
end
6.8
4.0
2.0
0.0
−2.0
−4.0
−6.2
***P<2.2e−16

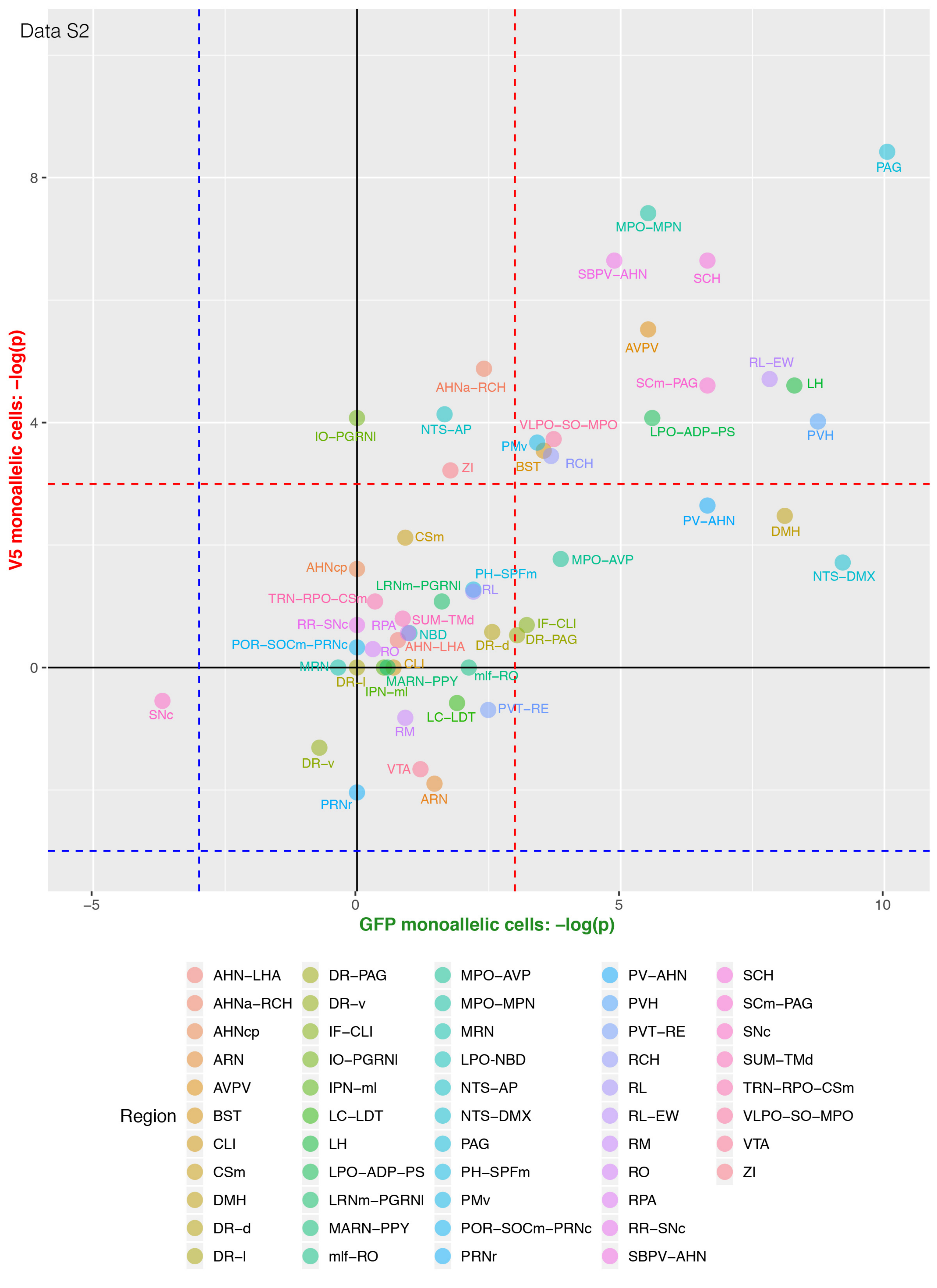

Data S2
V5 monoallelic cells: −log(p)
GFP monoallelic cells: −log(p)
8
4
0
−5
0
5
10
PAG
MPO−MPN
SBPV−AHN
SCH
AVPV
RL−EW
LH
SCm−PAG
AHNa−RCH
IO−PGRNI
NTS−AP
VLPO−SO−MPO
LPO−ADP−PS
PVH
PMv
BST
RCH
ZI
PV−AHN
DMH
CSm
MPO−AVP
NTS−DMX
AHNcp
PH−SPFm
LRNm−PGRNI
RL
TRN−RPO−CSm
RR−SNc
RPA
SUM−TMd
IF−CLI
NBD
DR−PAG
POR−SOCm−PRNc
RO
AHN−LHA
DR−d
MRN
CLI
DR−l
MARN−PPY
mlf−RO
IPN−ml
SNc
PVT−RE
LC−LDT
RM
DR−v
VTA
ARN
PRNr
Region
AHN−LHA
AHNa−RCH
AHNcp
ARN
AVPV
BST
CLI
CSm
DMH
DR−d
DR−l
DR−PAG
DR−v
IF−CLI
IO−PGRNI
IPN−ml
LC−LDT
LH
LPO−ADP−PS
LRNm−PGRNI
MARN−PPY
mlf−RO
MPO−AVP
MPO−MPN
MRN
LPO-NBD
NTS−AP
NTS−DMX
PAG
PH−SPFm
PMv
POR−SOCm−PRNc
PRNr
PV−AHN
PVH
PVT−RE
RCH
RL
RL−EW
RM
RO
RPA
RR−SNc
SBPV−AHN
SCH
SCm−PAG
SNc
SUM−TMd
TRN−RPO−CSm
VLPO−SO−MPO
VTA
ZI

Data S3

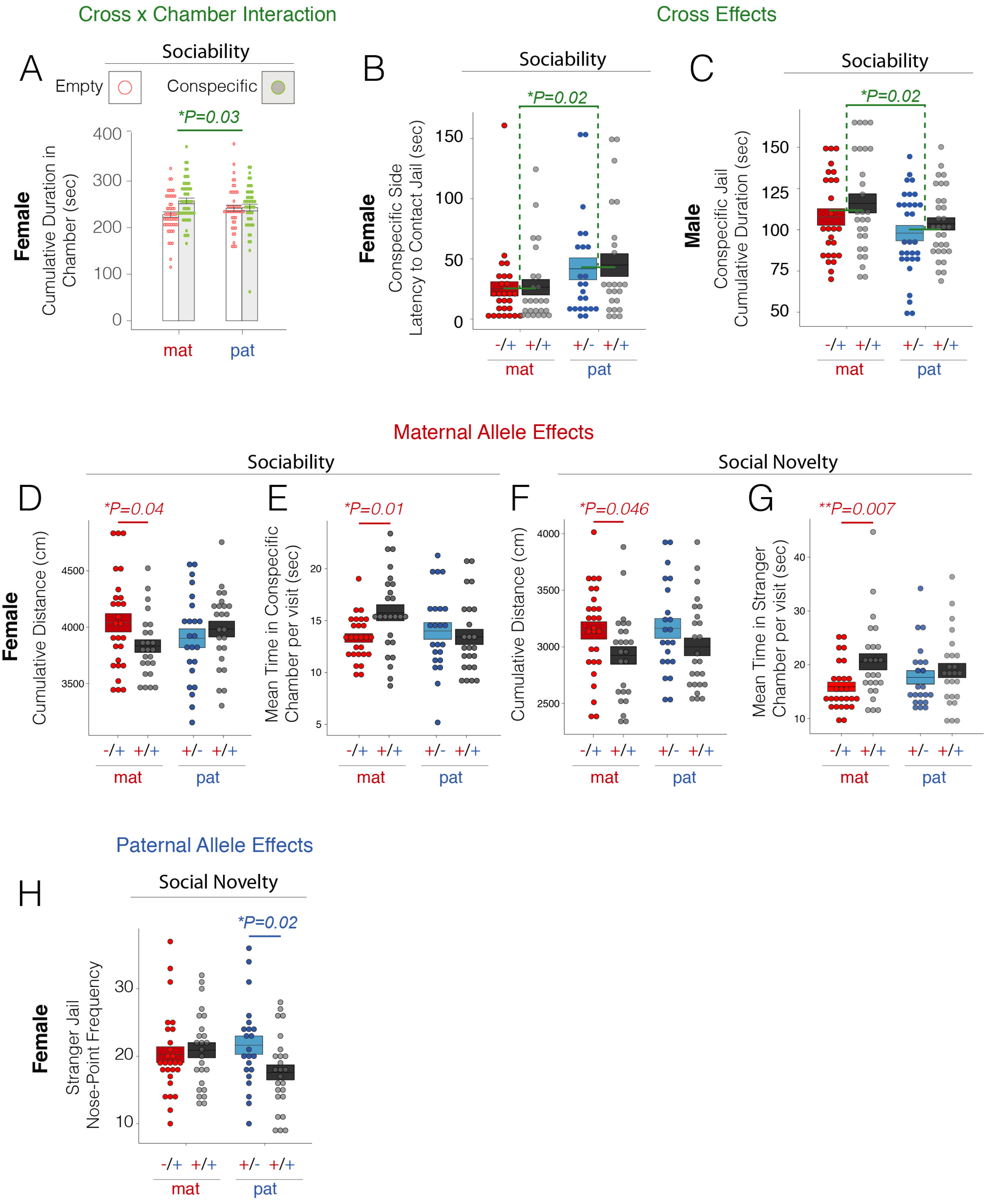
Cross x Chamber Interaction
Cross Effects
A
Sociability
Empty
Conspecific
*P=0.03
Female
Cumulative Duration in Chamber (sec)
400
300
200
100
0
mat
pat
B
Sociability
*P=0.02
Female
Conspecific Side Latency to Contact Jail (sec)
150
100
50
0
-/+ +/+ +/- +/+
mat
pat
C
Sociability
*P=0.02
Male
Conspecific Jail Cumulative Duration (sec)
150
125
100
75
50
-/+ +/+ +/- +/+
mat
pat
Maternal Allele Effects
Sociability
Social Novelty
D
*P=0.04
Female
Cumulative Distance (cm)
4500
4000
3500
E
*P=0.01
Mean Time in Conspecific Chamber per visit (sec)
20
15
10
5
F
*P=0.046
Cumulative Distance (cm)
4000
3500
3000
2500
G
**P=0.007
Mean Time in Stranger Chamber per visit (sec)
40
30
20
10
-/+ +/+ +/- +/+
mat
pat
Paternal Allele Effects
H
Social Novelty
*P=0.02
Female
Stranger Jail Nose-Point Frequency
30
20
10
-/+ +/+ +/- +/+
mat
pat

Data S4

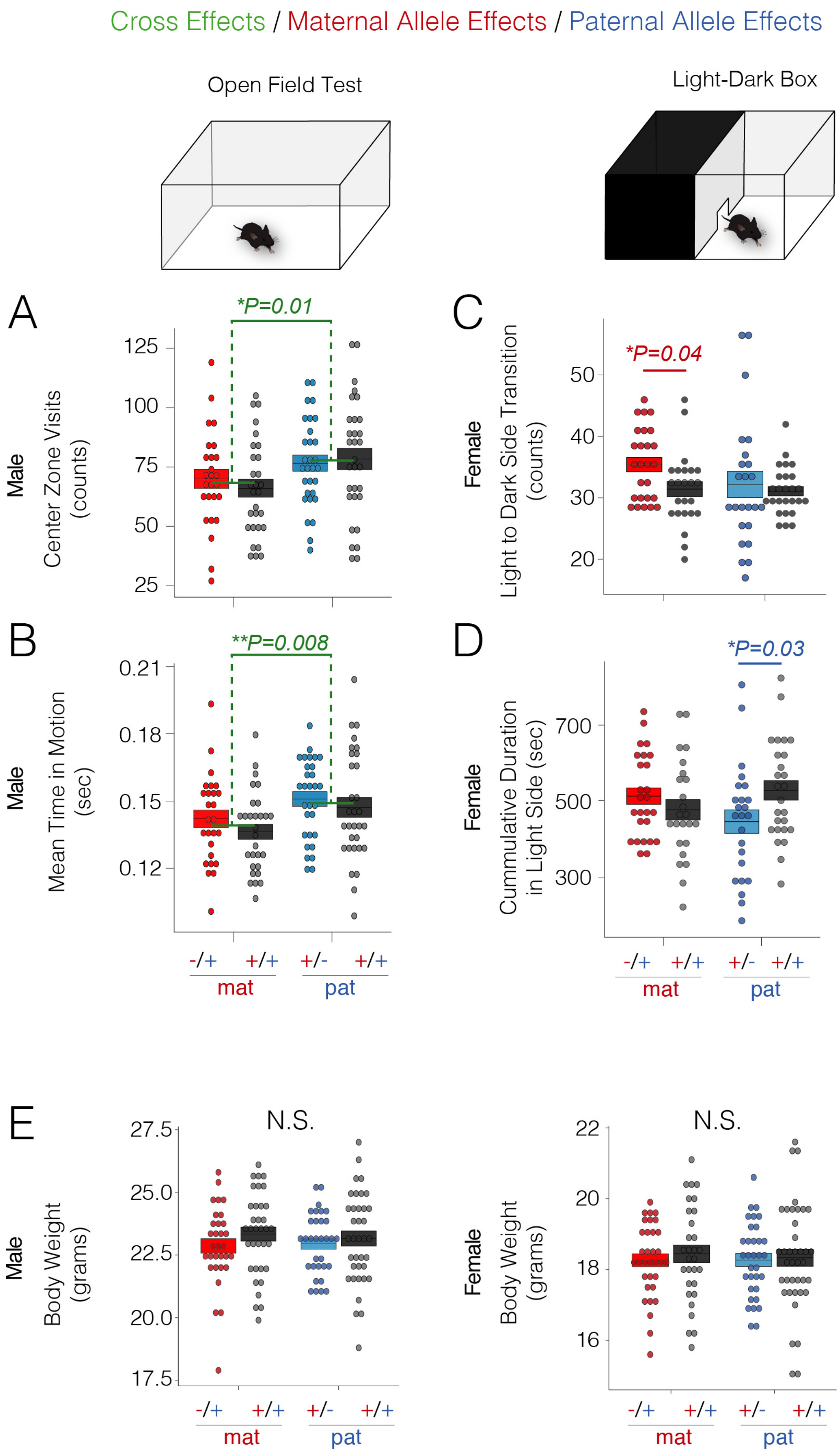

Cross Effects / Maternal Allele Effects / Paternal Allele Effects
Open Field Test
Light-Dark Box
A
*P=0.01
Male
Center Zone Visits (counts)
125
100
75
50
25
C
*P=0.04
Female
Light to Dark Side Transition (counts)
50
40
30
20
B
**P=0.008
Male
Mean Time in Motion (sec)
0.21
0.18
0.15
0.12
D
*P=0.03
Female
Cummulative Duration in Light Side (sec)
700
500
300
-/+ +/+ +/- +/+
mat pat
E
N.S.
Male
Body Weight (grams)
27.5
25.0
22.5
20.0
17.5
Female
Body Weight (grams)
22
20
18
16
